# A miniature bioluminescent platform for imaging structured RNAs

**DOI:** 10.64898/2026.10.01.755752

**Authors:** Christelle E. T. Chan, Erin B. Fuller, Claire C. Chen, Omeed Rabie, Nicholas T. Seto, Oswald Steward, Jennifer A. Prescher, Andrej Lupták

## Abstract

A broader understanding of RNA biology requires methods for visualizing transcript dynamics in native environments. Common imaging approaches rely on fluorescent probes that require excitation light and often depend on large aptamer arrays, limiting sensitivity and compatibility with many non-coding RNAs, particularly highly structured folds. To address these challenges, we developed a miniaturized bioluminescent platform for RNA imaging. This system employs a compact RNA tag that assembles luciferase fragments upon binding of monomeric RNA-binding proteins (RNA lanterns). Systematic optimization of the RNA component yielded a structured 24-nucleotide tag, representing greater than 3-fold reduction in size relative to the original *C-11-B* RNA tag, placing it among the smallest RNA imaging tags reported to date. The optimized tag enabled RNA-dependent bioluminescence detection in live cells, notably as a small, single insert within highly structured non-coding RNAs, including the *MALAT1 ENE* domain and a small HDV-like ribozyme. Collectively, this work establishes a miniaturized bioluminescent platform for imaging a broad range of RNA targets in living systems.

## INTRODUCTION

RNAs are synthesized through tightly regulated pathways and, once mature, execute and coordinate a wide range of cellular processes(Sharp 2009). Diverse classes of non-protein-coding RNAs (ncRNAs), including long non-coding RNAs (lncRNAs) and small non-coding RNAs (sncRNAs), regulate transcription, RNA processing, localization, and translation(Cech and Steitz 2014; Morris and Mattick 2014; Kung et al. 2013). These activities are intrinsically dynamic and spatially organized, because many transcripts are actively trafficked to defined subcellular compartments where localization dictates function(Martin and Ephrussi 2009; Buxbaum et al. 2015). Disruption of these processes contributes to developmental abnormalities and disease, and numerous RNAs have emerged as biomarkers and therapeutic targets(Loganathan and Doss C 2023; Gutschner et al. 2013). Yet, for many transcripts, the relationship between localization, dynamics, and biological function remain unclear, and additional RNA-mediated cellular mechanisms remain to be discovered. Despite their central regulatory roles, RNA dynamics in living systems remain difficult to study.

A deeper understanding of RNA biology requires methods that capture transcript behavior across the full RNA life cycle, from synthesis to decay, in their native cellular environment, ideally traced throughout the lifetime of a cell or a whole animal. Conventional RNA imaging approaches often rely on fluorescent optical probes(Brodsky and Silver 2002; Wu et al. 2014; Larson et al. 2011), with a widely used strategy employing the bacteriophage MS2 and PP7 coat proteins (MCP and PCP, respectively) fused to a split fluorescent reporter and assembled upon binding MS2 and PP7 RNA aptamer tags (Wu et al. 2014; Bertrand et al. 1998). Although this method has enabled studies of RNA localization, these reporters have a notable limitation: achieving sufficient signal-to-noise typically requires tandem arrays of aptamer tags, often spanning hundreds of nucleotides (nts) which can perturb RNA structure, localization, and downstream function(Wu et al. 2014; Larson et al. 2011; Gerber et al. 2023). Efforts to reduce RNA tag size while maintaining efficient photon emission have led to the development of fluorogenic RNA aptamers, which generate fluorescence upon binding small-molecule dyes(Paige et al. 2011; Filonov et al. 2014; Dolgosheina et al. 2014). Recent variants, such as *i*Mango-III, exhibit brightness exceeding enhanced green fluorescent protein (eGFP), and advances in fluorescence lifetime imaging have further enabled imaging with a single fluorogenic aptamer tag(Trachman et al. 2019; Sarfraz et al. 2023). However, both probe-based systems and fluorogenic aptamer-dye systems require external excitation light, which can introduce cellular toxicity and induce autofluorescence(Icha et al. 2017). In addition, fluorescent proteins are prone to photobleaching during prolonged excitation, whereas fluorogenic aptamers often exhibit limited photostability, necessitating pulsed irradiation and restricting their use in prolonged imaging applications(Trachman et al. 2019; Tung et al. 2016; Autour et al. 2018).

Bioluminescence-based approaches offer an attractive alternative because light production does not require external excitation. Instead, light is emitted through luciferase-catalyzed oxygenation of a luciferin substrate, resulting in low background and high sensitivity(Tung et al. 2016). Despite these advantages, few bioluminescence-based strategies for RNA imaging have been established(Eguchi et al. 2023; Jiang et al. 2023; Mi et al. 2023; Mitkas et al. 2022). To fill in this gap, we and others have developed RNA imaging platforms that couple RNA recognition to bioluminescent signal generation(Eguchi et al. 2023; Halbers et al. 2024; Fuller et al. 2026; Hall et al. 2012; Dixon et al. 2016). Among these systems, we recently developed a bioluminescent RNA imaging platform in which transcripts are fused with a single copy of an RNA aptamer tag that recruits specific RNA-binding proteins (RBPs). Each RBP is fused to a fragment of split NanoLuc luciferase (NanoBiT), such that binding to a specific RNA tag brings the fragments into proximity, reconstituting a functional light-emitting enzyme(Halbers et al. 2024; Fuller et al. 2026; Hall et al. 2012; Dixon et al. 2016). These luciferase fusions, termed RNA lanterns, produce photons upon RNA-dependent complementation and enable continuous imaging over many hours without detriment(Halbers et al. 2024; Fuller et al. 2026).

Early RNA lanterns featuring the MCP and PCP were used to recruit NanoBiT fragments to an engineered MS2 and PP7 aptamer tag appended to transcripts of interest(Halbers et al. 2024). This approach enabled low-background imaging of model transcripts in live cells and *in vivo* using a single copy of the tag without external excitation(Halbers et al. 2024). Orthogonal bioluminescence resonance energy transfer (BRET) lantern variants were also developed using MS2 and either PP7, CS2, and boxB aptamers to red-shift the emission and facilitate multiplex imaging of transcripts(Fuller et al. 2026; Bernardi and Spahr 1972; Lim et al. 2001; Faber et al. 2001; Chattopadhyay et al. 1995; Kuhn et al. 2002; Omer et al. 2002; Fukunaga and Yokobayashi 2022). However, because these lanterns rely on MCP and PCP, they retain a substantial footprint associated with multimeric coat proteins and may promote suboptimal stoichiometries for NanoBiT complementation(Bernardi and Spahr 1972; Lim et al. 2001; Valegård et al. 1997; Chao et al. 2008). These features limit compatibility with structured or functionally sensitive transcripts and highlight the need for more compact imaging strategies.

To address these limitations, a second-generation RNA lantern system was developed using orthogonal RBPs L7Ae and Nun, which bind to their respective RNA motifs (CS2 and boxB) as monomers with high binding affinities(Faber et al. 2001; Chattopadhyay et al. 1995; Kuhn et al. 2002; Omer et al. 2002; Fukunaga and Yokobayashi 2022). In this configuration, each RBP is fused to a NanoBiT fragment, promoting a one-to-one recruitment of LgBiT and SmBiT to the target RNA and resulting in improved lantern assembly. The resulting lanterns were further optimized for enhanced BRET, producing more red-shifted emission while maintaining a smaller probe size(Fuller et al. Accepted).

Here, we build on this framework to develop a miniaturized bioluminescent RNA imaging platform that reduces the size of the RNA tag while maintaining efficient lantern assembly. The approach uses the monomeric L7Ae-Nun lantern in combination with compact CS2-boxB RNA tags that were rationally designed and systematically optimized for luminescence activation. The complete imaging system can be implemented via a single-vector co-expression strategy, enabling coordinated delivery and RNA detection across different cell types. The minimized tags enable RNA imaging in live cells and are compatible with highly structured RNAs. We demonstrate successful incorporation of the minimized tag into two tightly folded RNAs, the *MALAT1 ENE* and an HDV-like self-cleaving ribozyme, without disrupting function. Together, this work establishes a miniaturized platform for imaging RNA in physiological contexts, expanding the scope of transcripts accessible to live-cell RNA imaging while preserving structural and functional integrity.

## RESULTS

### Minimization and biochemical evaluation of the RNA tag

We previously developed a series of structured RNA tags composed of the CS2 and boxB aptamers separated by variable spacer lengths to modulate their relative distance and orientation (*C-X-B*, where *X* denotes the number of nucleotides between aptamers)(Fuller et al. Accepted). These tags enabled the assembly of the L7Ae-Nun lantern, in which the RNA-binding protein L7Ae recognizes the kink-turn (k-turn) of an *in vitro* evolved CS2 aptamer, and Nun recognizes the boxB aptamer(Faber et al. 2001; Chattopadhyay et al. 1995; Kuhn et al. 2002; Omer et al. 2002; Fukunaga and Yokobayashi 2022). These tags embodied a four-way junction scaffold consisting of two aptamers for RNA lantern assembly and an established Ni-NTA-binding aptamer incorporated for tag rigidification and biochemical validation of RNA lantern complex formation(Halbers et al. 2024; Fuller et al. 2026; Chizzolini et al. 2020). To minimize this platform, we sought to reduce the size of the *C-X-B* tags while maintaining the ability to assemble the monomeric L7Ae-Nun lantern. In the minimized design, the Ni-NTA aptamer was removed and the CS2 and boxB aptamers were directly fused through a short helix. This design leverages the intrinsic kink-turn structure of CS2 and replaces its P2 stem-loop with the boxB hairpin. We generated a putative panel of minimized boxB-CS2 tags (*mB-X-C*), in which the intervening base pairs (denoted *X*) modulate aptamer spacing and orientation (**Fig. 1A**). Secondary structure modeling using ViennaRNA and Forna were used to inform minimized tag design and predicted stability(Lorenz et al. 2011; Kerpedjiev et al. 2015). The resulting panel (*mB-1-C* through *mB-15-C)*, sampled at least one full helical turn (∼360°) of phase and about 3.5 nm of distance between the two aptamers (**Supplementary Fig. 1A**).

**Figure 1.**
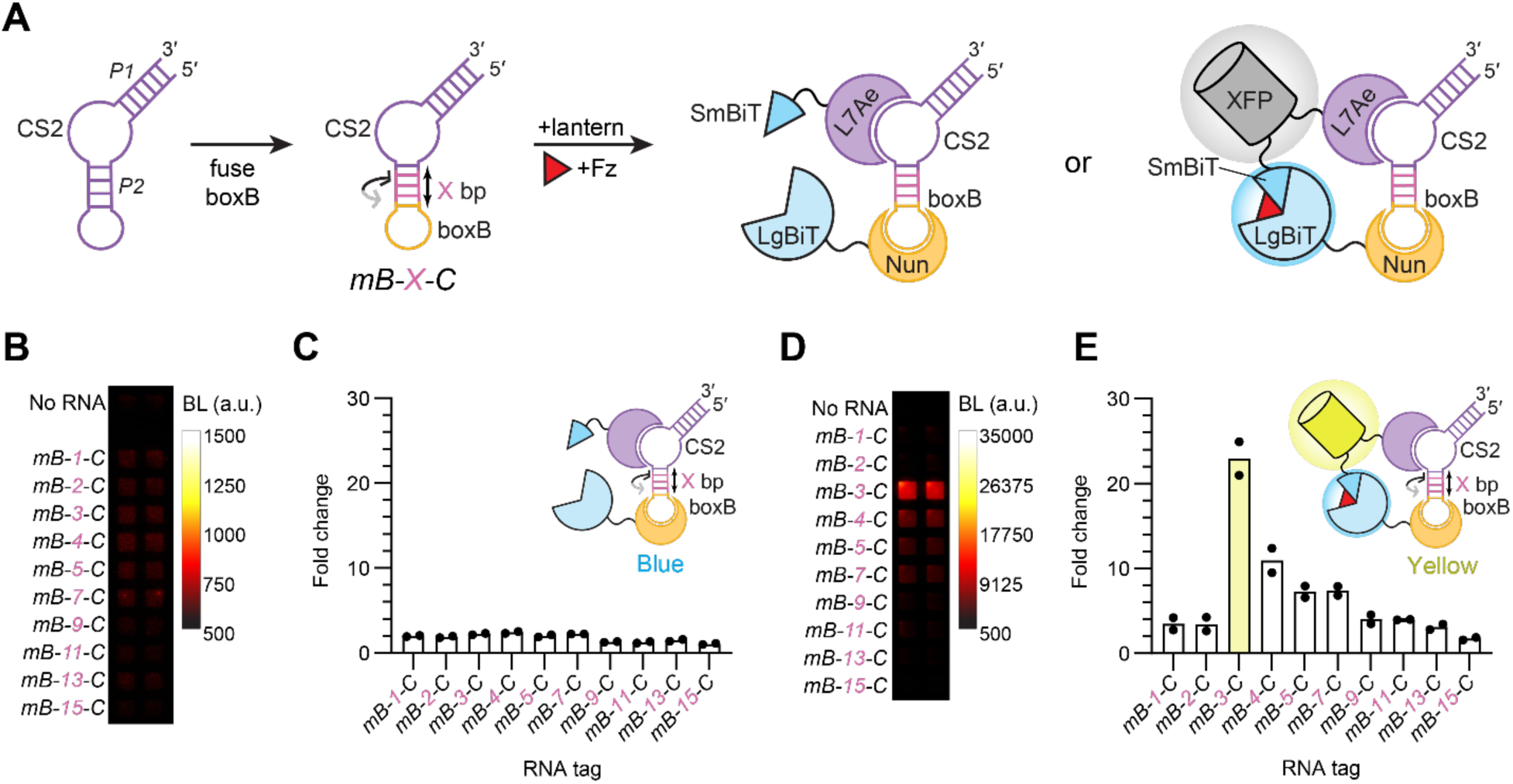
Minimization and initial evaluation of *boxB-CS2* RNA tags. (**A**) Schematic representation of the design strategy for minimized tags. The P2 stem-loop of the CS2 aptamer was replaced with the boxB aptamer to generate a panel of minimized *boxB-CS2* tags (*mB-X-C*), where *X* represents the number of base-pairs separating the two aptamers. Binding of L7Ae and Nun recruits the NanoBiT fragments (SmBiT and LgBiT) of the lantern into proximity. In the blue lantern configuration, linker lengths between NanoBiT fragments are insufficient to support efficient heterodimerization when bound to the minimized tag (left complex). In contrast, incorporation of a fluorescent protein (XFP) increases the effective spacing between NanoBiT fragments, enabling more efficient complementation (right complex). In the presence of furimazine (Fz), successful lantern assembly results in photon production. (**B**) Luminescence output from the panel of *mB-X-C* constructs combined with lysate from cells transiently expressing the blue variant of the L7Ae-Nun lantern. (**C**) Fold change in luminescence signal over no-RNA controls with blue L7Ae-Nun lantern. (**D**) Luminescence output from the same *mB-X-C* panel incubated with lysate from cells transiently expressing the yellow BRET variant (VenusΔC12) of L7Ae-Nun lantern (λ_max_ ∼530 nm, yellow glow). (**E**) Fold change in luminescence signal over no-RNA controls with yellow L7Ae-Nun lantern. (**C, E**) Varying spacer lengths, highlighted in pink, modulate the distance and rotational phase between the aptamers. Bars represent the mean of *n* = 2 replicates, with individual data points shown. BL, luminescence; a.u., arbitrary units.

As an initial test, the minimized RNA tags were transcribed *in vitro* and added exogenously to HEK293T cell lysate expressing the blue L7Ae-Nun lantern. Luminescence was measured across the minimized constructs to identify tags capable of activating the blue L7Ae-Nun lantern. Across the panel, the minimized tags produced only modest photon outputs about 1–2-fold over the no-RNA control, indicating inefficient lantern assembly (**Fig. 1B, C; Supplementary Fig. 2A**). Even so, these observations suggested that the lantern could be assembled with substantially smaller RNA tags, but that further optimization was required.

The same panel of tags was subsequently evaluated with the yellow variant (VenusΔC12) of the L7Ae-Nun BRET lantern under identical assay conditions. In this case, the *mB-3-C* tag (36 nts) exhibited the highest luminescent signal, yielding >20-fold signal enhancement relative to the no-RNA control. The improved performance of the yellow L7Ae-Nun BRET lantern is consistent with the increased effective distance between NanoBiT fragments introduced by the inclusion of the fluorescent protein. Extending the spacer length from *mB-3-C* to *mB-13-C*, corresponding to approximately one helical turn (∼360°) between boxB and CS2, failed to restore activity, suggesting that inefficient assembly cannot be rescued by altering rotational phase (**Fig. 1D, E; Supplementary Fig. 2B**). This behavior is likely due to a configuration in NanoBiT fragment orientation such that their complementation interfaces are directed away from one another, resulting in a strong preference for a single favorable orientation despite sampling different rotational alignments.

RNA-dependent lantern assembly was further validated *in vitro.* The *mB-3-C* tag was titrated in the same lysate containing the yellow L7Ae-Nun BRET lantern. Increasing *mB-3-C* concentrations (0–316 nM RNA) produced a corresponding increase in luminescence before reaching saturation (∼100 nM), consistent with RNA-mediated lantern assembly. At higher concentrations, the signal decreased, consistent with a “hook effect” arising from excess tag sequestering lantern components on separate RNAs. As anticipated, lantern formation was also limited at lower concentrations. In contrast, titration of an inactive variant, *mB-15-C,* failed to produce a measurable signal increase even at elevated RNA concentrations, demonstrating that lantern activation is dependent on both RNA presence and proper aptamer orientation (**Supplementary Fig. 3A–C**). RNA-dependent lantern assembly was further supported by pulldown of active RNA lantern complexes using the FLAG epitope tag engineered into a lantern component. (**Supplementary Fig. 4A–C**).

We next explored whether the *mB-3-C* scaffold could be further tuned to improve lantern assembly by introducing minor structural perturbations within the P2 helix connecting the two aptamers (**Supplementary Fig. 5A**). Insertion of a single-adenosine bulge at varying positions (*mB-3-C-b1* to *b3)* abolished luminescence in all cases, indicating that even minor distortions to the RNA tag structure disrupt NanoBiT complementation (**Supplementary Fig. 5B, C**). This result suggests that *mB-3-C* adopts a defined conformation that optimally positions the boxB and CS2 aptamers for simultaneous binding of L7Ae and Nun. We anticipate that such structural sensitivity may ultimately be advantageous for engineering RNA-based sensing devices.

### Biochemical optimization of the minimized RNA tag

Although lantern assembly is possible with a substantially reduced RNA tag, such as *mB-3-C*, further optimization of the tag design could improve lantern assembly efficiency and grafting into small structured RNAs, while maintaining compatibility with future multi-color applications (Fuller et al. 2026; Yao et al. 2022; Scipioni et al. 2026). We therefore revisited the tag design to further optimize aptamer arrangement. RNA tags can be rapidly iterated relative to protein-based lantern components, providing an efficient route to improve assembly efficiency and expand platform versatility.

To guide this effort, we performed structure-informed modeling using ChimeraX to evaluate alternative aptamer configurations and their potential to support lantern assembly(Goddard et al. 2018). Crude models of the *mB-X-C* and lantern suggested that the C-termini of L7Ae and Nun, which are fused to NanoBiT fragments, are oriented in opposite directions upon aptamer binding. Increasing spacer length would therefore further separate the luciferase fragments, likely contributing to the lack of signal recovery despite sampling a range of helical phases. These observations suggested that further RNA tag improvements would require a more substantial redesign rather than an adjustment of aptamer spacing.

We therefore explored a permuted tag configuration to reorient the bound lantern components, while preserving the compact nature of the RNA tag. In the original *mB-X-C* RNA tag, the boxB hairpin replaced the P2 stem-loop of the CS2 aptamer. In the revised tag, the boxB hairpin was repositioned to the P1 stem of the CS2. To accommodate this rearrangement, the P2 stem-loop was opened to generate new termini, yielding a distinct CS2-boxB configuration (*mC-X-B*). Although both tag configurations contain the same RNA recognition elements, this permutation alters the relative orientation of the bound L7Ae and Nun RBPs and, in turn, the positioning of the NanoBiT fragments (**Fig. 2A**). Crude modeling further supported that this rearrangement could improve the spatial orientation of the bound lantern components for efficient complementation while maintaining a highly compact RNA scaffold. Based on these insights, we generated a new panel of CS2-boxB tags (*mC-X-B),* in which the number of base-pairs separating the two aptamers (X) was systematically varied to explore spacing and helical phase (**Supplementary Fig. 6**).

**Figure 2.**
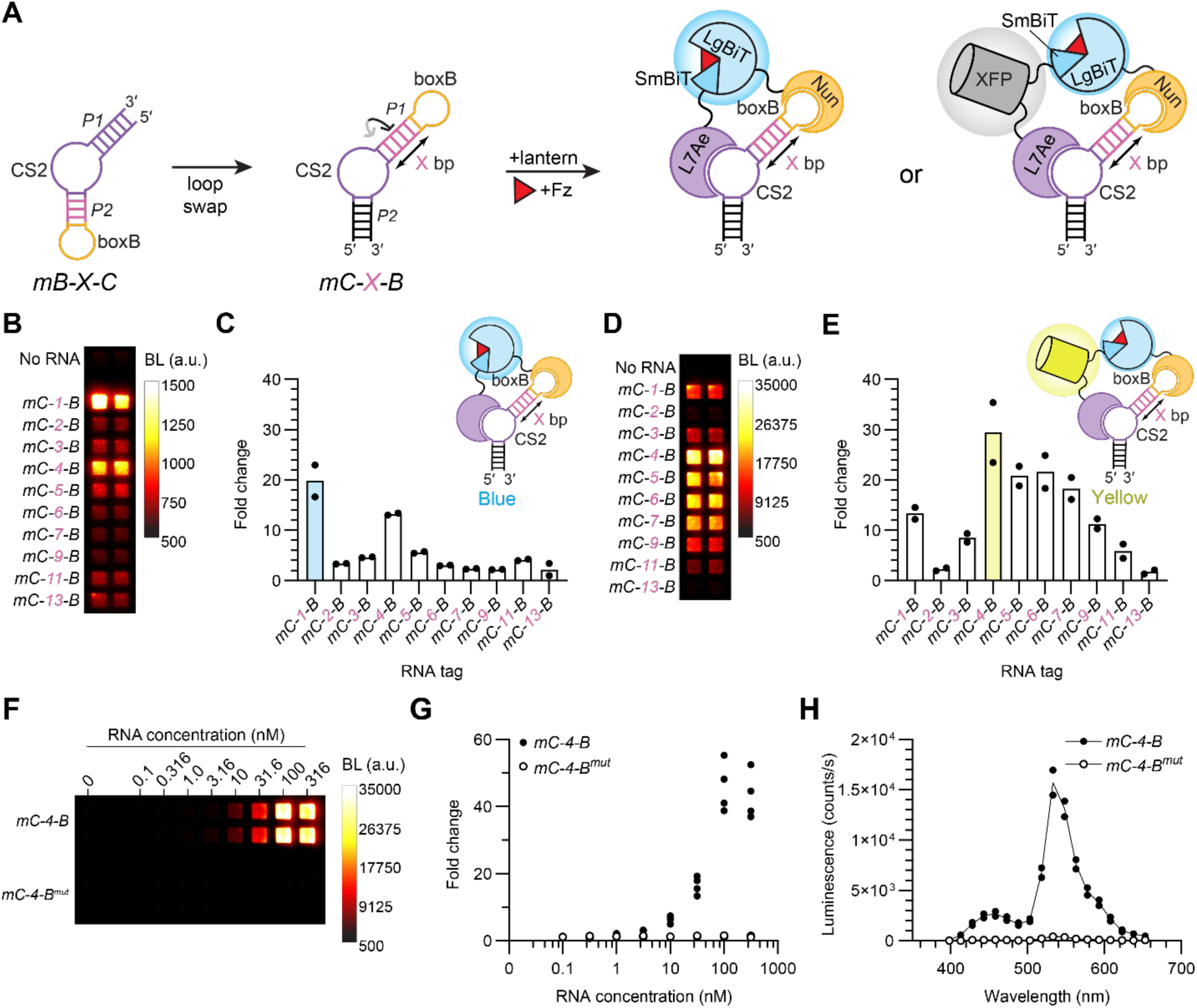
Minimized RNA tag optimization. (**A**) Schematic representation of minimized tag optimization. In the initial design (*mB-X-C*), the boxB hairpin was fused to the P2 stem of the CS2 aptamer (**Fig. 1**). Structure-informed modeling of the RNA tag and lantern structure suggested repositioning boxB to the P1 stem of CS2 to improve aptamer orientation for efficient NanoBiT complementation. This redesign generated a new panel of *CS2-boxB* tags (*mC-X-B)*, where X denotes the number of base pairs between the two aptamers. Binding of L7Ae and Nun recruits the NanoBiT fragments (SmBiT and LgBiT), enabling heterodimerization with or without a fluorescent protein present. In the presence of Fz, light is produced. (**B**) Luminescence output from the panel of *mC-X-B* constructs combined with lysate from cells transiently expressing the blue L7Ae-Nun lantern. (**C**) Fold change in luminescence signal relative to the no-RNA control for the blue L7Ae-Nun lantern. (**D**) Luminescence output from the same *mC-X-B* panel incubated with lysate from cells transiently expressing the yellow L7Ae-Nun-Venus lantern. (**E**) Fold change in luminescence signal over no-RNA controls with yellow L7Ae-Nun lantern. (**C, E**) Spacer lengths highlighted in pink modulate the distance and rotational phase between the aptamers. Bars represent the mean of *n* = 2 replicates, with individual data points shown. (**F**) Evaluation of minimized tag (*mC-4-B*) versus mutant (*mC-4-B^mut^*) titrated into lysate containing the yellow L7Ae-Nun-Venus lantern. (**G**) Fold change in luminescence signal relative to the no-RNA control. Individual data points (*n* = 4) are shown per condition. (**H**) Emission spectra of yellow L7Ae-Nun lantern complexes assembled with *mC-4-B* or the mutant control. RNA tags (100 nM) were added into HEK293T lysates expressing the yellow L7Ae-Nun-Venus lantern. Distinct BRET emission (λ_max_ ∼530 nm) was observed with *mC-4-B* but not with the inactive mutant. Spectra represent the mean of *n* = 2 replicates, with individual data points shown. BL, luminescence; a.u., arbitrary units.

When evaluated with the blue L7Ae-Nun lantern in HEK293T lysate, the *mC-1-B* tag (32 nts) produced the highest signal, yielding a 20-fold increase in luminescence over the no-RNA control. Luminescence declined sharply as spacer length increased, indicating that even modest spacer increases impair NanoBiT complementation. A partial recovery was observed with *mC-4-B*, consistent with a strong dependence on helical phase, where specific spacer lengths restore a more favorable rotational alignment between the binding domains (**Fig. 2B, C; Supplementary Fig. 7A**). Compared to the *mB-X-C* series, the *mC-X-B* tags exhibited greater sensitivity to changes in rotational phase with the blue lantern. These data suggest that efficient complementation requires a compact tag design that maintains close proximity between the two aptamers, while achieving appropriate rotational orientation.

The same *mC-X-B* tag panel was subsequently evaluated with the yellow L7Ae-Nun BRET lantern under identical conditions. In this case, the slightly larger *mC-4-B* tag (38 nts) produced the highest luminescence output, yielding a ∼30-fold signal enhancement relative to the no-RNA control (**Fig. 2D, E; Supplementary Fig. 7B**). This shift in optimal spacing likely reflects the increased bulk and effective distance between NanoBiT fragments introduced by the fluorescent protein fusion, requiring a longer RNA scaffold for efficient complementation. Further increases in spacer length reduced luminescence, consistent with the aptamers positioning the NanoBiT fragments too far apart for efficient reconstitution. However, less signal was observed due to changes in rotation, which can be attributed to the shorter effective distance between NanoBiT fragments.

To confirm that luminescence activation resulted from specific aptamer-protein interactions, we introduced a point mutation into the boxB hairpin of the *mC-4-B* tag (G19A) (**Supplementary Fig. 7C**), which was previously reported to reduce Nun binding activity to <0.5% of wild-type affinity(Cilley and Williamson 1997). Incorporation of this mutation into the *mC-4-B* construct (*mC-4-B^mut^*) attenuated luminescence, confirming that lantern assembly requires binding recognition of both aptamers (**Fig. 2F, G; Supplementary Fig. 7D**).

To assess the BRET efficiency of the system, we compared the emission spectra from lantern complexes assembled by *mC-4-B* to the mutant across a range of RNA concentrations (0 to 316 nM). Titration of the *mC-4-B* RNA resulted in concentration-dependent increase in luminescence, accompanied by a spectral shift in the emission profile from blue to yellow. This shift is consistent with increased BRET based on the RNA lantern complexes formed. In contrast, the mutant tag exhibited substantially lower BRET across concentrations similar to that of the no-RNA control. Low levels of non-specific complementation were observed under conditions with no-RNA tag (**Fig. 2H; Supplementary Fig. 7E, F**).

Because the NanoBiT systems can exhibit concentration-dependent background, we next examined the influence of lantern expression on signal-to-noise outputs. Titration of mC-4-B into into lysates containing higher higher concentrations of lantern components, prepared from lantern-expressing HEK293T cells (106 cells/mL), produced increased bioluminescence, with maximal signal observed at RNA tag concentrations between ∼100 and 316 nM. However, this condition was also accompanied by elevated background signal in the no-RNA control (0 nM), consistent with increased non-specific NanoBiT complementation at higher lantern concentrations. In contrast, lysates prepared from lower lantern expression levels (10^5^ cells/mL) exhibited less background complementation and improved overall signal-to-noise. Under these conditions, the optimal concentration shifted 10-fold lower, with maximal signal observed near 10 nM RNA. This shift is consistent with the 10-fold decrease in lantern expression, indicating that optimal assembly occurs when the lantern and tag components are present at comparable levels. Although total luminescence was lower in diluted lysate, the reduced background resulted in higher fold change relative to the no-RNA control (**Supplementary Fig. 8A, B**). These results demonstrate that both signal intensity and background luminescence are dependent on the relative stoichiometry of the RNA lantern and its cognate tag. By tuning lantern expression levels, experimental conditions can be adjusted to favor either maximal photon emission or improved RNA-dependent sensitivity and dynamic range.

Although *mC-4-B* provided optimal signal with the yellow L7Ae-Nun lantern, the smallest tag *mC-1-B* retained measurable luminescence. We therefore asked how much further could the RNA tag be minimized while still supporting lantern assembly. To define the lower size limit for functional RNA tags, we truncated the P1 helix of *mC-1-B* (**Supplementary Fig. 9A**). The P1 stem was decreased from 5 base pairs to 1 base pair in single-base-pair increments (*5-mC-1-B* through *1-mC-1-B*), excluding the base pair that closes the CS2 bulge. This yielded a minimal construct of 24 nts (*1-mC-1-B*). When evaluated with the yellow L7Ae-Nun lantern, these truncated constructs retained a similar luminescent signal comparable to the full-length P1 stem, indicating that progressive reduction of the P1 stem does not abolish lantern assembly. The retained signal suggests L7Ae binding stabilizes the kink-turn sufficiently to support NanoBiT complementation even with a minimal P1 stem. (**Supplementary Fig. 9B, C**). These data define the lower limit size for functional RNA tags, with *1-mC-1-B* representing the smallest tag capable of supporting lantern assembly. We anticipate that similar truncation of the P1 helix would support activity of other *mC-X-B* tags. For downstream analyses, the *mC-4-B* tag was paired with the yellow L7Ae-Nun BRET lantern, because this configuration yielded the highest signal, dynamic range, and benefits from red-shifted emission for improved tissue penetrance(Weissleder and Ntziachristos 2003).

### Live-cell imaging of minimized RNA tags

Having established that the minimized tags support L7Ae-Nun lantern assembly *in vitro,* we sought to determine whether these tags could support RNA imaging in live cells. The 38-nt *mC-4-B* tag and the boxB mutant were transcribed *in vitro* and transfected into HEK293T cells stably expressing the yellow L7Ae-Nun lantern. Following a 24-hour incubation to allow RNA uptake, the luciferin substrate was administered and luminescence was visualized. Robust signal was observed in cells receiving the *mC-4-B* tag, whereas cells transfected with the mutant produced only minimal emission comparable to background levels, demonstrating that the compact tag is sufficient to recruit lantern components and activate bioluminescence within the cellular environment (**Fig. 3A, B**). Because the lantern includes a fluorescent protein (VenusΔC12), fluorescence signal was used to gauge lantern expression independently of RNA-dependent luminescence. VenusΔC12 fluorescence was readily detected in cells expressing the lantern, providing a complementary readout to confirm probe expression and allowing luminescent signal to be interpreted in the context of lantern abundance (**Fig. 3B; Supplementary Fig.10**).

**Figure 3.**
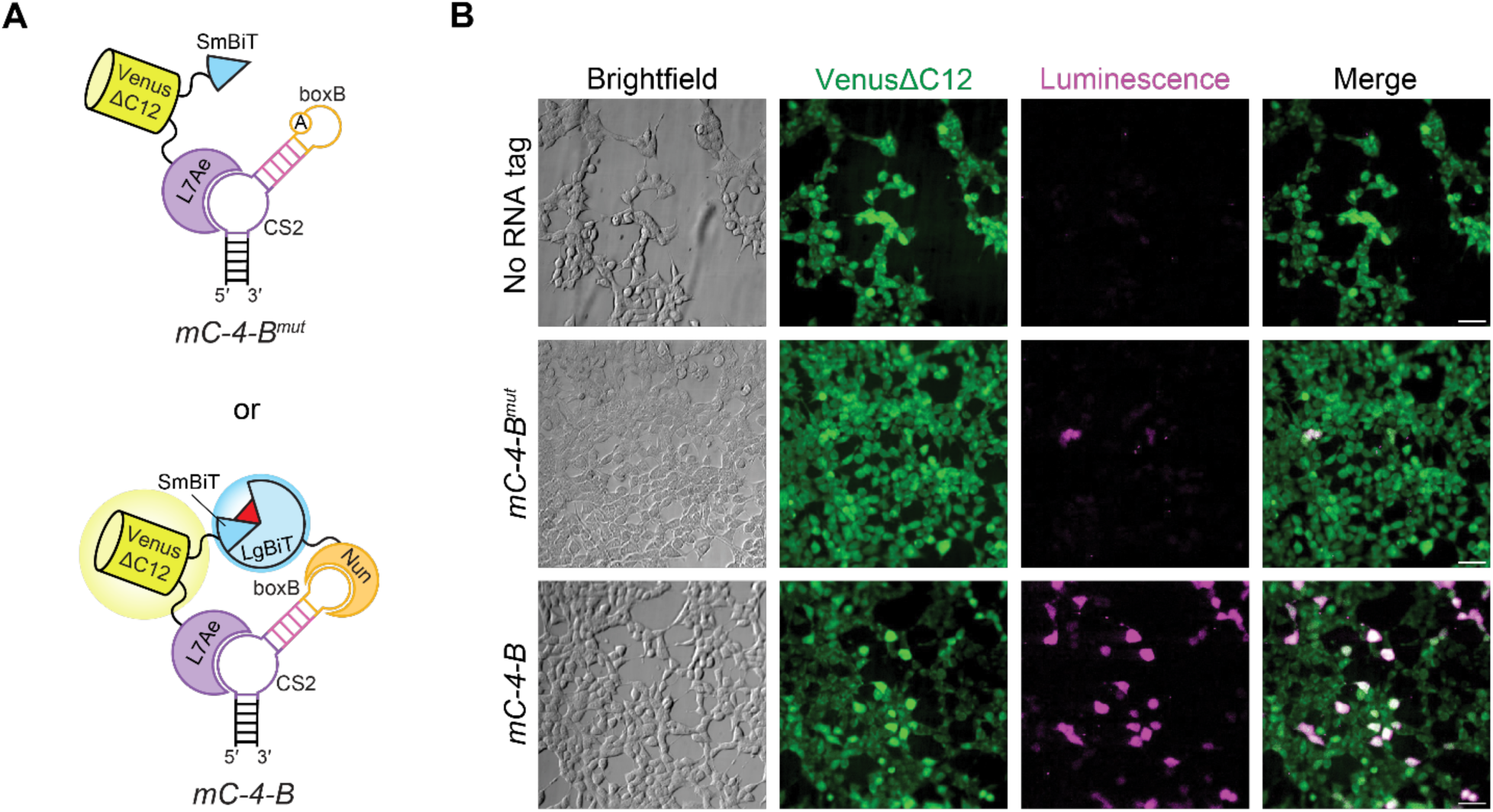
Bioluminescence of minimized RNA tags in live cells. (**A**) Schematic of yellow L7Ae-Nun lantern assembly mediated by the *mC-4-B* RNA tag. Binding of L7Ae-VenusΔC12-SmBiT and Nun-LgBiT to the CS2 and boxB motifs, respectively, promotes NanoBiT complementation and light emission. The mutant tag (*mC-4-B^mut^*) disrupts Nun-LgBiT binding and prevents lantern assembly. (**B**) HEK293T cells stably expressing the yellow L7Ae-Nun lantern were transiently transfected with *in vitro* transcribed *mC-4-B* and *mC-4-B^mut^* RNA tags or left untransfected (no RNA). Cells were imaged using a super-cooled EMCCD camera. VenusΔC12 fluorescence (green) indicates lantern expression. Bioluminescence (magenta) was observed primarily in samples transfected with the cognate RNA tag. Scale bars = 50 μm.

### Single-vector co-expression and optimization of expression of RNA lantern–tag complexes

To enable coordinated expression of both the lantern and RNA tag components in cells, we developed a single-vector, bidirectional system encoding the RNA lantern and its cognate tag. In this construct, opposing promoters direct expression of the lantern and RNA tag through distinct RNA polymerases. A cytomegalovirus (CMV) promoter drives RNA polymerase II-based bicistronic expression of the lantern components via an internal ribosome entry sequence (IRES), ensuring coordinated expression of L7Ae-VenusΔC12-SmBiT and Nun-LgBiT. In the opposing direction, a minimal U6 (miniU6) promoter drives expression of the RNA tag, followed by a minimal HDV-like ribozyme (*drz-Mtgn-3*) to generate a defined 3′ end, and a downstream poly-T tract that serves as the RNA polymerase III termination signal (**Fig. 4A**)(Preece et al. 2020; Riccitelli et al. 2014; Gao et al. 2018). In contrast to previous strategies that required separate delivery of the lantern or generation of stable cell lines, this approach enables compact, single-vector delivery of a complete imaging system.

**Figure 4.**
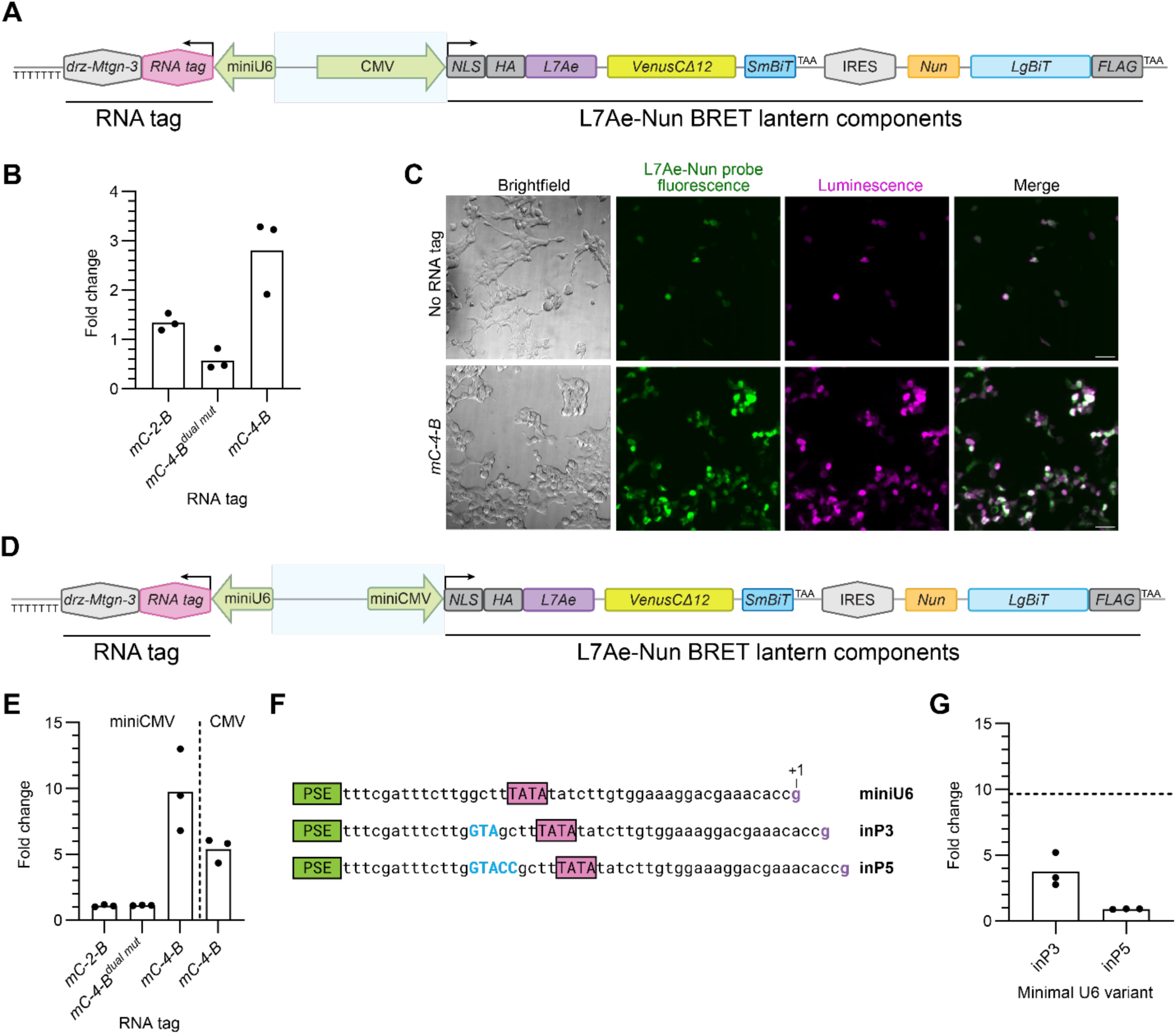
Evaluation of single-plasmid co-expression of RNA lantern and cognate tag in live cells. (**A**) Schematic of the bidirectional expression construct used for co-expression of the RNA lantern and its cognate RNA tag. A minimal U6 (miniU6) promoter drives expression of the RNA tag followed by a small HDV-like ribozyme (*drz-Mtgn-3*, 54 nts) to generate a defined 3′ end, and a poly-U tract (7 nts) that serves as the RNA polymerase III termination signal. In the opposite direction, a CMV promoter with an enhancer drives expression of the yellow L7Ae-Nun lantern, with L7Ae-VenusΔC12-SmBiT and Nun-LgBiT separated by an internal ribosome entry sequence (IRES). A nuclear localization signal (NLS) is fused to L7Ae-VenusΔC12-SmBiT to separate the two proteins in cells and improve signal- to-noise ratio. HA and FLAG epitope tags were included to enable downstream biochemical analyses. (**B**) Fold change in bulk luminescence over no-RNA tag control. Wildtype HEK293T cells were transfected with the co-expression vector containing the full length CMV promoter and enhancer, with or without RNA tag expression from the miniU6-driven operon. Inactive tag controls (*mC-2-B* and *mC-4-B^dual^ ^mut^*) exhibit luminescence comparable to the no-RNA control. (**C**) Bioluminescence imaging of HEK293T cells transiently expressing the L7Ae-Nun BRET lantern alone or in the presence of the *mC-4-B* tag. VenusΔC12 fluorescence (green) indicated RNA lantern expression. Bioluminescence (magenta) was observed primarily in cells containing the *mC-4-B* tag. Scale bars = 50 μm. (**D**) Schematic of the bidirectional expression construct as in (A), except with lantern expression driven by a minimal CMV (miniCMV) promoter. All other construct elements are identical. (**E**) Fold change in bulk luminescence over no-RNA tag control. HEK293T cells were transfected with the co-expression vector containing the miniCMV, in the presence or absence of minimized tag expression driven by the miniU6 promoter. Inactive tag controls (*mC-2-B* and *mC-4-B^dual^ ^mut^*) remain at background levels. (**F**) Modified miniU6 promoters (inP3 and inP5) showing the indicated spacing changes between the proximal sequence element (PSE, green) and the TATA box (pink) compared to the wildtype (WT) minimal U6. The inserted bases are capitalized (blue). The +1 “g” (purple) denotes the canonical transcription start site. (**G**) Fold change in bulk luminescence for constructs containing modified miniU6 promoters (inP3 and inP5) driving *mC-4-B* tag expression, relative to the no-RNA tag control. Data are derived from the same experiment shown in (E) and share the controls; the dotted line indicates the fold change for the miniU6 promoter. Data in (**B**), (**E**), and (**G**) are presented as mean values for *n* = 3 replicates, with individual data points shown.

The co-expression construct was initially transfected into HEK293T cells to evaluate lantern expression and RNA-dependent assembly. Bulk luminescence measurements were used to assess overall signal across a population of cells, enabling direct comparison of signal-to-noise ratios and overall photon emission. Cells expressing the functional *mC-4-B* tag exhibited robust luminescence following substrate addition, whereas constructs containing the dual mutant tag (*mC-4-B^dual^ ^mut^*), in which both CS2 and boxB aptamers are scrambled, and a structurally inactive tag (*mC-2-B)* generated luminescence comparable to the no-RNA control (**Fig. 4B; Supplementary Fig. 11A**). Live-cell imaging of HEK293T cells transfected with vectors encoding the lantern either in the presence or absence of the RNA tag further supported the conclusion that lantern assembly is RNA-dependent (**Fig. 4C**).

Having established RNA-dependent lantern assembly, we next sought to optimize the relative expression levels of the lantern components and RNA tag. To modulate lantern expression levels, a minimal CMV promoter (miniCMV) was also evaluated (**Fig. 4D**). HEK293T cells are highly transfectable, and in combination with the full-length CMV promoter and enhancer, they express lantern components at relatively high levels. This elevated expression can increase background signal by shifting the stoichiometric balance between lantern components and the RNA tag and lead to increased basal background(DuBridge et al. 1987; Xia et al. 2006). In contrast, using the miniCMV resulted in lower lantern expression but improved overall sensitivity by lowering background signal, at the cost of reduced signal intensity (**Fig. 4E; Supplementary Fig. 11B)**. The full-length promoter and enhancer produced substantially higher photon output, which can be advantageous for rapid signal acquisition or imaging through thick tissue, despite elevated background signal. Importantly, with both designs, clear separation between cognate and control constructs was maintained (**Fig. 4B, E**).

In parallel, to probe lantern assembly across a range of RNA tag expression levels and assess how tag abundance influences signal output, we decreased RNA tag expression by introducing small insertions (3 or 5 nts) into the miniU6 promoter between the proximal sequence element (PSE) and TATA box. This region is known to influence RNA polymerase III transcriptional output of the full-length U6 promoter (**Fig. 4F**), such that a 3-nt insertion reduces transcriptional activity to ∼40% of wildtype U6 levels, whereas a 5-nt insertion results in a more pronounced loss of activity(Goomer and Kunkel 1992). Consistent with reduced transcriptional efficiency, both modified miniU6 variants (inP3 and inP5) exhibited decreased luminescence relative to the reference promoter. The inP3 variant showed a reduction in luminescence by 5-fold, compared to the miniU6, whereas further extension to inP5 resulted in a near-complete loss of signal, approaching levels for the no-RNA and inactive tag controls (*mC-2-B* and *mC-4-B^dual^ ^mut^*). (**Fig. 4E, G; Supplementary Fig. 11C**). Reducing RNA tag expression (inP3 and inP5) diminished signal without an accompanying increase in fold change, indicating that RNA tag levels are not in excess relative to lantern component levels under miniU6-driven expression. These results demonstrate that lantern assembly efficiency is sensitive to RNA tag abundance and suggest that the reference miniU6-driven promoter provides a balance between RNA tag and lantern component expression (**Fig. 4G**).

The performance of the lantern–tag co-expression system was further evaluated in additional cell lines, including U-251 glioblastoma and HepG2 hepatocellular carcinoma cells, which differ from HEK293T cells in transcriptional activity and ploidy. These cells were transiently transfected with varying amounts of the co-expression vector to assess RNA imaging performance across diverse cellular environments.

In U-251 cells, RNA-dependent lantern assembly was observed, with peak signal attained 48 hours post-transfection. These cells exhibited a 5-fold increase in luminescence over the no-RNA tag control and a modest luminescence output at the optimal condition (300 ng DNA) (**Supplementary Fig. 12A, C**). The miniCMV-driven construct produced lower luminescence and did not yield clearly distinguishable RNA-dependent activation, reflecting suboptimal lantern expression (**Supplementary Fig. 12B, C**). In HepG2 cells, the CMV-driven construct produced higher total luminescence, but also elevated background signal, resulting in a ∼3.5-fold increase in luminescence at the optimal condition (300 ng DNA). Peak signal was observed 48 hours post-transfection, with an increase in signal accompanied by proportional increases in background luminescence (**Supplementary Fig. 12D, E**). As observed in U-251 cells, the miniCMV construct produced low luminescence with negligible separation from background when the RNA tag was present under the same conditions (**Supplementary Fig. 12F, G**).

For downstream imaging studies in both U251 and HepG2 cells, the full-length CMV-driven construct was selected, because it provided the strongest RNA-dependent signal in each cellular context. Cells expressing the *mC-4-B* tag were compared to no-tag controls by live-cell imaging, demonstrating that luminescence remained dependent on RNA-mediated lantern assembly (**Supplementary Fig. 13A and 14A**). However, in both cancer cell lines, consistent differences in VenusΔC12 fluorescence were observed during imaging and bulk luminescence measurements, with tagged constructs exhibiting modestly increased fluorescence relative to the no-tag controls. Similar increases in fluorescence were also observed in bulk measurements with the *mC-2-B* tag, which produces weaker luminescence activation than *mC-4-B* (**Supplementary Fig. 13B–D and 14B–D**). These observations are consistent with a model in which lantern binding with cognate RNA tags partially stabilizes the protein components, increasing their steady-state levels, and resulting in an increase in both fluorescence and luminescence.

Together, the live-cell imaging results demonstrate that the co-expression vector supports coordinated production of the lantern and tag in live cells. This design enables compact, single-vector delivery of the complete imaging system, facilitating rapid evaluation of tagged transcripts and providing a flexible framework for future integration of engineered RNA-based devices. Modulating lantern expression allows tuning between overall signal intensity and signal-to-noise ratio, with higher expression favoring brightness and lower expression improving sensitivity. We anticipate bright configurations to be advantageous for imaging in animal models, whereas lower-expression conditions enhance detection of RNA-dependent luminescence activation in cell culture. We also anticipate that compatibility across cell types will support broader application of the platform for imaging cell-type-specific transcripts.

Since RNA-dependent lantern assembly depends on the relative abundance of tagged RNAs and lantern components, factors such as transfection efficiency, promoter strength, and transcriptional activities of RNA polymerases II and III are expected to influence detection performance. Therefore, achieving sufficient lantern expression relative to target RNA levels will be critical for maximizing signal production across diverse cell types.

### Incorporation of minimized tags in structured RNA elements

We next asked whether the minimized tags could be incorporated into structurally constrained RNAs. Many transcripts, particularly functional RNAs that do not code for proteins, contain highly folded domains that are sensitive to sequence insertions or structural perturbations. These elements often perform critical molecular roles, such as quaternary interactions with other molecules and catalytic activity(Dethoff et al. 2012; Fedor and Williamson 2005). Consequently, such RNAs are challenging targets for conventional RNA imaging strategies, which often rely on large insertions that can disrupt native RNA structure and function.

To determine whether the minimized tags could be tolerated within structured RNAs, we incorporated them (lacking the P1 helix) into two well-characterized structured RNAs. First, we inserted the tag into the expression and nuclear retention element (*ENE*) of the lncRNA metastasis-associated lung adenocarcinoma transcript 1 (*MALAT1*). Mature *MALAT1* is a highly conserved ∼8-kb lncRNA implicated in transcript regulation, RNA processing, and cancer progression(Gutschner et al. 2013; Tripathi et al. 2010; Arun et al. 2020). Unlike most RNAs produced by RNA polymerase II, *MALAT1* does not undergo canonical polyadenylation. Instead, its precursor transcript is processed by RNase P to generate a mature 3′ end containing a short A-rich tract and a downstream tRNA-like domain known as MALAT1-associated small cytoplasmic RNA (mascRNA)(Wilusz et al. 2008; Brown et al. 2012; Wilusz et al. 2012). The 3′ end of mature *MALAT1* contains the *ENE+A* triple helix, a unique structural element, consisting of a conserved U-rich internal loop that forms a triple helix with the 3′ terminal A-rich tail, that stabilizes the transcript by protecting it from exonucleolytic degradation(Wilusz et al. 2008; Brown et al. 2014). This domain also includes a peripheral, sequence-divergent stem-loop. Because *MALAT1* lacks a conventional 3′ UTR and undergoes post-transcriptional processing, it represents a challenging transcript to tag using conventional imaging strategies. Previous studies have shown that the *ENE+A* core is sufficient to preserve transcript stability comparable to the wild-type *MALAT1 ENE*(Brown et al. 2014). We therefore introduced the *mC-4-B* tag by replacing the peripheral stem-loop and fusing the modified tag to the *MALAT1 ENE+A* core, rather than appending the tag to the terminus, to evaluate whether the minimized tag could be accommodated within a structured RNA element while still supporting lantern assembly (**Fig. 5A**).

**Figure 5.**
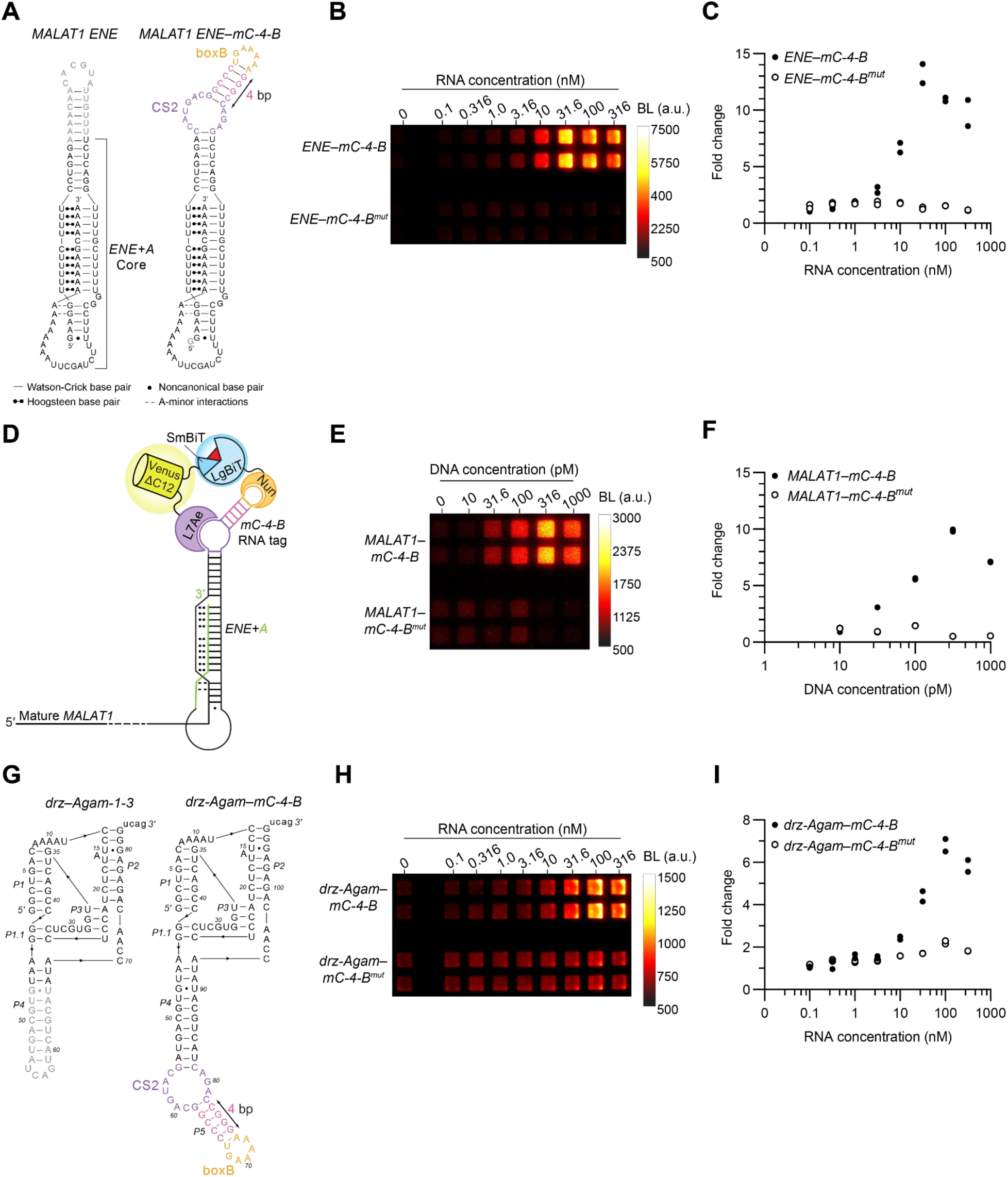
*In vitro* evaluation of minimized tags incorporated into highly structured RNAs. (**A**) Design schematic of the expression and nuclear retention element (ENE) from human *MALAT1* with incorporation of the minimized RNA tag. The native *ENE* peripheral stem-loop (light grey) was replaced with the *mC-4-B* tag lacking the P1 closing helix and fused to the *MALAT1 ENE+A core* (*ENE–mC-4-B*) to assess compatibility with a structured RNA scaffold. The *ENE* triple-helix is preserved through interactions between the U-rich internal loop and the 3′ A-rich tract. Hydrogen bonding interactions are annotated as follows: dashes (−) indicate Watson-Crick base pairs, dots (•) denote non-canonical base pairs, Leontis-Westhof notation(Leontis and Westhof 2001) for Hoogsteen base pairs, and double dashes (--) represent A-minor interactions. (**B**) Luminescence of *ENE–mC-4-B* versus mutant *ENE–mC-4-B^mut^* titrated into HEK293T lysate containing the yellow L7Ae-Nun lantern. (**C**) Fold change in luminescence signal of *ENE–mC-4-B* and mutant relative to the no-RNA control. Individual data points (*n* = 2) are shown per condition. (**D**) Schematic representation of *MALAT1* lncRNA with a single copy of the *mC-4-B* RNA tag incorporated in the *ENE* domain (*MALAT1–mC-4-B*). (**E**) Luminescence from *MALAT1–mC-4-B* evaluated via an *in vitro* transcription/translation assay. (**F**) Fold change in bioluminescent signal of *MALAT1–mC-4-B* and mutant relative to the no-RNA control. Individual data points (*n* = 2) are shown per condition. (**G**) Design schematic of *drz-Agam-1-3* with an internal *mC-4-B* tag (*drz-Agam–mC-4-B*). The P4 helix of *drz-Agam-1-3* (light grey), which does not participate in self-cleavage, was used as the insertion site for the *mC-4-B* tag. (**H**) Bioluminescent output from *drz-Agam–mC-4-B* compared to the mutant *drz-Agam–mC-4-B^mut^* titrated in HEK293T cells expressing the yellow L7Ae-Nun lantern. (**I**) Fold change in bioluminescent signal of *drz-Agam– mC-4-B* and mutant relative to the no-RNA control. Individual data points (*n* = 2) are shown per condition. BL, bioluminescence; a.u., arbitrary units.

We first evaluated the isolated *ENE* construct tagged with the *mC-4-B* (*ENE–mC-4-B*) and a mutant control (*ENE-mC-4-B^mut^) in vitro* with HEK293T lysate expressing the yellow L7Ae-Nun lantern. The *ENE–mC-4-B* produced >10-fold luminescence signal increase compared to the no-RNA control at the optimal concentration. In contrast, the mutant construct did not exhibit a measurable increase in signal even at higher RNA concentrations (**Fig. 5B, C; Supplementary Fig. 15A**). These results indicate that the inserted minimized tag retains a functional conformation capable of recruiting the lantern components within the structured *ENE* scaffold. To further minimize the tagging footprint, a smaller variant (*mC-1-B*) was also incorporated into the *ENE* (*ENE–mC-1-B*) and compared to the *ENE–mC-4-B* with the L7Ae-Nun BRET lantern to determine if the reduced tag size could be accommodated without disrupting folding or lantern assembly (**Supplementary Fig. 16A–C**). Given that *mC-1-B* has a shorter P2 helix and reduced base-pairing interactions compared to *mC-4-B*, it is expected to have decreased thermodynamic stability and weaker structural organization. These features can increase conformational flexibility and reduce the likelihood of the smallest tag maintaining the structure needed for lantern binding. *ENE-mC-1-B* RNA remained functional when titrated across increasing concentrations, exhibiting clear RNA-dependent lantern assembly. However, as anticipated, this construct is not fully optimized for the yellow L7Ae-Nun BRET lantern, and overall luminescence output was approximately 2-fold lower than *ENE–mC-4-B* at the optimal concentration (100 nM) in lysate. This observation suggests that the smallest tag can be adopted into structured RNAs, albeit with modest reduction in signal output with this lantern variant (**Supplementary Fig. 16A–C**).

We next evaluated full-length *MALAT1* RNA containing the *mC-4-B* insertion within the *ENE* domain (*MALAT1–mC-4-B;* 8361 nts) (**Fig. 5D**). The tagged transcript as well as its mutant variant *(MALAT1–mC-4-B^mut^)* were generated by T7-driven *in vitro* transcription in a coupled *in vitro* transcription-translation (IVTT) system, in which the yellow L7Ae-Nun lantern components were expressed in rabbit reticulocyte lysate. To assess the dependence of lantern assembly on transcript expression levels, the DNA template encoding *MALAT1–mC-4-B* was titrated across reactions. Under these conditions, the tagged full-length transcript supported lantern assembly. At higher template concentrations, variability in background luminescence was observed with the mutant control, likely due to competition between transcription of the *MALAT1–mC-4-B^mut^*template and translation of the lantern components within the coupled IVTT system (**Fig. 5E, F; Supplementary Fig. 15B**). These results indicate that the minimized tag remains functional within the context of a structured element of a full-length lncRNA.

To further determine whether the minimized tag could be accommodated within highly structured RNAs, we examined the insertion of the tag into a catalytically active RNA. We selected the HDV-like ribozyme *drz-Agam-1-3*, which adopts a conserved, highly compact catalytic fold and exhibits a rapid cleavage rate constant (2.7 ± 0.2 min^-1^)(Webb et al. 2009; Webb and Lupták 2011). HDV-like ribozymes provide a stringent model system, because their compact catalytic cores are sensitive to perturbation(Reymond et al. 2009). A successful tag insertion without loss of self-scission would suggest that the minimized tag can be accommodated within functionally constrained RNAs.

The P4 stem-loop of *drz-Agam-1-3* does not directly contribute to catalysis and is tolerant to sequence substitution(Webb et al. 2009). We therefore replaced the loop of the P4 helix with the *mC-4-B* tag to generate *drz-Agam–mC-4-B* (**Fig. 5G**). Secondary structure predictions generated for both tagged and untagged constructs using IPknot++ suggested that the nested double-pseudoknot of the ribozyme is preserved following tag insertion(Sato et al. 2011).

Catalytic activity was evaluated by incorporating a short 5′ leader sequence (22 nts) upstream of both the tagged and untagged variants of *drz-Agam-1-3*, enabling the detection of cleavage products (**Supplementary Fig. 17A**). To assess cleavage efficiency, transcripts were generated by *in vitro* transcription and assayed across a range of MgCl_2_ concentrations. Mg^2+^ ions are required for both efficient transcription by T7 RNA polymerase and for proper folding and catalytic activity of HDV-like ribozymes, where it primarily stabilizes the folded structure and directly participates in the catalysis. Varying Mg^2+^ concentrations were selected to probe both suboptimal and saturating conditions. In the presence of ∼8 mM total rNTPs, transcription reactions were incubated at 7 mM and 9 mM Mg^2+^ to represent suboptimal and near-optimal conditions, respectively, and at 16 mM Mg^2+^ to achieve higher ribozyme activity. Cleavage products were analyzed by denaturing PAGE. Transcription reactions at lower Mg^2+^ concentrations showed some accumulation of uncleaved precursor RNA, providing a reference to confirm that the observed bands arise from ribozyme-mediated cleavage. *drz-Agam–mC-4-B* exhibited efficient self-cleavage across all Mg^2+^ conditions, comparable to the wild-type construct, confirming that insertion of the minimized tag does not disrupt catalytic function (**Supplementary Fig. 17B**).

Furthermore, *drz-Agam–mC-4-B* supported efficient lantern assembly relative to the mutant variant (*drz-Agam–mC-4-B^mut^*) when assessed in HEK293T lysate expressing the yellow L7Ae-Nun lantern (**Fig. 5H, I; Supplementary Fig. 15C**). Background luminescence was elevated, likely due to higher lantern expression and variability arising from transient transfection in the lysate preparation. At higher RNA concentrations, the mutant construct exhibited low but detectable signal, likely reflecting residual, low-affinity Nun-LgBiT binding, given that the single-point mutation does not fully abrogate the interaction. Collectively, these results show that the tagged ribozyme remains catalytically active and structurally compatible with lantern assembly, supporting its potential for visualization of structured RNAs in cellular contexts.

Live-cell imaging of highly structured RNA constructs was next evaluated using *ENE–mC-4-B* and *drz-Agam–mC-4-B* under the same conditions established for minimized tag imaging (**Fig. 6A**). RNAs were *in vitro* transcribed and transfected into HEK293T cells stably expressing the yellow L7Ae-Nun lantern. After 24 hours, luciferin was added and luminescence was imaged using an EMCCD camera. VenusΔC12 fluorescence was used to assess lantern expression. Under these conditions, cells receiving the functional tagged constructs exhibited bright bioluminescence, compared to mutant variants (**Fig. 6B; Supplementary Fig. 18**). Additional control constructs, including variants with a fully scrambled boxB sequence (*ENE–mC-4-B^scramble^*) or point mutations in both aptamers (*ENE–mC-4-B^dp^ ^mut^*), exhibited abrogated bioluminescence signal (**Supplementary Fig. 18**). These observations demonstrate that minimized RNA tags can be incorporated into tightly folded RNAs without disrupting their ability to support lantern assembly in live cells. Signal generation remains strongly dependent on tag integrity, indicating that observed luminescence predominantly arises from RNA-specific interactions.

**Figure 6.**
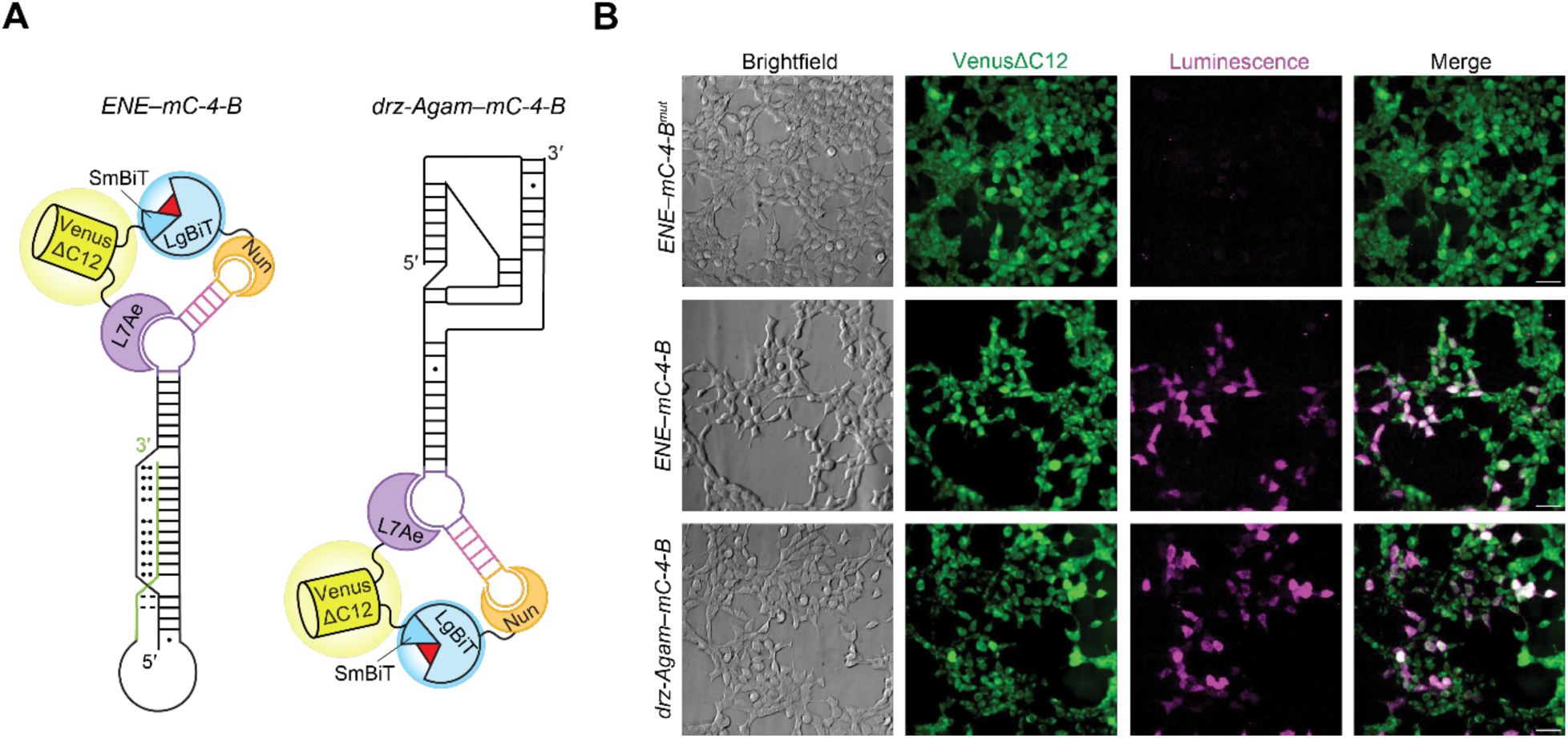
Bioluminescence imaging of highly structured RNAs with internal minimized RNA tag in live cells. (**A**) Schematic of *ENE–mC-4-B* and *drz-Agam–mC-4-B* RNA mediating assembly of the yellow L7Ae-Nun BRET lantern. (**B**) HEK293T cells stably expressing the yellow L7Ae-Nun lantern were transfected with *in vitro* transcribed *ENE–mC-4-B*, its mutant variant (*ENE–mC-4-B^mut^*), or with *drz-Agam–mC-4-B*. Cells were imaged using a super-cooled EMCCD camera. VenusΔC12 fluorescence (green) indicates lantern expression. Bioluminescence (magenta) as a result of lantern assembly. Scale bars = 50 μm.

## Discussion

Tracking RNA dynamics in living systems remains challenging due to limited availability of tools that support long-term imaging under physiological conditions. Conventional fluorescence-based approaches rely on external excitation, which can introduce phototoxicity and high background levels, thereby limiting their use in continuous or deep tissue imaging applications. In contrast, bioluminescence-based approaches do not require external illumination to generate photons, thus producing far less background signal and enabling serial imaging. Despite these advantages, bioluminescence-based strategies for RNA imaging remain limited.

In prior work, we developed a bioluminescent platform for RNA imaging that utilizes split-luciferase fragments each fused to unique RBPs (RNA lanterns). These lanterns bind to cognate RNA aptamer tags, enabling proximity-driven complementation of the luciferase fragments and subsequent light emission. Early-generation lanterns employed multimeric RNA binding proteins, including MCP and PCP, and facilitated RNA imaging in live cells and animal models(Halbers et al. 2024; Fuller et al. 2026). However, broader application of this system has been limited by the size of the lantern–tag complex, restricting compatibility with structured or functionally constrained RNAs.

In this work, a miniaturized bioluminescent RNA imaging platform was developed by combining a previously characterized monomeric RNA lantern (L7Ae-Nun) with newly minimized RNA tags. Unlike the MCP-PCP lantern, which employs dimeric RBPs and is therefore predicted to recruit at least two MCP and two PCP lantern fragments per MS2-PP7 RNA tag, the L7Ae-Nun lantern utilizes monomeric RBPs, reducing the overall protein load imposed on the target RNA. This reduction is reflected in the smaller size of the lantern components in the absence of fluorescent protein fusion partners (for the blue MCP-PCP lantern, the predicted dimeric assembly molecular weight is ∼93 kDa, whereas the blue L7Ae-Nun lantern is ∼33 kDa).

This molecular tool was further miniaturized by reducing the original 4-way junction RNA tag down to its core elements—two RNA binding domains, CS2 and boxB, fused to one another via a short helical spacer. Reduction of the RNA element yielded functional tags as small as 24 nts, representing a substantial decrease in size compared to conventional MS2/PP7 systems. Tag size reduction maintained efficient lantern assembly and BRET efficiency, preserving bright and spectrally resolved bioluminescence. This small RNA element enables insertion directly into compact or structured regions of RNAs, rather than requiring fusion to transcript termini, thereby minimizing disruption to native RNA structure, localization, and cellular behavior.

Structure-guided optimization revealed that efficient lantern assembly depends on both spacing and rotational alignment of the aptamer-binding domains. Repositioning of the aptamers improved signal output, while further truncation defined a lower size limit for functional tags, highlighting the balance between structural stability and minimal footprint. These findings establish design principles for compact RNA scaffolds and insertion strategies into transcripts of interest that support split-luciferase complementation.

The minimized tags supported RNA-dependent signal generation in lysate and live cells, and a single-vector co-expression system enabled coordinated delivery of the lantern and tag components with tunable expression. Signal output was sensitive to relative concentrations of both lantern and RNA, with either insufficient or excess RNA tag reducing signal intensity. These observations are consistent with concentration-dependent effects, such as the hook effect, and emphasize the importance of maintaining balanced lantern component expression for optimal imaging performance. Evaluation across other cell types further demonstrated that the lantern platform supports RNA imaging in diverse cellular environments, although optimal lantern expression levels vary and must be considered in future applications.

Furthermore, the compact tags remained functional when embedded within highly structured and functional RNAs, including the *MALAT1 ENE* and an HDV-like ribozyme, both *in vitro* and in live cells. This experimental validation highlights a key advantage of the miniature platform over conventional imaging approaches that rely on larger or repetitive tag insertions, often incompatible with structured or catalytically active transcripts.

Together, this work establishes a compact and adaptable bioluminescent RNA imaging platform that expands access to transcripts compatible with live-cell imaging. We anticipate the reduced footprint will facilitate interrogation of structurally constrained RNAs while preserving continuous visualization of RNA dynamics across biological scales.

## ACKNOWLEDGMENTS

We would like to thank the members of the UCI RNA Imaging group for their insightful discussions.

## AUTHOR CONTRIBUTION

Conceptualization: O.S., J.A.P., and A.L.; Methodology: C.E.T.C., E.B.F, C.C.C., J.A.P., and A.L.; Investigation: C.E.T.C., E.B.F, C.C.C., O.R., and N.T.S.; Formal analysis: C.E.T.C. and E.B.F.; Writing—original draft: C.E.T.C., J.A.P., and A.L.; Writing—review and editing: All authors; Visualization: C.E.T.C.; Funding acquisition: O.S., J.A.P., and A.L.

## SUPPLEMENTAL INFORMATION

### MATERIALS AND METHODS

#### General cloning methods

Cloning was performed using the Q5 DNA polymerase kit (New England Biolabs). Restriction enzymes and all buffers were obtained from New England Biolabs. dNTPs were purchased from ThermoFisher Scientific. Primers were purchased from Integrated DNA Technologies (IDT). Luria-Bertani medium (LB) was purchased from Genesee Scientific, and antibiotics (ampicillin and carbenicillin) were obtained from Sigma-Aldrich. Plasmids were sequenced by Azenta Life Sciences or Plasmidsaurus. All primers and plasmid stocks were stored at −20 °C unless otherwise specified. Plasmid maps were designed and sequencing traces were analyzed using Benchling. Plasmid construct details are listed in List S1.

Genes of interest were amplified via PCR using the Q5 DNA polymerase kit, supplemented with dNTPs. Site-directed mutagenesis was performed using Q5-based PCR with mutagenic primers containing the desired nucleotide substitutions, followed by treatment with the KLD Enzyme Mix (New England Biolabs) to phosphorylate and ligate the synthesized DNA, and degrade template DNA according to the manufacturer’s instructions. PCR reactions and Gibson assemblies were performed using a BioRad C3000 thermocycler. Genes of interest were amplified using the following conditions: (1) 95 °C for 3 min; (2) 95 °C for 30 s; (3) annealing temperature for primers, 30 s (4) 72 °C, time dependent on amplicon length; (5) repeat steps 2–4 repeated 20–times; (6) final extension at 72 °C for 5 min, then hold at 12 °C until retrieval. PCR products were analyzed by gel electrophoresis, excised, and purified using a ZymoClean Gel DNA Recovery Kit. Purified gene fragments were inserted into destination vectors via Gibson assembly under the following conditions: 50 °C for 60 min and hold at 12 °C until retrieval^1^. Assembled plasmids were transformed into TOP10 or DH5α *E. coli* competent cells by the heat shock method (42 °C for 45 s, then ice). Transformants were plated on LB agar supplemented with carbenicillin (100 µg/mL) and incubated at 37 °C for 18–24 h. Individual colonies were picked and expanded overnight in 5 mL LB broth with the ampicillin (100 µg/mL). Plasmid DNAs were extracted from bacterial cultures using the Zymo Research Plasmid Mini-prep Kit-Classic, analyzed by gel electrophoresis, then sequenced. Detailed plasmid construct information is provided in List S1.

#### General cell culture methods

HEK293T cells (HEK, ATCC) and stable cell lines derived from HEK293T cells were cultured in complete medium: Dulbecco’s Modified Eagle Medium (Corning) containing 10% (v/v) fetal bovine serum (FBS, Life Technologies), penicillin-streptomycin (100 U/mL, Gibco), and plasmocin prophylactic treatment (0.0025 mg/mL, Invivogen). Cell were maintained at 37 °C in a 5% CO_2_ humidified chamber and serially passaged using trypsin-EDTA (0.25 % in Hank’s Balanced Salt Solution, Gibco). Cell-counting was performed using an automated cell counter (Countess II, Invitrogen).

HepG2 cells (ATCC) were kindly provided by Brian Paegel at the University of California, Irvine. Cells were cultured in complete medium: Dulbecco’s Modified Eagle Medium (Life Technologies), supplemented with 10% (v/v) fetal bovine serum (FBS, Life Technologies) and penicillin-streptomycin (100 U/mL, Life Technologies). Cells were maintained at 37 °C in a 5% CO_2_ humidified chamber.

U-251 cells (ATCC) were kindly provided by Daniela Bota at the University of California, Irvine. Cells were cultured in complete medium: Dulbecco’s Modified Eagle Medium (Life Technologies), supplemented with 10% (v/v) fetal bovine serum (FBS, Life Technologies) and penicillin-streptomycin (100 U/mL, Life Technologies). Cells were maintained at 37 °C in a 5% CO_2_ humidified chamber.

#### Stable cell line generation

Stable cell lines expressing RNA lanterns were generated via lentiviral transduction and antibiotic selection. Lantern sequences were cloned into pLenti-puro vector backbones. pLenti-puro was gifted by le-Ming Shih (Addgene #39481)^2^. Packaging was performed using the psPAX2 plasmid (Addgene #12260) and the envelope-expressing plasmid pMD2.G (Addgene #12259), both gifted by Didier Trono. HEK293T cells (5.7 × 10^5^) were seeded in 6-well culture plates (Corning) and transfected ∼18 h later with pLenti-puro plasmids constructs together with psPAX2 and pMD2.G using Lipofectamine 3000 (3 μL/well, ThermoFisher) and P3000 reagent (3 μL/well, ThermoFisher). Approximately 20 h post transfection, polybrene (8 μg/mL, Thomas Scientific) was added to the growth medium. The following day, the medium was replaced with fresh medium containing puromycin dihydrochloride (20 μg/mL, Sigma Aldrich) for selection. Transduced cells were maintained in puromycin (20 μg/mL) to preserve stable gene integration.

#### HEK293T lysate preparation

HEK293T cells (∼6 × 10⁵) were plated in 6-well culture plates (Corning) and transfected ∼18 h later with plasmids encoding the RNA lantern components using Lipofectamine 3000 (3 μL/well, Thermo Fisher) and P3000 reagent (3 μL/well, Thermo Fisher). Cells were harvested for lysate preparation ∼18–24 h post-transfection. In parallel, HEK293T cells stably expressing the RNA lantern were harvested for lysate preparation.

Cells were lifted using trypsin–EDTA (0.25%), neutralized, and pelleted by centrifugation at 500 ×*g* for 5 min at 4 °C (Thermo Fisher Sorvall Legend Micro 17). Supernatants were discarded, and cells were washed with sterile phosphate-buffered saline (PBS, pH 7.4) and pelleted again by centrifugation. Cell pellets were resuspended in fresh sterile PBS and counted. Cells were lysed by passage through a 30-gauge needle (1 mL syringe; Fisher Scientific). Cell debris was removed by centrifugation at 500 ×*g* for 5 min at 4 °C. Clarified lysates were aliquoted and stored at −80 °C.

#### *In vitro* transcription of RNA tags

DNA templates encoding the RNA constructs, which contained a T7 promoter, were generated by primer extension of synthetic DNA oligonucleotides (Integrated DNA Technologies) using the Q5 DNA polymerase kit from New England Biolabs. DNAs encoding *mB-X-C, mB-3-C_b1*–*3,* or *mC-X-B* contained either a 5′ *hhrI-SewB1_01643s-1* hammerhead ribozyme^3^ and/or a 3′ *drz-Mtgn-3* ribozyme^4^.

RNAs were transcribed *in vitro* for 2 h at 37 °C in 1× transcription buffer [50 mM Tris-HCl pH 8, 2 mM spermidine, 0.01% Triton X-100, 2 mM rNTPs (each), 16 mM MgCl_2_, and 8 mM dithiothreitol] and recombinant T7 RNA polymerase. T7 polymerase was expressed using p6XHis-T7(P266L), which was gifted by Anna Pyle (Addgene plasmid # 174866)^5^. Transcription reactions were quenched with 30 mM EDTA and subsequently ethanol-precipitated with co-precipitant (Invitrogen GlycoBlue, AM9516). RNA was purified by denaturing polyacrylamide gel electrophoresis (12%, 7M urea). Purified RNAs were eluted from gel pieces into a solution containing 300 mM KCl and 0.1 mM EDTA over a 3 h period. Eluted RNAs were ethanol-precipitated with co-precipitant, and pellets were washed with 70% ethanol, dried, and resuspended in TET buffer (10 mM Tris-HCl pH 7.5, 0.1 mM EDTA, and 0.001% Triton X-100) supplemented with 30 mM KCl. RNAs were refolded by heating to 65 °C for 5 min, followed by cooling to room temperature. RNA concentrations were determined by UV-Vis absorption spectroscopy (Thermo Scientific NanoDrop 2000, ND-2000). See List S1 for RNA construct details.

#### Minimized RNA tag optimization

HEK293T cell lysate expressing the lanterns was prepared as described above then diluted in sterile PBS to the desired cell-equivalent concentration (∼10^6^ cells/mL). Purified RNAs (1 µM stocks) were diluted in TET supplemented with 30 mM KCl prior to use. Reactions (10 µL total volume) were assembled by diluting lysate to the final volume and adding RNA (1 µL) for a final concentration of 100 nM. Following assembly, 1 µL of furimazine stock (200 µM stock, Nano-Glo Luciferase Assay System, Promega) prepared in physiological-like buffer (140 mM KCl, 10 mM NaCl, 20 mM Tris-HCl pH 7.5, and 5 mM MgCl_2_) was added to each reaction. Reactions were transferred to a black, clear bottom 384-well plate (Nunc) and imaged from below using a supercooled (−100 °C) Andor iXon Ultra 888 EMCCD camera (Oxford Instruments) equipped with an f/0.8 lens (Voigtländer Super Nokton) at an EM gain of 500–1000. Images were collected (5– 60s acquisition) over 5–30 min. Images were processed in ImageJ (Fiji 3) by Z-stacking and denoising, and peak intensities were quantified by densitometric analysis. All data were exported to Microsoft Excel and Prism 10 (GraphPad) for further analysis.

#### *In vitro* transcription and translation (IVTT) of RNA lanterns in rabbit reticulocyte lysate

IVTT reactions were prepared on ice in a total volume of 10 μL with 70% v/v rabbit reticulocyte lysate (Promega, L4960), 0.02 mM amino acid mix, rNTPs (0.35 mM GTP and 0.15 mM each ATP/CTP/UTP), 1 mM MgCl_2_, 1 nM plasmid DNA template encoding the RNA lantern, 2 U RNase inhibitor (Invitrogen, AM2694), and 0.5 μL of recombinant T7 RNA polymerase. T7 polymerase was expressed using p6XHis-T7(P266L), which was gifted by Anna Pyle (Addgene plasmid #174866)^5^. Reactions were incubated at 34 °C for 10 min and 30 °C for 140 min.

#### RNA lantern complex pull downs

Pierce magnetic anti-FLAG beads (1 μL) were washed three times in physiological buffer (140 mM KCl, 10 mM NaCl, 20 mM Tris-HCl pH 7.5, 5 mM MgCl_2_, and 0.01% Tween-80) at 10× their initial volume (300 μL and 10 μL, respectively) for 5 min with agitation. For binding, 50 μL reactions containing HEK293T cell lysate expressing the yellow L7Ae-Nun lantern, with or without 100 nM of *mB-3-C* RNA tag, were added to the beads and incubated at room temperature for 1 h. Following incubation, the flow through was collected and stored on ice. Beads were then washed three times in physiological buffer (50 μL per wash), and all wash fractions were kept on ice. Beads were resuspended in physiological buffer (50 μL) supplemented with 20 μM furimazine (Nano-Glo Luciferase Assay System, Promega). Flow through and wash fractions (25 μL each) were similarly supplemented with 20 μM of furimazine. Samples were plated in duplicate (beads) or a single replicate (flow through and washes) in a black, clear bottom 384-well plate (Nunc). Reactions were imaged from below using a supercooled (−100 °C) Andor iXon Ultra 888 EMCCD camera (Oxford Instruments) equipped with an f/0.8 lens (Voigtländer Super Nokton) at an EM gain of 500. Images were acquired (10 s acquisition) over 5 min. Images were Z-stacked in ImageJ (Fiji 3), and peak intensities were quantified by densitometric analysis. Data were exported to Microsoft Excel and Prism 10 (GraphPad) for further analysis.

#### *In vitro* evaluation of modified *mB-3-C* and *mC-1-B* with truncated P1 helix

IVTT was performed as described above for the expression of the lanterns. Following incubation, each RNA construct (1 μL, 100 nM final) was added, followed by the addition of 20 µM furimazine. Reactions were plated to a black, clear bottom 384-well plate (Nunc) and imaged from below using a supercooled (−100 °C) Andor iXon Ultra 888 EMCCD camera (Oxford Instruments) equipped with an f/0.8 lens (Voigtländer Super Nokton) at an EM gain of 1000. Images were collected (5 min acquisition) over 30 min. Images were processed in ImageJ (Fiji 3) by Z-stacking and denoising, and peak intensities were quantified by densitometric analysis. All data were exported to Microsoft Excel and Prism 10 (GraphPad) for further analysis.

#### RNA construct titration in HEK293T lysate

Lysate from lantern-expressing cells was prepared as described above and diluted in sterile PBS to the desired cell-equivalent concentration (∼10^6^ cells/mL). Purified RNAs (10× stocks) were serially diluted in TET buffer supplemented with 30 mM KCl prior to use. Reactions (10 µL total volume) were prepared by diluting lysate to volume and adding RNA (1 µL) to the indicated final concentrations. Furimazine (1 µL of 200 µM stock) was then added to each reaction. Samples were transferred to a black, clear bottom 384-well plate (Nunc) and imaged from below using a supercooled (−100 °C) Andor iXon Ultra 888 EMCCD camera equipped with an f/0.8 lens (Voigtländer Super Nokton) at an EM gain of 500–1000. Images were collected (5–60 s acquisition) over 5–20 min then Z-stacked and denoised in ImageJ (Fiji 3). Densitometry was used to measure peak intensity values.

#### Minimized tag bioluminescence resonance transfer spectra

Titrations of *mC-4-B* and *mC-4-B^mut^* RNAs (0–316 nM) were set up as described above. Spectra were acquired using a TECAN Spark multimode microplate reader using the luminescence scanning method. Emission was measured (counts/s) at wavelengths ranging from 398 to 653 nm with a 15 nm step size and 1 s integration time using the SparkControl software.

#### Titration of *MALAT1–mC-4-B* and *MALAT1–mC-4-B^mut^*

Plasmid DNAs encoding tagged *MALAT1* lncRNA were 3′ linearized after the polyA tract for run-off transcription using Esp3I (New England Biolabs). DNAs were kit purified (DNA Clean and Concentrator; Zymo Research, D4004) and eluted with TET buffer. Linearized plasmids (10× stocks) were serially diluted in TET buffer prior to use. IVTT reactions (10 μL total volume) were prepared as described above supplemented with 1 µL of RNA construct DNA template to the indicated final concentrations. Reactions were incubated at 34 °C for 10 min and 30 °C for 140 min. Furimazine was added to each reaction at 20 µM then plated transferred to a black, clear bottom 384-well plate (Nunc) and imaged from below using a supercooled (−100 °C) Andor iXon Ultra 888 EMCCD camera equipped with an f/0.8 lens (Voigtländer Super Nokton) at an EM gain of 1000. Images were collected (5 min acquisition) over 1 hr then Z-stacked and denoised in ImageJ (Fiji 3). Densitometry was used to measure peak intensity values.

#### Bioluminescence microscopy of *in vitro* transcribed RNA constructs in live cells

HEK293T cells (∼4 × 10^4^) stably expressing the RNA lantern were plated in fibronectin-coated (Sigma-Aldrich) 8-well Ibidi µ-Slides. After 24 h, the cells were transiently transfected with 200 ng of *in vitro* transcribed RNA constructs using 0.6 µL Lipofectamine 3000 and 0.6 µL P3000 reagent. After ∼24 h, cell media was replaced with phenol red-free DMEM (Gibco FluoroBrite DMEM) and supplemented with 20 µM furimazine (Nano-Glo Luciferase Assay System, Promega). Static live-cell images were captured on an Olympus IX71 microscope equipped with a Princeton Instruments ProEM-HS 512BX3 EMCCD camera cooled to –90 °C and a 20× air objective (Olympus UPLXAPO 20×). Luminescence images were captured with an EM gain of 1000, 10 MHz horizontal read-out rate, 4.33 μs vertical clock speed, and an acquisition time of 60 s. Multiple frames were Z-stacked to improve signal-to-noise. Fluorescent images were captured immediately before luminescence imaging with 100–500 ms acquisition, EM gain of 50, and a 470 nm blue LED excitation source (ThorLabs) with YFP emission filter (Olympus U-MF2). Images were processed (Z-stacked and brightness/contrast normalized) with ImageJ (Fiji3) and data were exported to Microsoft Excel and transferred to Prism 10 (GraphPad) for further analysis.

#### *In vitro* evaluation of tagged *drz-Agam-1-3* cleavage

DNA templates encoding *drz-Agam-1-3* and *drz-Agam-mC-4-B,* with or without a 5′ leader sequence (22 nts), were transcribed *in vitro* under the same conditions described above with MgCl_2_ concentrations of 7, 9, or 16 mM. Reactions were quenched immediately by buffer exchange using G-15 Sephadex into denaturing loading buffer (95% formamide, 25 mM EDTA, 0.5× Tris-borate-EDTA (TBE), bromophenol blue). Samples were resolved by denaturing PAGE (12%, 7M urea), then post-stained with 1× SYBR Gold (Invitrogen) in 0.5× TBE for 20 min. Fluorescence signal was detected using a Cytiva Amersham Typhoon Biomolecular Imager with 488 nm excitation (Cy2 channel, 525BP20 emission filter). Images were acquired at 100 µm pixel resolution with a PMT voltage of 369 V using ImageQuant TL software. See List S1 for *drz-Agam*-*1-3* and 5′ leader RNA sequences.

#### Initial bulk evaluation of RNA lantern and tag co-expression in varying cell-lines

##### HEK293T

2 x 10^4^ cells/well were plated in a black, clear bottom 96-well plate (Greiner). The next day, cells were transfected with DNA encoding either the lantern alone, or the lantern and tag together (100 ng/well) using Lipofectamine 3000 and P3000 reagent (0.3 µL each per well). After 24 h, cells were supplemented with furimazine (20 µM, Nano-Glo Luciferase Assay System, Promega) and total luminescence was measured using a TECAN Spark multimode microplate reader (1 s integration, controlled using SparkControl software).

##### HepG2

2.5 x 10^4^ cells/well were plated in a black, clear bottom 96-well plate. The following day cells were transfected with either 100, 300, or 500 ng of each DNA construct using Lipofectamine 3000 and P3000 reagent (0.15 µL and 0.2 µL, respectively per well). The next day, cells were supplemented with 20 µM furimazine and total luminescence was measured as described above. Bulk luminescence was measured 24, 48, and 72 h post-transfection. Cell culture media were replaced daily following measurement.

##### U-251

1 x 10^4^ cells/well were plated in a black, clear bottom 96-well plate in antibiotic-free media. The following day cells were transfected with either 100 or 300 ng of each DNA construct using Lipofectamine 3000 and P3000 reagent (0.15 µL and 0.2 µL, respectively per well). After ∼5 h, the media were replaced. The next day, cells were supplemented with 20 µM furimazine and total luminescence was measured as described above. Bulk luminescence was measured 24, 48, and 72 h post-transfection. Cell culture media was replaced daily following measurement.

#### Bulk fluorescence evaluation of RNA lantern and tag co-expression in varying cell-lines

HepG2 and U-251 cells were prepared as described above. After 72 h, cells were supplemented with 20 µM furimazine. Bulk fluorescence was measured using an Agilent BioTek Cytation 5 multimode microplate reader (excitation: 500/20 nm, emission:541/20 nm) controlled using Gen5 software. Bulk luminescence was subsequently measured using the same instrument with a 1 s integration time.

#### Bioluminescence microscopy of RNA lantern and tag co-expression vector in live cells

##### HEK293T

Wildtype HEK293T cells (∼4 × 10^4^) were plated per well in fibronectin-coated (Sigma-Aldrich) 8-well Ibidi µ-Slides. After 24 h, the cells were transiently transfected with 200 ng/well of DNA using 0.6 µL Lipofectamine 3000 and 0. 6 µL P3000 reagent. After ∼24 h, culture medium was replaced with with phenol red-free DMEM (Gibco FluoroBrite DMEM) and supplemented with 20 µM furimazine (Nano-Glo Luciferase Assay System, Promega).

##### HepG2

HepG2 cells (∼4 × 10^4^) were plated per well in 8-well Ibidi µ-Slides coated in collagen type I, rat tail (Advanced BioMatrix). After 24 h, the cells were transiently transfected with 1000 ng/well of DNA using 0.375 µL Lipofectamine 3000 and 0. 5 µL P3000 reagent. After ∼48 h, culture medium was replaced with phenol red-free DMEM (Gibco FluoroBrite DMEM) and supplemented with 20 µM furimazine (Nano-Glo Luciferase Assay System, Promega).

##### U-251

U-251 cells (∼2 × 10^4^) were plated per well in fibronectin-coated (Sigma-Aldrich) 8-well Ibidi µ-Slides. After 24 h, the cells were transiently transfected with 600 ng/well of DNA using 0.375 µL Lipofectamine 3000 and 0.5 µL P3000 reagent in antibiotic-free media. Culture medium was replaced ∼5 h post-transfection. After ∼48 h, culture medium was replaced with phenol red-free DMEM (Gibco FluoroBrite DMEM) and supplemented with 20 µM furimazine (Nano-Glo Luciferase Assay System, Promega).

Live-cell images were captured on an Olympus IX71 microscope equipped with a Princeton Instruments ProEM-HS 512BX3 EMCCD camera cooled to –90 °C and a 20× air objective (Olympus UPLXAPO 20×). Luminescence images were captured with an EM gain of 500–1000, 10 MHz horizontal read-out rate, 4.33 μs vertical clock speed, and an acquisition time of 1–10 s. Multiple frames were Z-stacked to improve signal-to-noise. Fluorescence images were captured before luminescence imaging with 100–500 ms acquisition, EM gain of 10–50, and a 470 nm blue LED excitation source (ThorLabs) with fluorescence emission filtered using a YFP emission filter (Olympus U-MF2). Images were processed (Z-stacked and brightness/contrast normalized) with ImageJ (Fiji3).

## Supplementary Information

**Figure S1.**
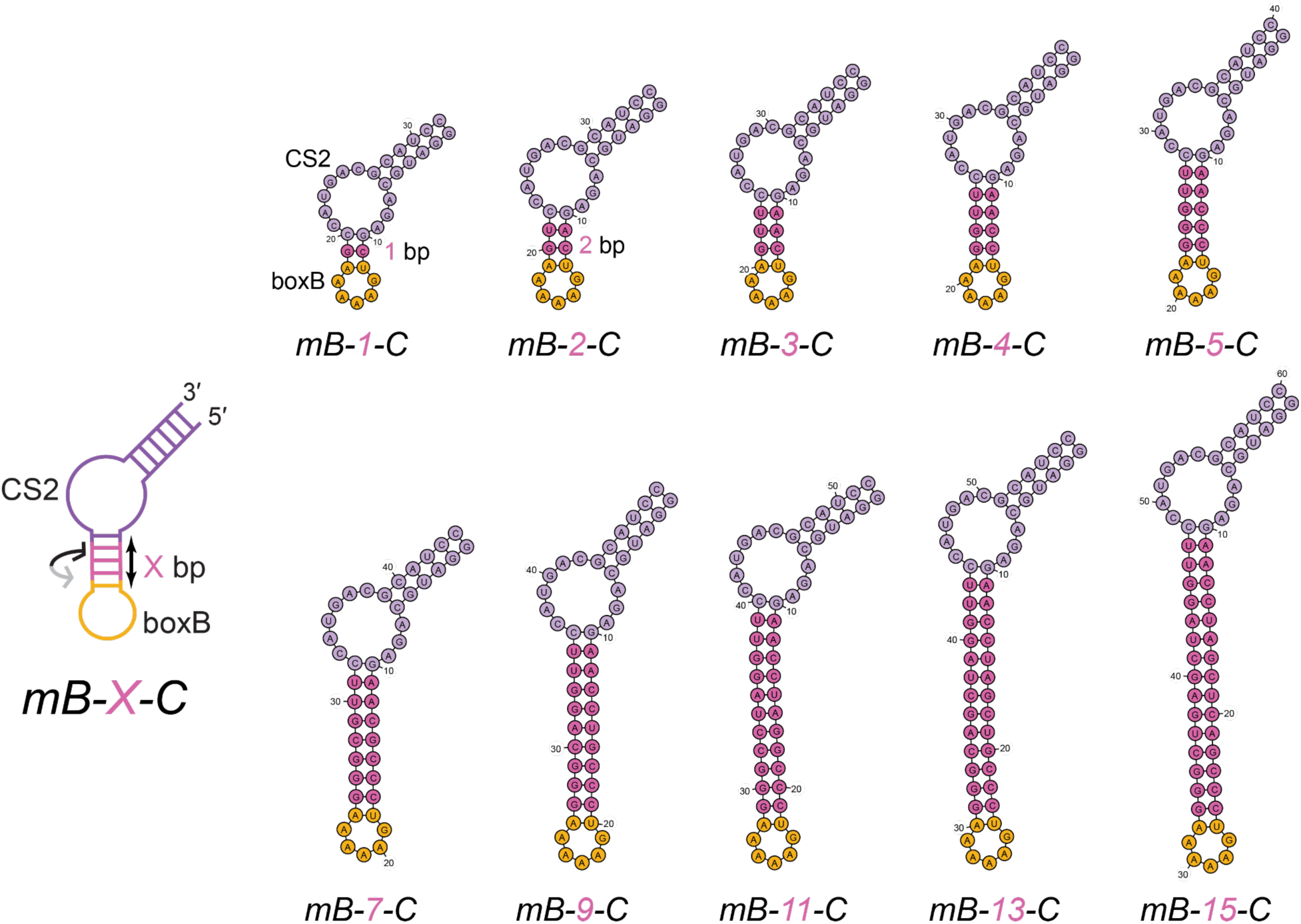
Initial design of minimized CS2-boxB RNA tags. Minimized tags were designed with a varying number of base pairs (X, pink) that modulate the helical phase and distance between the CS2 k-turn (purple) and boxB (orange) aptamers.

**Figure S2.**
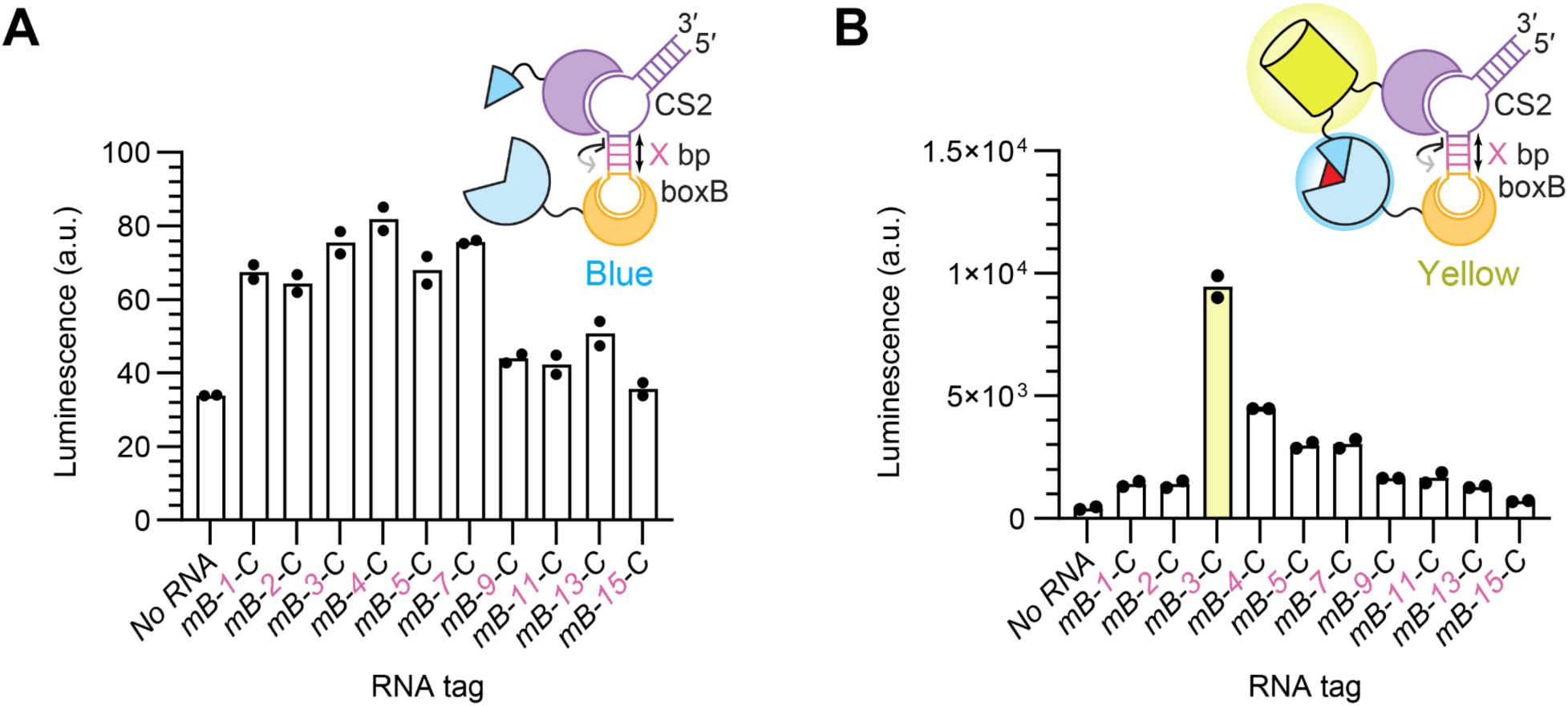
Luminescence of putative minimized RNA tag complexes. Individual components of the RNA lantern were expressed separately in HEK293T cells, their lysates were combined equally and supplemented with 100 nM of each minimized RNA tag (*mB-X-C*). The tags consist of boxB and CS2 aptamers fused within a single junction separated by varying spacer lengths (X denotes number of base pairs). Luminescence was imaged using a super-cooled EMCCD camera. (**A**) Total luminescence from *mB-X-C* panel with the blue L7Ae-Nun lantern, corresponding to images shown in **Fig. 1B** and fold changes plotted in **Fig. 1C**. (**B**) Total luminescence from *mB-X-C* panel using the yellow L7Ae-Nun BRET lantern, corresponding to images shown in **Fig. 1D** and fold changes plotted in **Fig. 1E**. Bars represent mean luminescence for each sample (*n* = 2), with individual data points shown. a.u., arbitrary units.

**Figure S3.**
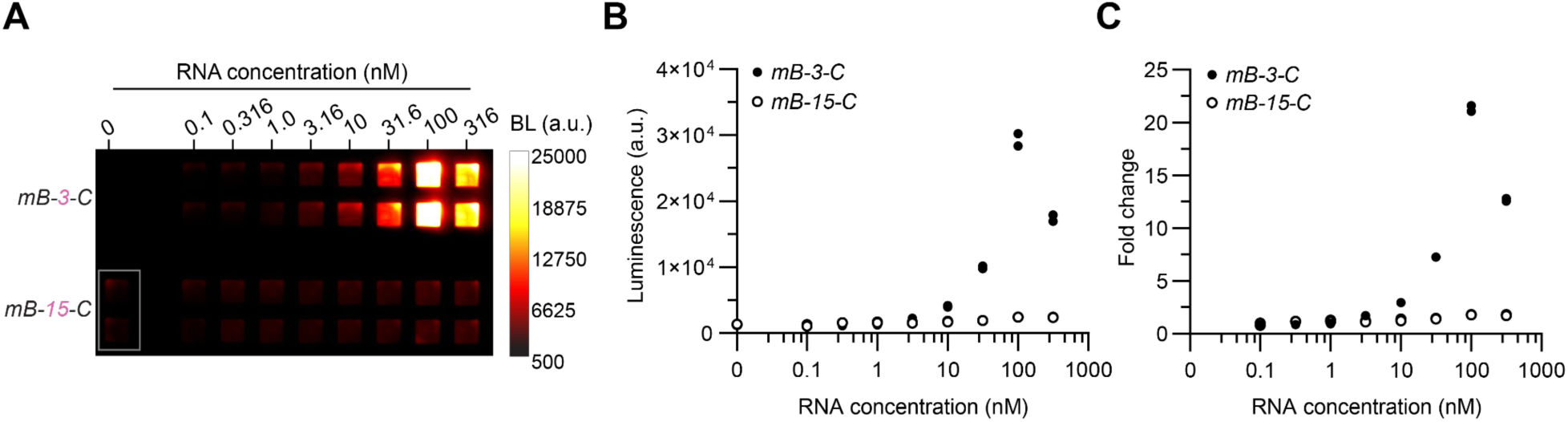
Minimized RNA tag structure influences lantern assembly. Each component of the yellow L7Ae-Nun BRET lantern was expressed separately in HEK293T cells, their lysates combined in equal proportions and supplemented with varying concentrations (0–316 nM) of each minimized RNA tag (*mB-3-C* and *mB-15-C*). (**A**) Luminescence of the optimal tag (*mB-3-C*) compared to the structurally inactive tag (*mB-15-C*), imaged using a super-cooled EMCCD camera. The boxed 0 nM RNA tag image indicates that the same no-RNA control was used for both conditions. (**B**) Relative luminescence of the *mB-3-C* and *mB-15-C* tags across RNA concentrations. (**C**) Fold change in luminescence relative to the no-RNA control (0 nM) for each tag across concentrations. A hook effect is observed with the *mB-3-C* tag at higher RNA concentrations. Individual data points (*n* = 2) per condition are shown. BL, luminescence; a.u., arbitrary units.

**Figure S4.**
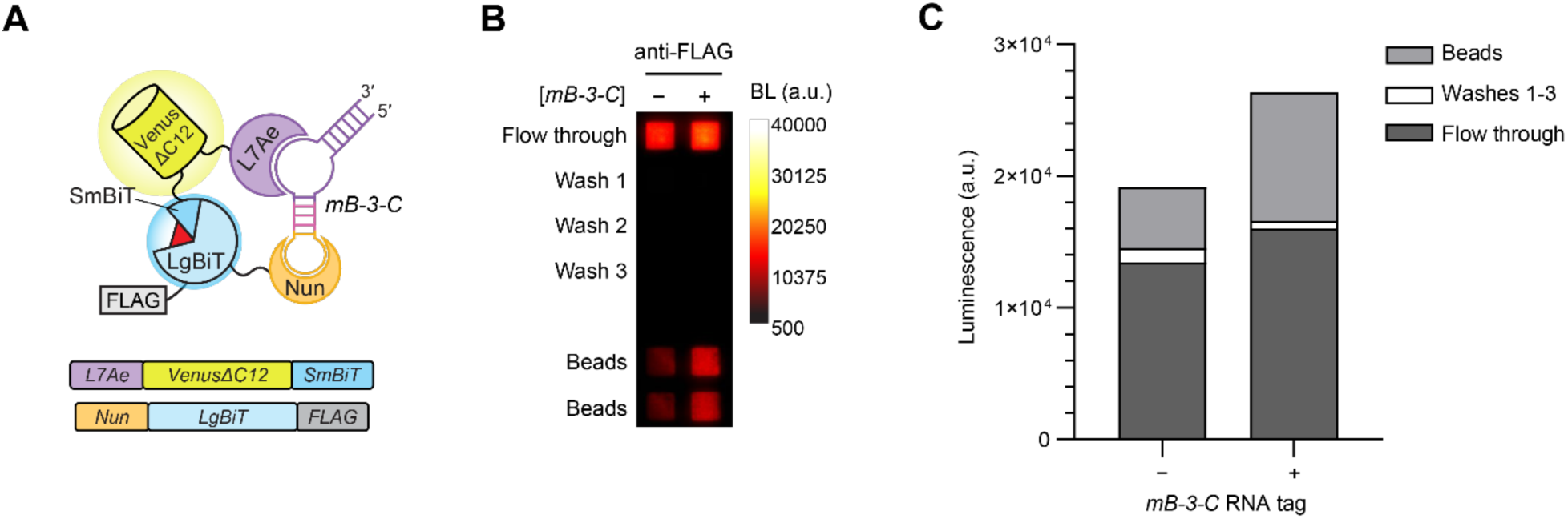
Lantern complex formation with minimized RNA tag. The L7Ae-Nun BRET lantern was expressed via *in vitro* transcription-translation (IVTT) and incubated in presence or absence of *mB-3-C* RNA tag. RNA lantern complexes were captured with magnetic anti-FLAG conjugate-agarose beads. Flow-through fractions were collected, and beads were washed three times prior to imaging. Luminescence was measured in the flow-through, wash fractions, and bead-bound complexes. (**A**) Schematic of the minimized RNA tag, comprising a single fusion of the boxB and CS2 aptamers, which mediate the assembly of RNA lantern components (Nun-LgBiT-FLAG and L7Ae-VenusΔC12-SmBiT). (**B**) Luminescence of lantern complexes captured using anti-FLAG conjugate-agarose beads. (**C**) Distribution of luminescence signal across bead-bound lantern complexes, washes, and flow-through fractions shown in (B), demonstrating luminescence increase resulting from RNA-bound lantern complexes. BL, luminescence; a.u., arbitrary units.

**Figure S5.**
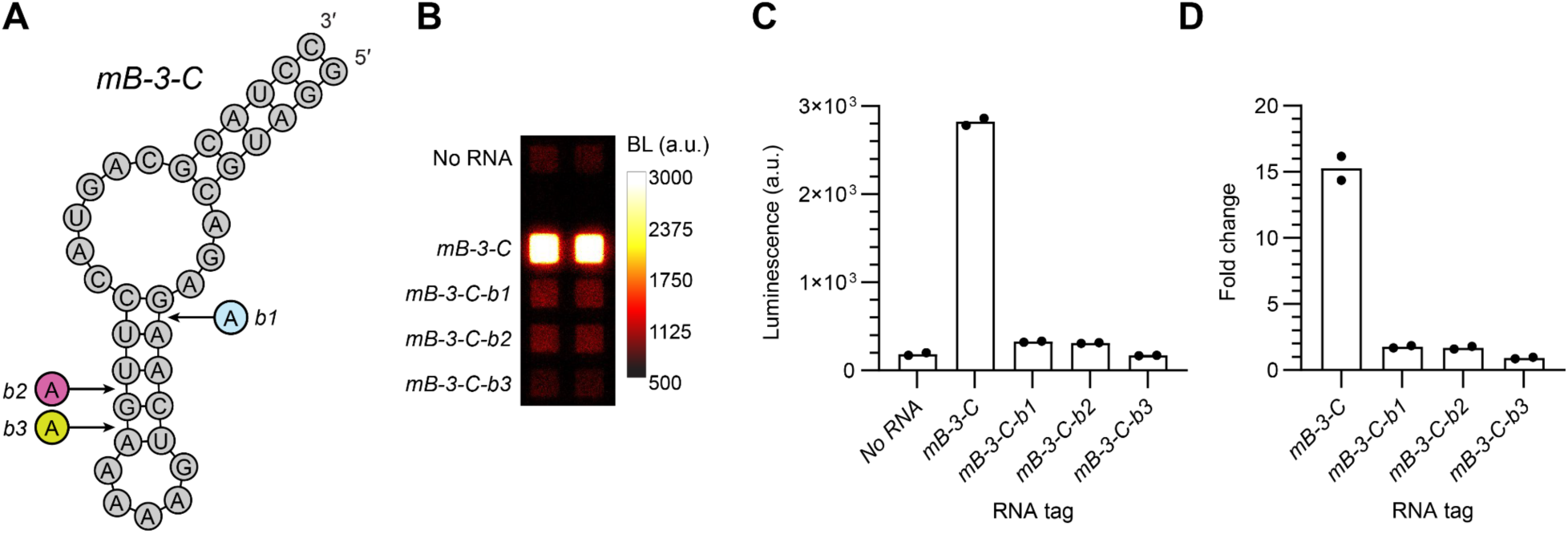
Tuning helical phase via single-nucleotide spacer insertions. The *mB-3-C* RNA tag was modified by introducing single-nucleotide insertions within the helical spacer region between the boxB and CS2 aptamers to tune relative orientation and assess effects on lantern assembly. The L7Ae-Nun BRET lantern was expressed by *in vitro* transcription/translation in reticulocyte lysate. (**A**) Schematic of RNA tag spacer variants (*mC-3-B-b1 to -b3)* containing single-adenosine insertions that introduce a bulge, designed to modulate alignment between lantern binding sites. (**B**) Luminescence of spacer variants compared to the parent *mB-3-C* tag. (**C**) Relative luminescence of yellow L7Ae-Nun lanterns with *mC-3-B-b1 to b3*. (**D**) Fold change over no-RNA control for each variant. Bars represent mean luminescence and fold change for each sample (*n* = 2), with individual data points indicated. BL, luminescence; a.u., arbitrary units.

**Figure S6.**
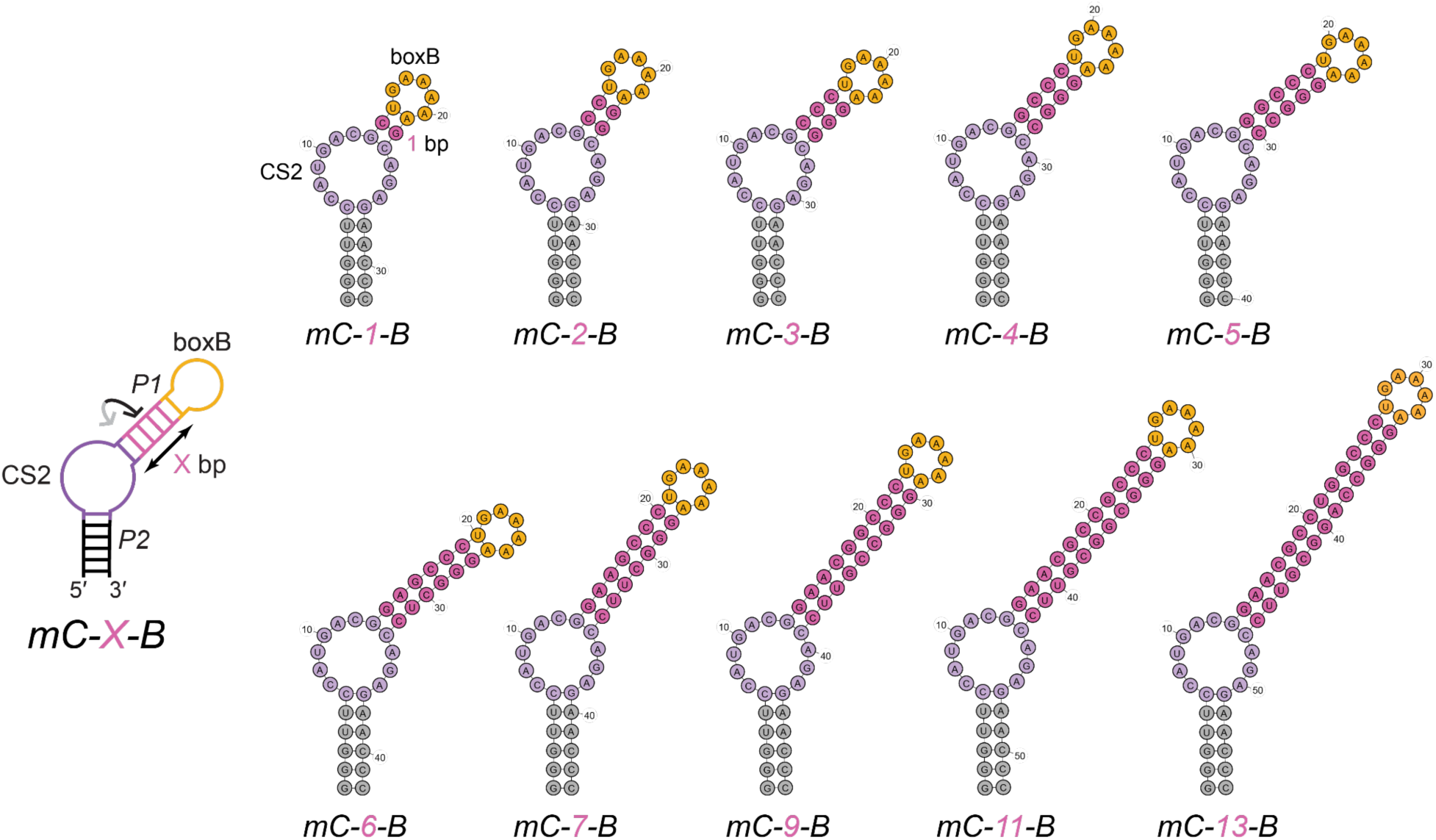
Optimized design of minimized CS2-boxB tags. Optimized tags were designed with a varying number of base pairs (X, pink) that modulate the helical phase and distance between the CS2 k-turn (purple) and boxB (orange) aptamers.

**Figure S7.**
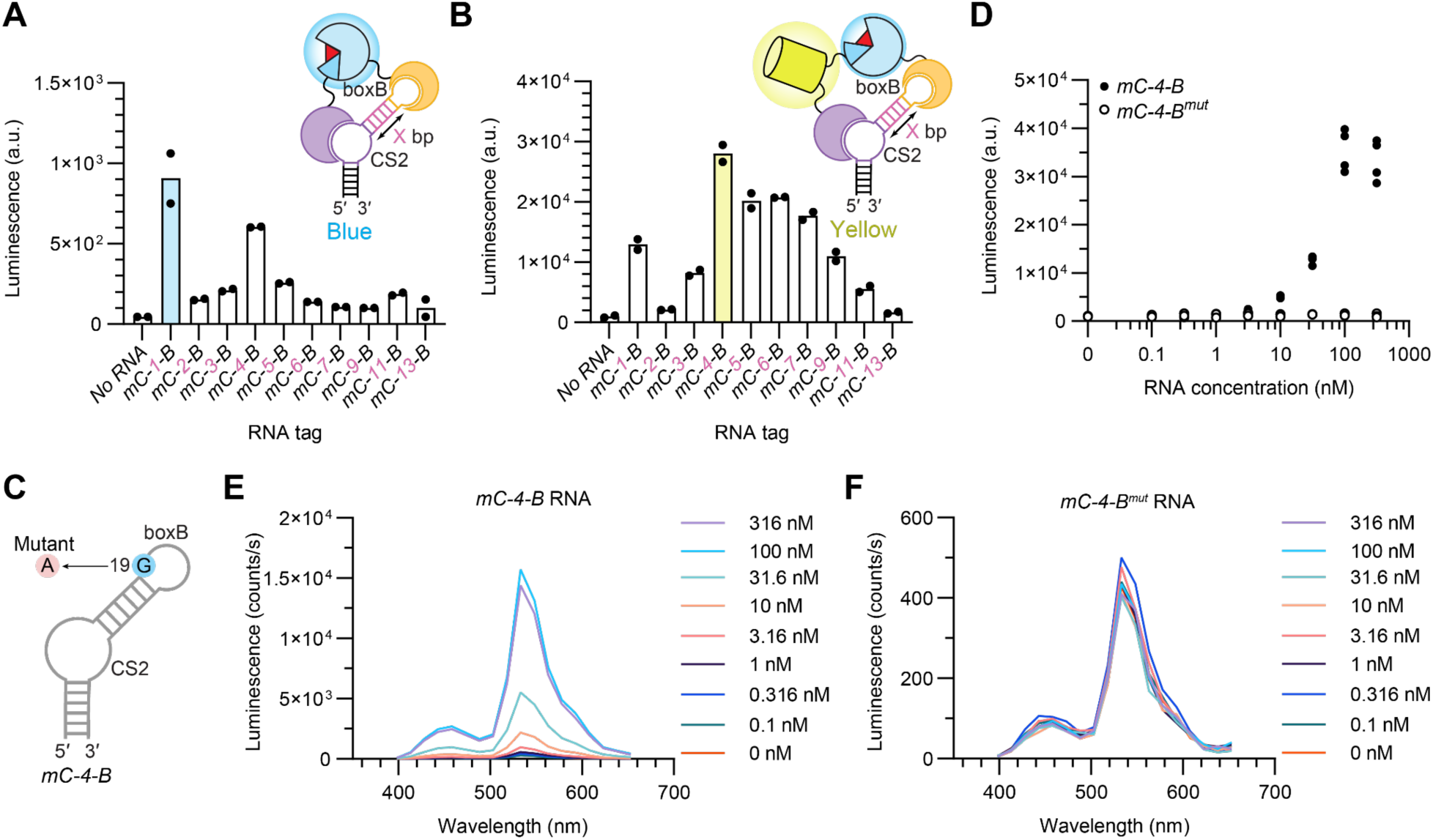
Minimized RNA tag optimization. (**A**) Absolute luminescence values corresponding to luminescence in **Fig. 2B** and fold change plot in **Fig. 2C**. (**B**) Absolute luminescence values corresponding to luminescence image in **Fig. 2E** and fold change plot in **Fig. 2F**. (**C)** Schematic of *mC-4-B^.mut^* tag with a single G19A substitution of the boxB aptamer. (**A, B**) Mean luminescence for each sample (*n* = 2) is shown, with individual data points indicated. (**D**) Total luminescence of *mC-4-B* and *mC-4-B^mut^* titrations with yellow L7Ae-Nun lantern in HEK293T lysate. Individual data points for each condition (*n = 2*) are shown. Luminescence spectra of yellow L7Ae-Nun lantern assembled with varying concentrations of (**E**) *mC-4-B* (0–316 nM) or (**F**) *mC-4-B^mut^*(0–316 nM) corresponding to the fold total luminescence in (D) and fold changes plotted in **Fig. 2G**. Luminescence spectra represent the mean of *n* = 2 replicates. (**E, F**) The spectra of the 100 nM conditions are shown separately in **Fig 2H**. a.u., arbitrary units.

**Figure S8.**
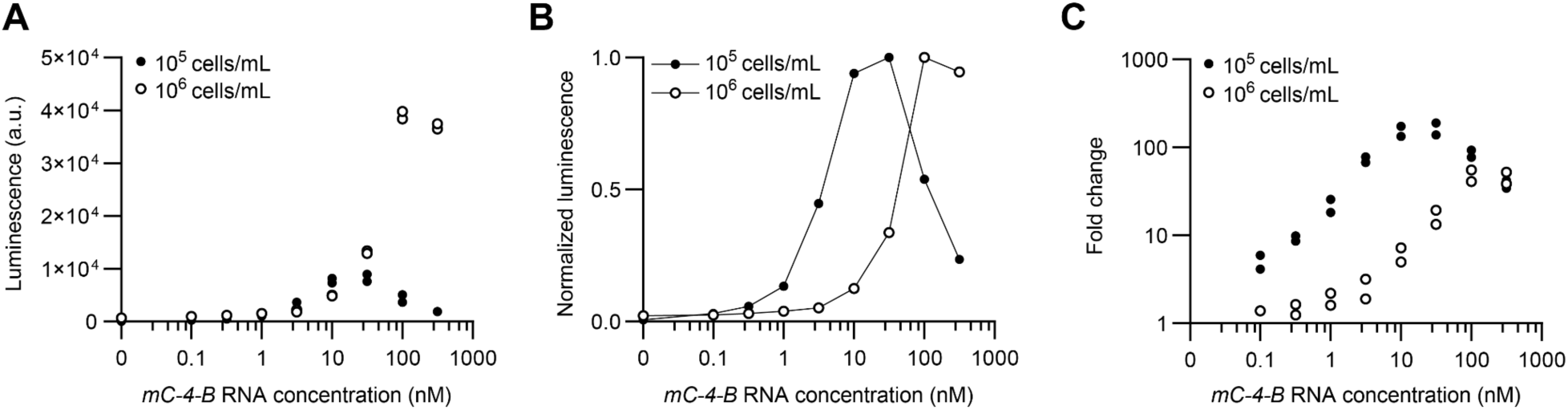
RNA tag sensitivity is dependent on lantern concentration. The L7Ae-Nun BRET lantern was expressed in HEK293T lysate and diluted according to cell count. (**A**) Total luminescence was imaged using a super-cooled EMCCD following addition of increasing concentrations of *mC-4-B* RNA tag (0–316 nM). (**B**) Luminescence intensities were normalized across to the maximum value for each titration to determine the optimal RNA tag concentration. Points represent the mean (*n = 2*). (**C**) Fold-change in luminescence over the no-RNA control (0 nM) for each lantern condition, indicating higher signal-to-background ratio for lower concentrations of the lantern components. An RNA-dependent hook effect was observed under both lysate conditions. (**A, C**) Individual data points (*n* = 2) are shown per condition. a.u., arbitrary units.

**Figure S9.**
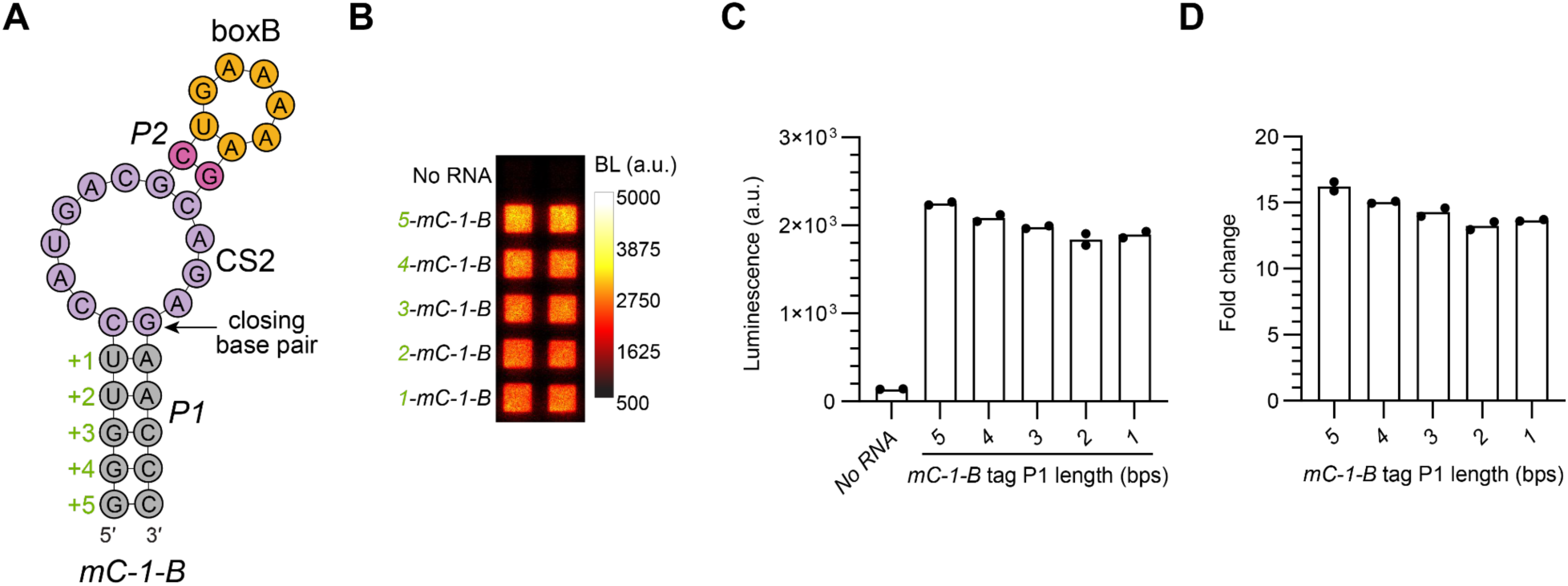
Truncation of the mC-1-B P1 helix defines minimal RNA tag requirements for lantern assembly. The *mC-1-B* RNA tag was systematically truncated within the P1 helix to reduce overall tag size and assess compatibility with the yellow L7Ae-Nun lantern. The lantern was expressed via IVTT and supplemented with 100 nM of the truncated *mC-1-B* variants (P1 helices ranging from 5 to 0 base-pairs). (**A**) Schematic of *mC-1-B* P1 helix truncation variants. The 5-base-pair P1 construct corresponds to the parent tag assessed in **Fig. 2** and **Fig. S7**. (**B**) Luminescence of truncated *mC-1-B* variants. (**C**) Relative luminescence shown in (B). (**D**) Fold change in luminescence over no-RNA control for each variant. Bars represent the mean (*n = 2*) with individual data points shown. BL, luminescence; a.u., arbitrary units.

**Figure S10.**
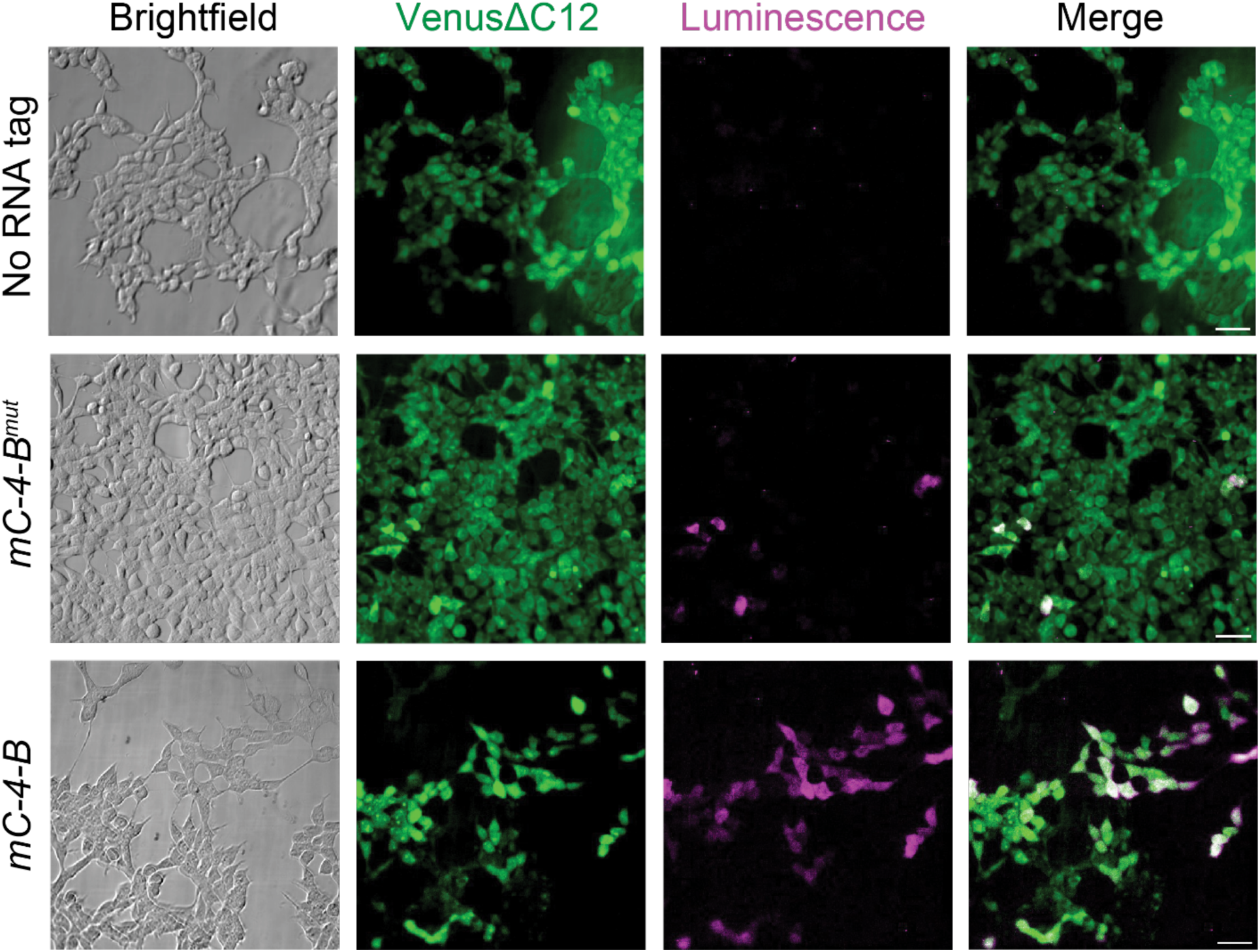
Bioluminescence imaging of minimized RNA tag in live cells. HEK293T cells stably expressing the yellow L7Ae-Nun lantern were transiently transfected with *in vitro* transcribed *mC-4-B* and *mC-4-B^mut^* RNA tags or left untransfected (no RNA). Cells were imaged using an ultra-cooled EMCCD camera. VenusΔC12 fluorescence (green) indicates lantern expression. Bioluminescence (magenta) was observed primarily in samples transfected with the cognate RNA tag. The images shown are separate replicates from those described in **Fig. 3B**. Scale bars = 50 μm.

**Figure S11.**
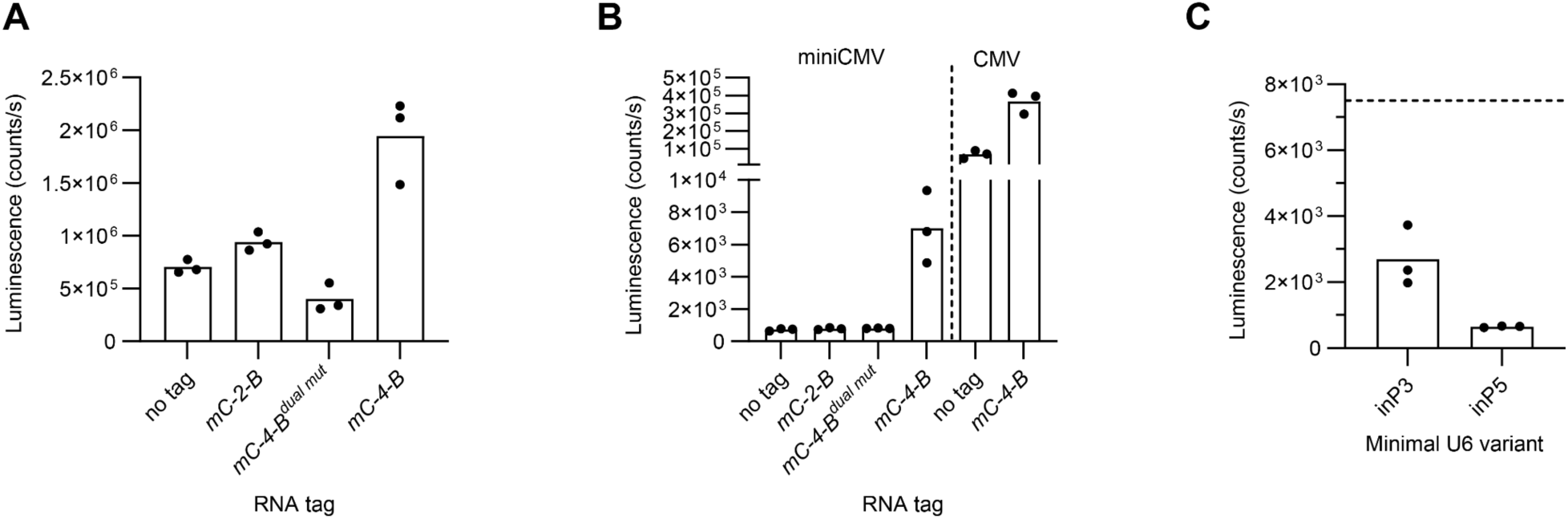
Evaluation of single-vector co-expression of RNA lantern and cognate tag in live cells. (**A, B**) Bulk luminescence corresponding to fold-change data shown in **Figs. 4B** and **4E**, respectively. Wildtype HEK293T cells were transfected with the co-expression vector in which lantern expression was driven either by the (**A**) full-length CMV promoter/enhancer or (**B**) the miniCMV promoter, with or without tag expression via the miniU6 promoter. Inactive tag controls (*mC-2-B and mC-4-B^dual^ ^mut^*) produced luminescence comparable to the no-RNA tag control. (**C**) Bulk luminescence corresponding to fold-change data shown in **Fig. 4G**. Data are derived from the same experiment shown in (B) and share the same controls. Wildtype HEK293T cells were transfected with the co-expression vector in which lantern expression was driven by the miniCMV promoter, and *mC-4-B* expression was modulated using miniU6 variants (inP3 or inP5). The dashed line indicates the bulk luminescence of *mC-4-B* from (B). Bars represent mean values for *n* = 3 replicates, with individual data points shown.

**Figure S12.**
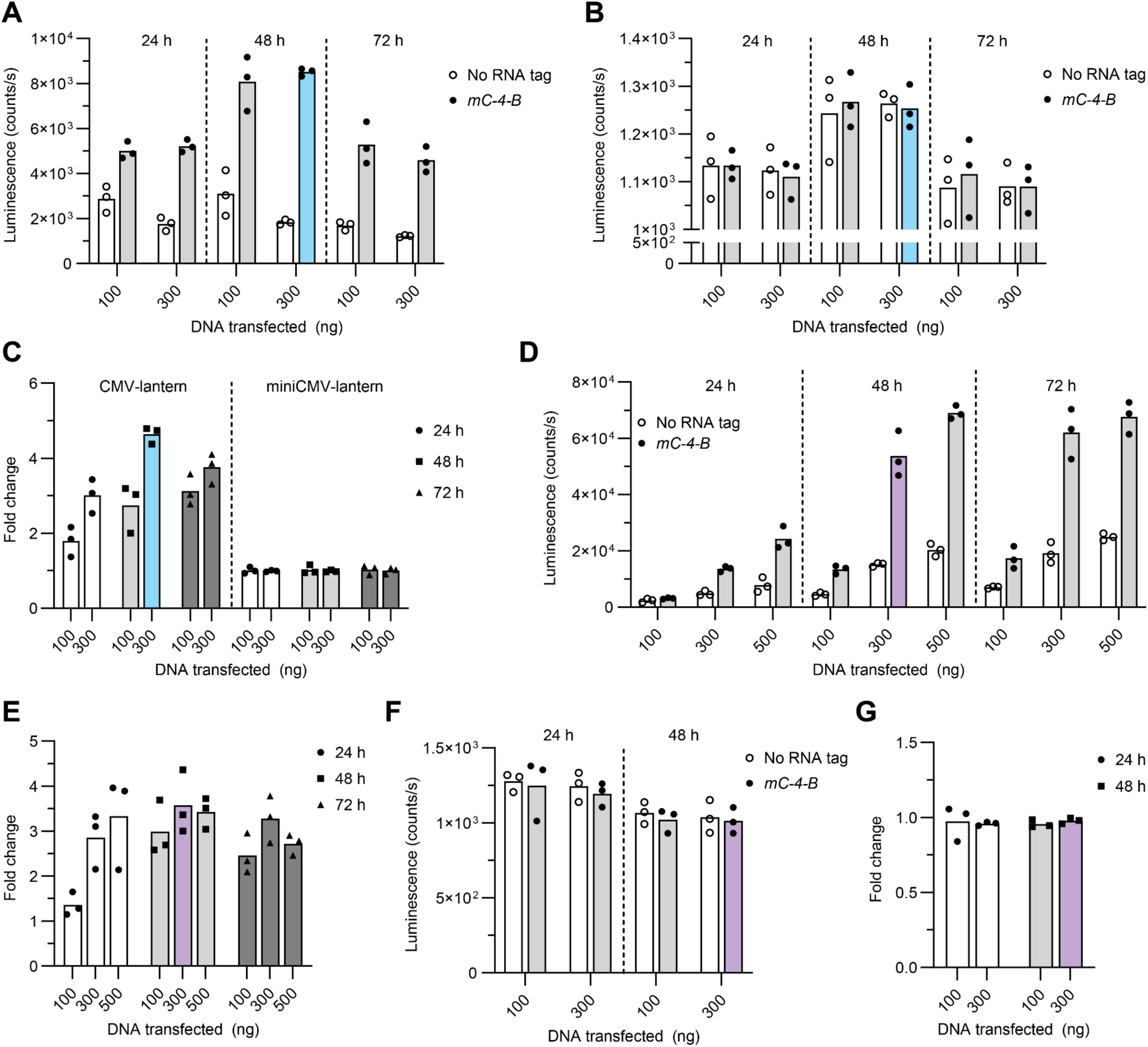
Single-vector co-expression of RNA lantern and cognate tag in U-251 and HepG2 cells. (**A, B**) Bulk luminescence in U-251 glioblastoma cells was measured over 72 h following transfection with 100 or 300 ng of the co-expression plasmid, from which lantern expression was driven either by (**A**) the full-length CMV promoter and enhancer or (**B**) the miniCMV promoter, with or without *mC-4-B* tag expression driven by the miniU6 promoter. U-251 data represent the full time-course from which the optimal condition (blue) was selected. (**C**) Fold-change in bulk luminescence relative to the no-RNA tag control for the data shown in (A, B). (**D**) Bulk luminescence in HepG2 hepatocarcinoma cells was measured over 72 h following transfection with 100, 300, or 500 ng of the co-expression vector, from which lantern expression was driven by the full-length CMV promoter and enhancer, with or without *mC-4-B* tag expression. (**E**) Fold-change in bulk luminescence observed from HepG2 cells relative to the no-RNA tag control for the data shown in (D). (**F**) Bulk luminescence in HepG2 hepatocarcinoma cells was measured over 48 h following transfection with 100 or 300 ng of the co-expression vector, in which lantern expression was driven by the miniCMV, with or without *mC-4-B* tag expression. (**G**) Fold-change in bulk luminescence relative to the no-RNA tag control for the data shown in (F). HepG2 data represent the full time-course from which the optimal condition (highest fold-change; purple) was selected. Bars represent mean values for *n* = 3 replicates, with individual data points shown.

**Figure S13.**
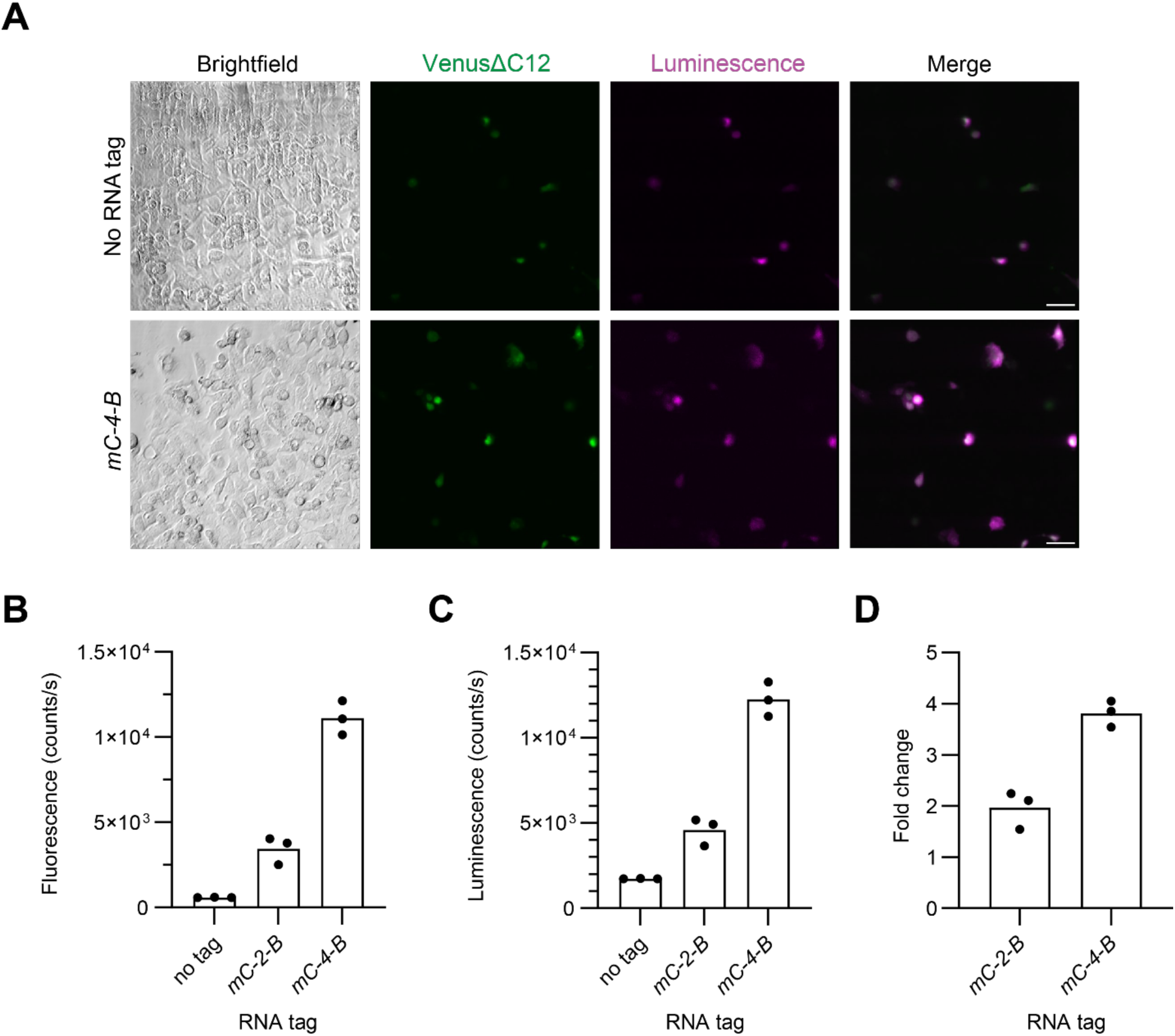
HepG2 bioluminescence imaging with single-vector co-expression of RNA lantern and cognate tag. (**A**) HepG2 hepatocarcinoma cells were transiently transfected with the co-expression vector in which lantern expression was driven either by the full-length CMV promoter/enhancer with or without tag expression via the miniU6 promoter. Cells were imaged using a super-cooled EMCCD camera. VenusΔC12 fluorescence (green) indicates lantern expression. Bioluminescence (magenta) was observed primarily in samples transfected with the *mC-4-B* tag. Scale bars = 50 μm. (**B**) Bulk fluorescence, reporting lantern expression, of HepG2 cells was measured after 72 h following transfection. (**C**) Bulk luminescence of HepG2 cells 72 h post-transfection. (**D**) Fold-change in bulk luminescence relative to the no-RNA tag control for the data shown in (D). Bars represent mean values for *n* = 3 replicates, with individual data points shown.

**Figure S14.**
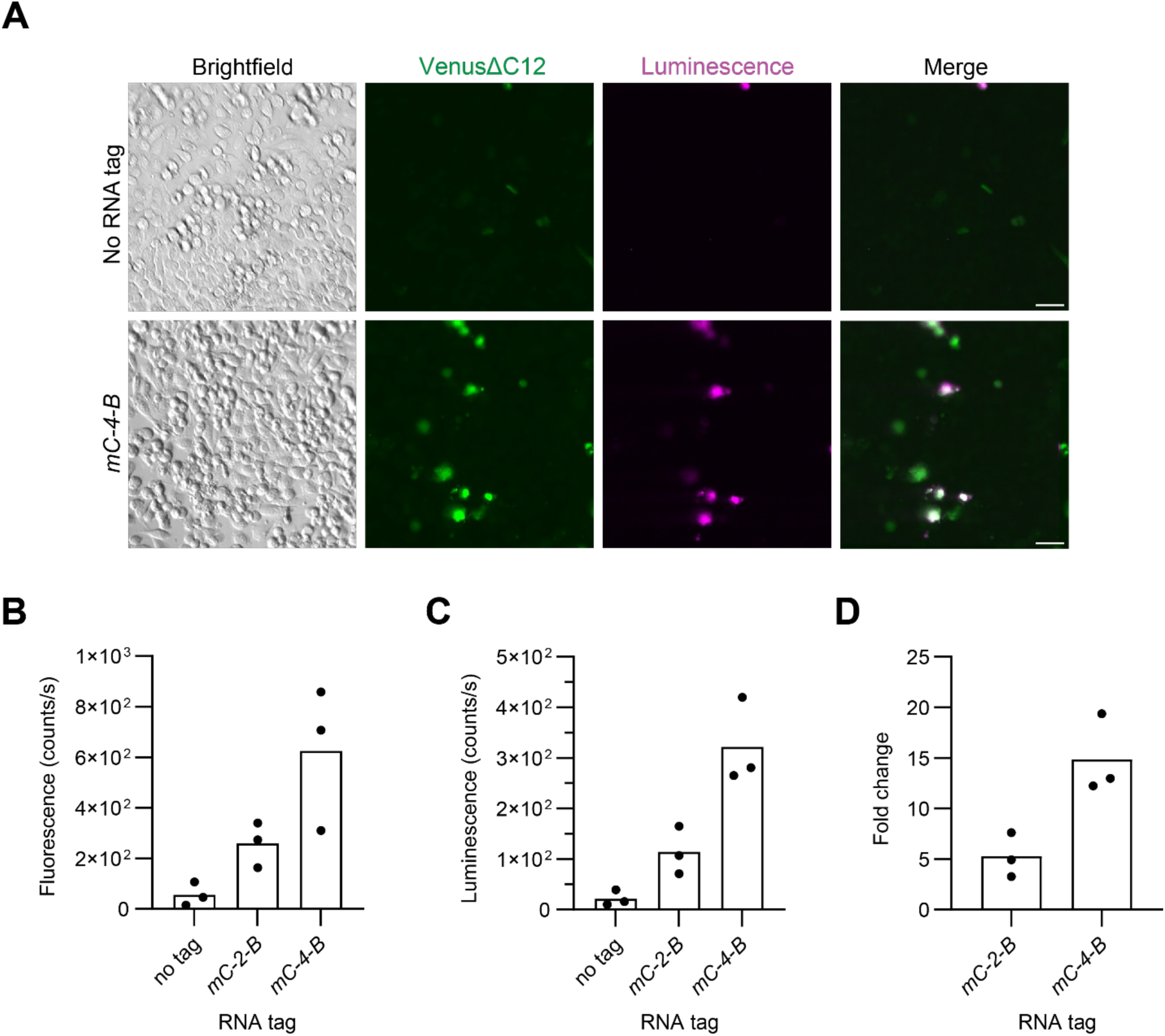
U-251 bioluminescence imaging with single-vector co-expression of RNA lantern and cognate tag. (**A**) U-251 glioblastoma cells were transiently transfected with the co-expression vector in which lantern expression was driven either by the full-length CMV promoter and enhancer with or without tag expression via the miniU6 promoter. Cells were imaged using an ultra-cooled EMCCD camera. VenusΔC12 fluorescence (green) indicates lantern expression. Bioluminescence (magenta) was observed primarily in samples transfected with the *mC-4-B* tag. Scale bars = 50 μm. (**B**) Bulk fluorescence of U-251 cells was measured after 72 h following transfection, corresponding to lantern expression. (**C**) Bulk luminescence of U-251 cells 72 h post-transfection. (**D**) Fold-change in bulk luminescence relative to the no-RNA tag control for the data shown in (D). Bars represent mean values for *n* = 3 replicates, with individual data points indicated.

**Figure S15.**
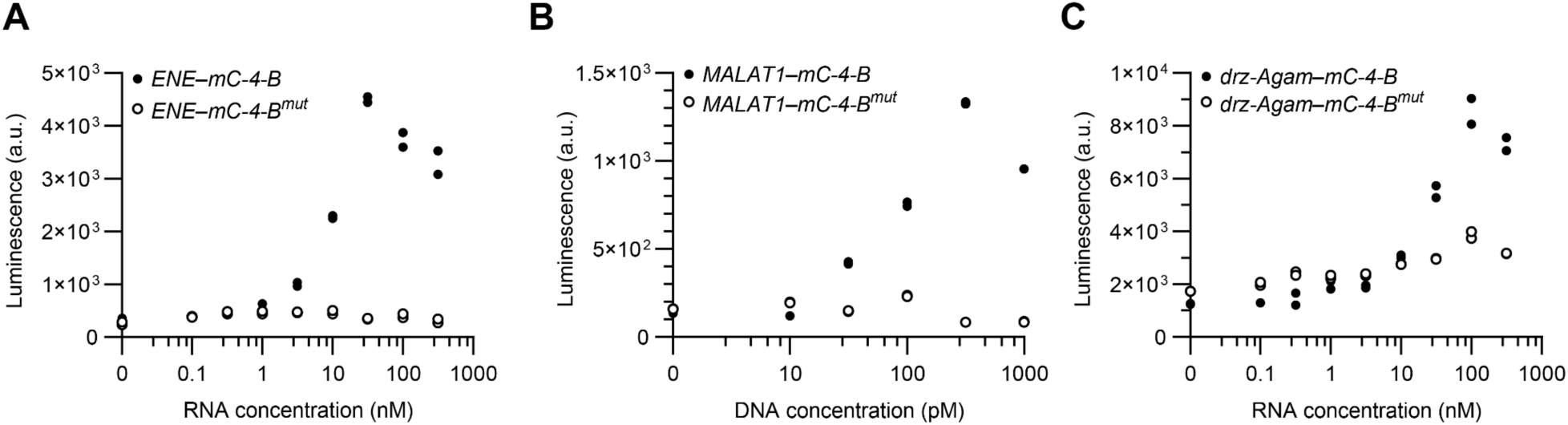
Total luminescence of minimized tag incorporated into highly structured RNAs. (**A**) Total luminescence from titration of *ENE–mC-4-B* and *ENE–mC-4-B^mut^*in HEK293T lysate containing the yellow L7Ae-Nun lantern, corresponding to the fold-change data shown in **Fig. 5C**. (**B**) Total luminescence from titration of *MALAT1–mC-4-B* and *MALAT1–mC-4-B^mut^*evaluated with the yellow L7Ae-Nun lantern via *in vitro* transcription and translation (IVTT). Data correspond to the fold-change plot shown in **Fig. 5F**. (**C**) Total luminescence from titration of *drz-Agam–mC-4-B* and *drz-Agam–mC-4-B^mut^* in HEK293T lysate containing the yellow L7Ae-Nun lantern, corresponding to the fold-change data shown in **Fig. 5I**. Individual data points (*n* = 2) are shown per condition. a.u., arbitrary units.

**Figure S16.**
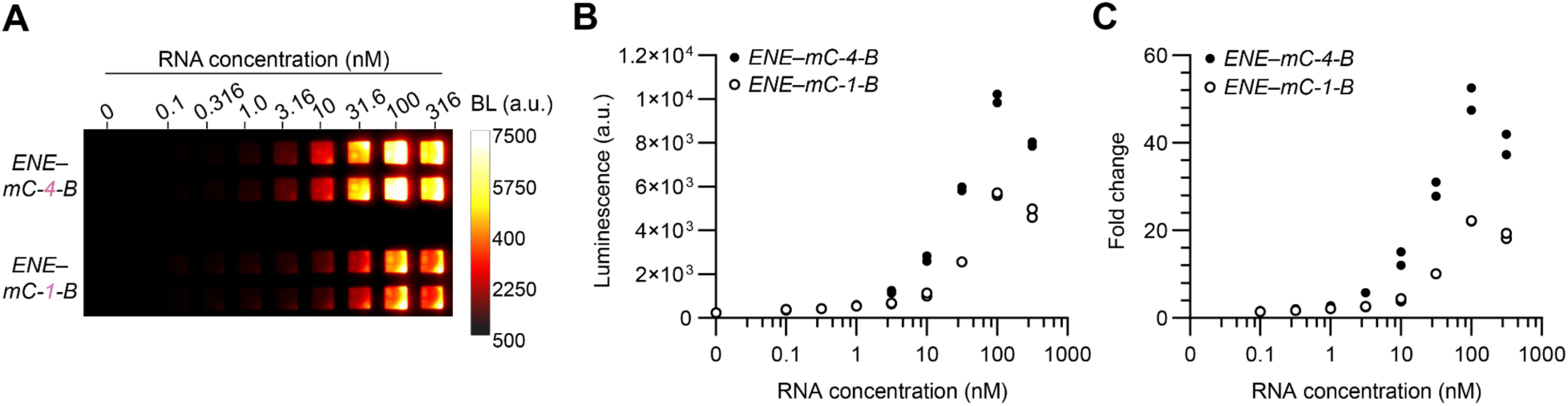
Evaluation of different minimized tags in *MALAT1 ENE* domain. Each component of the yellow L7Ae–Nun BRET lantern was expressed separately in HEK293T cell lysate, combined in equal proportions, and supplemented with varying concentrations (0–316 nM) of ENE-tagged RNA constructs (*ENE–mC-4-B* and *ENE–mC-1-B*). (**A**) Luminescence of *ENE– mC-4-B* compared to *ENE–mC-1-B*, imaged using a ultra-cooled EMCCD camera. (**B**) Total luminescence of *ENE–mC-4-B* and *ENE–mC-1-B* across RNA concentrations. (**C**) Fold change in luminescence relative to the no-RNA control (0 nM) for each construct across concentrations. Individual data points (*n* = 2) are shown per condition. BL, luminescence; a.u., arbitrary units.

**Figure S17.**
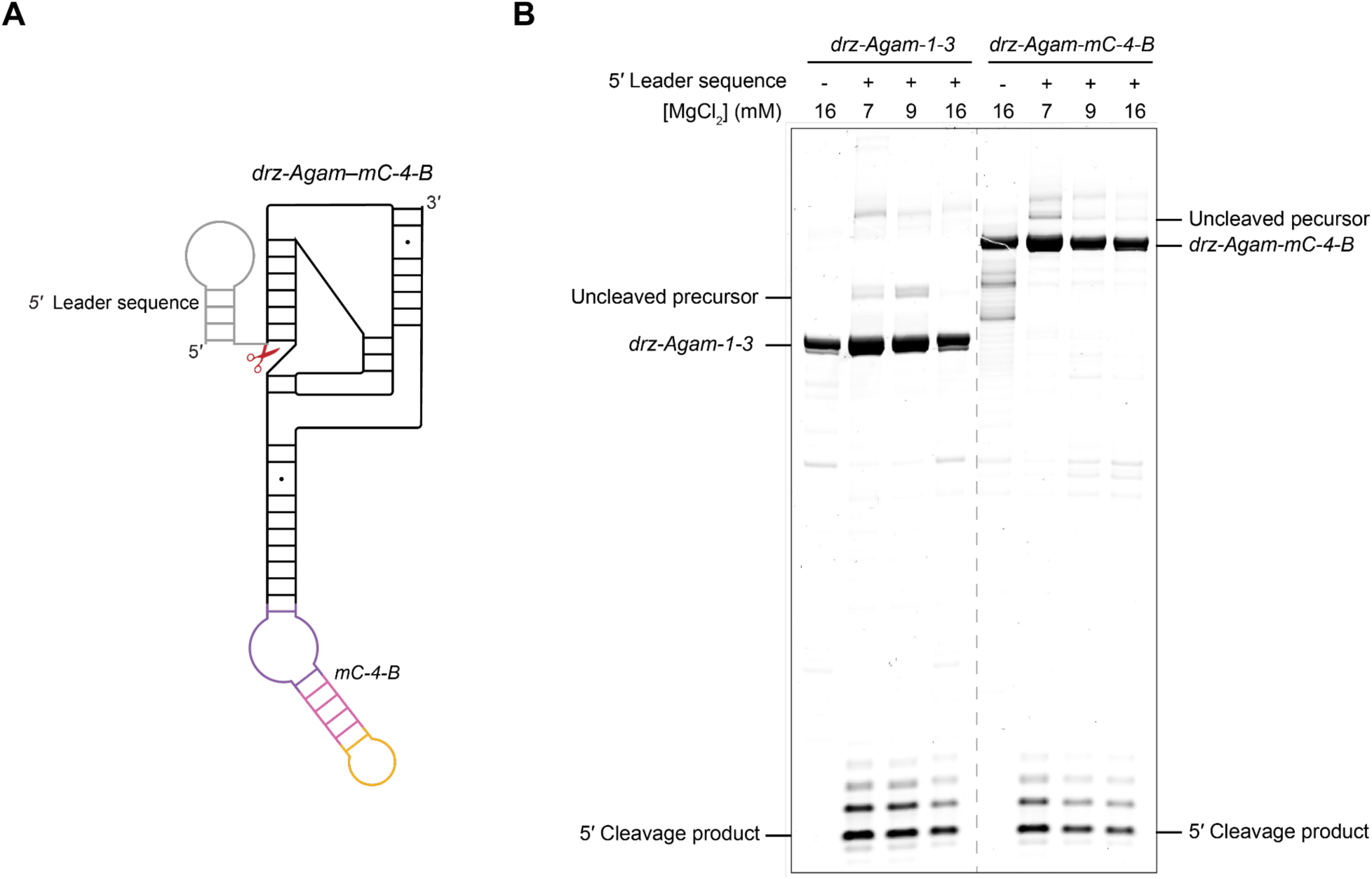
Evaluation of catalytic activity of a tagged HDV-like ribozyme. Transcripts containing a short 5′ leader sequence (22 nucleotides) upstream of tagged and untagged variants of *drz-Agam-1-3* were generated by *in vitro* transcription to enable detection of cleavage products. Reactions were performed across a range of MgCl_2_ concentrations (7, 9, and 16 mM) in the presence of 8 mM rNTPs to probe suboptimal, near-optimal, and high-activity conditions. Cleavage products were analyzed by denaturing PAGE. (**A**) Schematic of *drz-Agam–mC-4-B* ribozyme with a structured 5′ leader sequence (a Ni-NTA aptamer) used to enable detection of cleavage products by intercalating fluorophores. (**B**) Denaturing PAGE analysis showing cleavage of wild-type and tagged *drz-Agam–mC-4-B* constructs across MgCl_2_ conditions. Accumulation of uncleaved precursor RNA at lower Mg^2+^ ion concentrations serve as a reference for ribozyme-mediated cleavage, demonstrating that incorporation of the minimized RNA tag does not disrupt catalytic activity.

**Figure S18.**
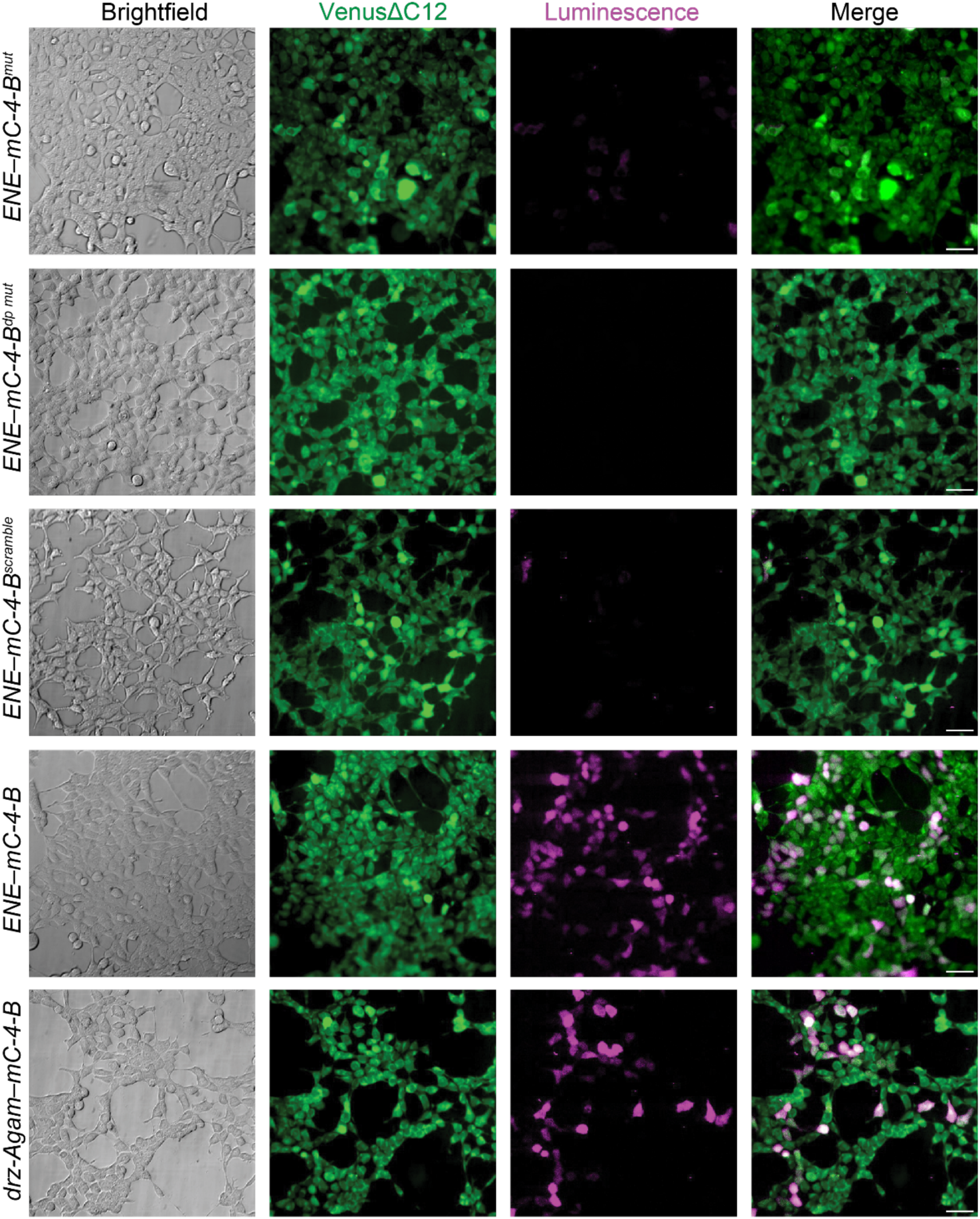
Bioluminescence imaging of highly structured RNAs with internal minimized RNA tag in live cells. HEK293T cells stably expressing the yellow L7Ae-Nun Lantern were transfected with *in vitro* transcribed *ENE–mC-4-B*, mutant variants (*ENE–mC-4-B^mut^*, *ENE–mC-4-B^scramble^*, and *ENE–mC-4-B^dp^ ^mut^*), or *drz-Agam–mC-4-B*. Cells were imaged using a super-cooled EMCCD camera. VenusΔC12 fluorescence (green) indicates lantern expression, and bioluminescence (magenta) reflects RNA-dependent lantern assembly. Images for *ENE–mC-4-B*, *ENE–mC-4-B^mut^*, and *drz-Agam–mC-4-B* are replicates that correspond to **Fig. 6B**. Scale bars = 50 μm.

**Table S1.**
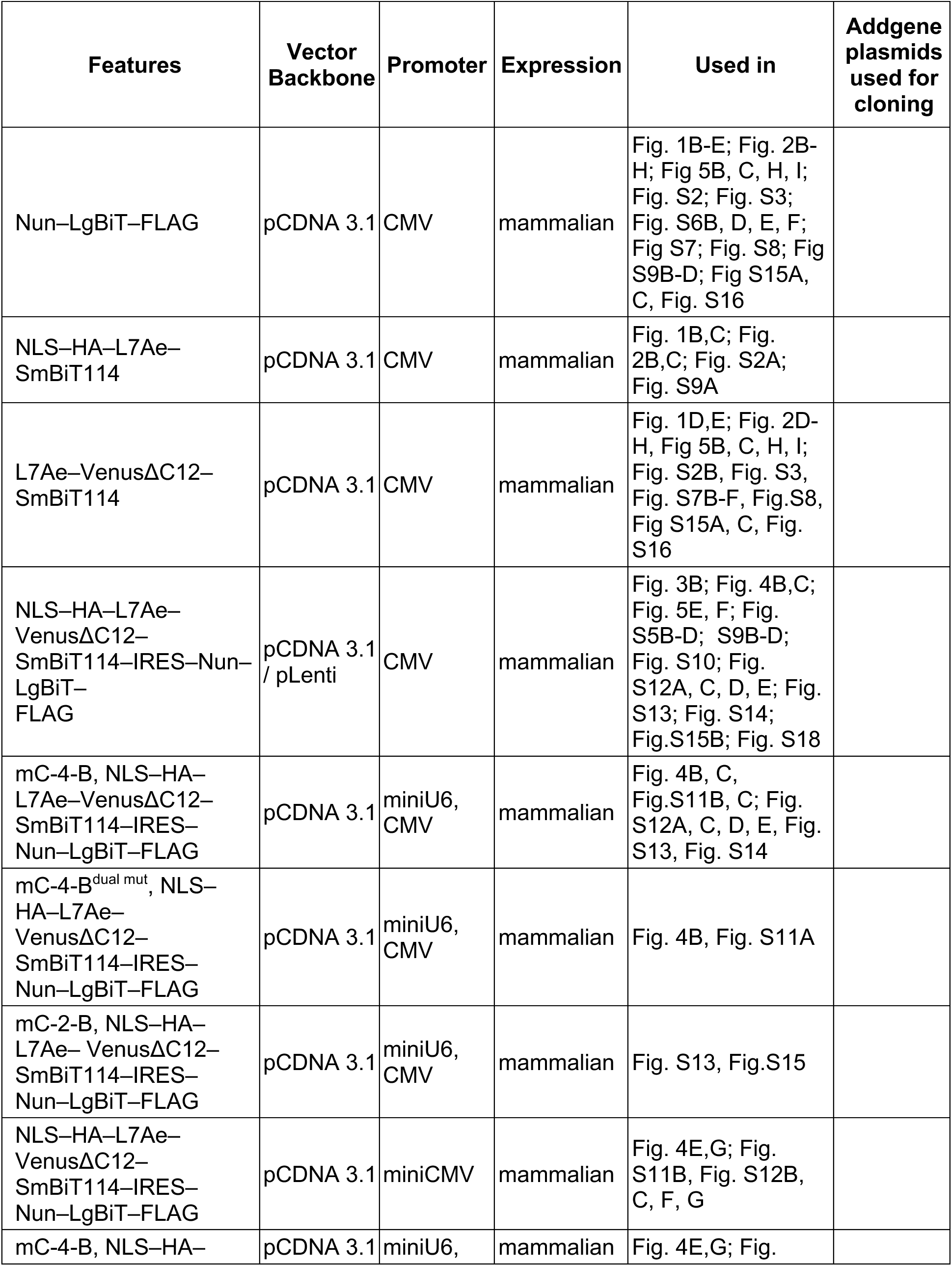
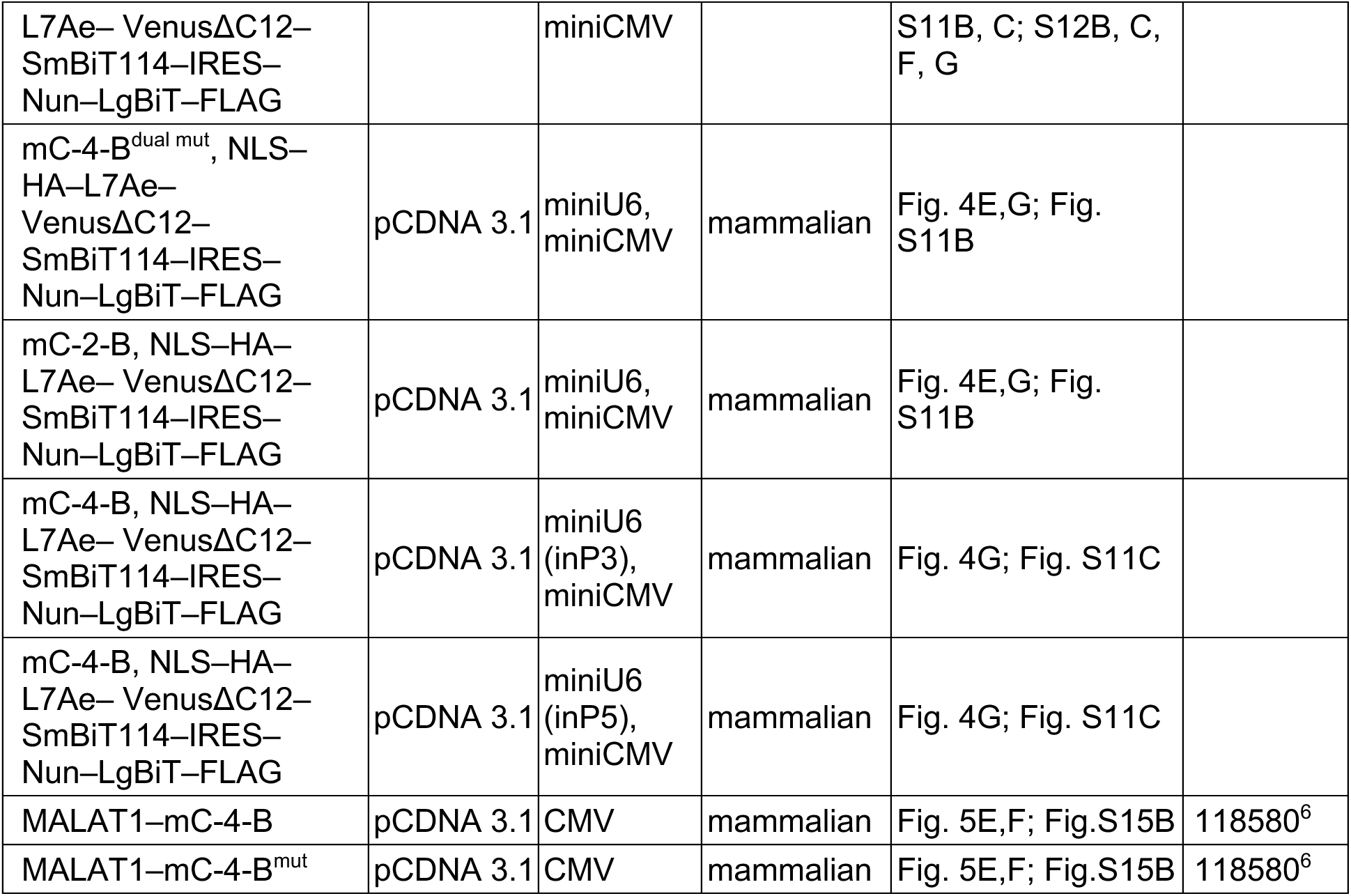

### List S1. Plasmid DNA and RNA tag sequences

*mini-boxB-1-CS2 (mB-1-C):*

ggaugcagagcugaaaaagccaugacgcaucc

*mini-boxB-2-CS2 (mB-2-C):*

ggaugcagagacugaaaaaguccaugacgcaucc

*mini-boxB-3-CS2 (mB-3-C):*

ggaugcagagaacugaaaaaguuccaugacgcaucc

*mini-boxB-4-CS2 (mB-4-C):*

ggaugcagagaaccugaaaaagguuccaugacgcaucc

*mini-boxB-5-CS2 (mB-5-C):*

ggaugcagagaacccugaaaaaggguuccaugacgcaucc

*mini-boxB-7-CS2 (mB-7-C):*

ggaugcagagaacgcccugaaaaagggcguuccaugacgcaucc

*mini-boxB-9-CS2 (mB-9-C):*

ggaugcagagaaccugcccugaaaaagggcagguuccaugacgcaucc

*mini-boxB-11-CS2 (mB-11-C):*

ggaugcagagaaccuaggcccugaaaaagggccuagguuccaugacgcaucc

*mini-boxB-13-CS2 (mB-13-C):*

ggaugcagagaaccuagcugcccugaaaaagggcagcuagguuccaugacgcaucc

*mini-boxB-15-CS2 (mB-15-C):*

ggaugcagagaaccuagcucagcccugaaaaagggcugagcuagguuccaugacgcaucc

*mini-boxB-3-CS2-bulge_1 (mB-3-C_b1):*

ggaugcagagaaacugaaaaaguuccaugacgcaucc

*mini-boxB-3-CS2-bulge_2 (mB-1-C_b2):*

ggaugcagagaacugaaaaaguauccaugacgcaucc

*mini-boxB-3-CS2-bulge_3 (mB-3-C_b3):*

ggaugcagagaacugaaaaagauuccaugacgcaucc

*mini-CS2-4-boxB^mut^ (mC-4-B^mut^):*

gguuccaugacggcccuaaaaaagggccagagaaccc

*mini-CS2-1-boxB (mC-1-B):*

gguuccaugacgcugaaaaagcagagaaccc

*mini-CS2-2-boxB (mC-2-B):*

gguuccaugacgccugaaaaaggcagagaaccc

*mini-CS2-3-boxB (mC-3-B):*

gguuccaugacgcccugaaaaagggcagagaaccc

*mini-CS2-4-boxB (mC-4-B):*

gguuccaugacggcccugaaaaagggccagagaaccc

*mini-CS2-5-boxB (mC-5-B):*

gguuccaugacgggcccugaaaaagggcccagagaaccc

*mini-CS2-6-boxB (mC-6-B):*

gguuccaugacggagcccugaaaaagggcuccagagaaccc

*mini-CS2-7-boxB (mC-7-B):*

gguuccaugacggaagcccugaaaaagggcuuccagagaaccc

*mini-CS2-9-boxB (mC-9-B):*

gguuccaugacggaacggcccugaaaaagggccguuccagagaaccc

*mini-CS2-11-boxB (mC-11-B):*

gguuccaugacggaacgccgcccugaaaaagggcggcguuccagagaaccc

*mini-CS2-13-boxB (mC-13-B):*

gguuccaugacggaacgccuggcccugaaaaagggccaggcguuccagagaaccc

*5-mini-CS2-1-boxB (5-mC-1-B):*

gguuccaugacgcugaaaaagcagagaaccc

*4-mini-CS2-1-boxB (4-mC-1-B):*

guuccaugacgcugaaaaagcagagaacu

*3-mini-CS2-1-boxB (3-mC-1-B):*

gguccaugacgcugaaaaagcagagacu

*2-mini-CS2-1-boxB (2-mC-1-B):*

ggccaugacgcugaaaaagcagagcu

*1-mini-CS2-1-boxB (1-mC-1-B):*

gccaugacgcugaaaaagcagagc

*MALAT1-ENE –mC-4-B:*

ggaagguuuuucuuuuccugagaccaugacggcccugaaaaagggccagagucucagguuuugcuuuuuggccuuuuucuagcuuaaaaaaaaaaaaagcaaaa

*MALAT1-ENE –mC-4-B^mut^:*

ggaagguuuuucuuuuccugagaccaugacggcccuaaaaaagggccagagucucagguuuugcuuuuuggccuuuuucuagcuuaaaaaaaaaaaaagcaaaa

*MALAT1-ENE –mC-4-B^scramble^:* ggaagguuuuucuuuuccugagaccaugacggcccuuacauagggccagagucucagguuuugcuuuuuggccuuuuucuagcuuaaaaaaaaaaaaagcaaaa

*MALAT1-ENE –mC-4-B^dp^ ^mut^:*

ggaagguuuuucuuuuccugagaccaugacggcccuaaaaaagggccagcgucucagguuuugcuuuuuggccuuuuucuagcuuaaaaaaaaaaaaagcaaaa

*5’ leader sequence for drz-Agam-1-3 and drz-Agam–mC-4-B: gggcaauugaaaaaacgcccau drz-Agam-1-3:*

ggctgacaaaatcctatctctacctcctcgtggtgtcagccggaatgtgcagtatcagtactgcatataccaacagagagggucag

*drz-Agam–mC-4-B:* ggcugacaaaauccuaucucuaccuccucguggugucagccggaaugugcaguaccatgacggcccugaaaaagggccagaguacugcauauaccaacagagagggucag

*drz-Agam–mC-4-B^mut^:* ggcugacaaaauccuaucucuaccuccucguggugucagccggaaugugcaguaccatgacggcccuaaaaaagggccagaguacugcauauaccaacagagagggucag

#### Nun–LgBiT–FLAG

atgggcatggatcgcgggcttacgtcgcgtgaccgccgccgtatcgcgcgttgggaaaagcgcatcgcttacgccctgaagaacggagcggatccaccggtatgcatcgccaccgccttcacactcgaagatttcgttggggactgggaacagacagccgcctacaacctggaccaagtccttgaacagggaggtgtgtccagtttgctgcagaatctcgccgtgtccgtaactccgatccaaaggattgtccggagcggtgaaaatgccctgaagatcgacatccatgtcatcatcccgtatgaaggtctgagcgccgaccaaatggcccagatcgaagaggtgtttaaggtggtgtaccctgtggatgatcatcactttaaggtgatcctgccctatggcacactggtaatcgacggggttacgccgaacatgctgaactatttcggacggccgtatgaaggcatcgccgtgttcgacggcaaaaagatcactgtaacagggaccctgtggaacggcaacaaaattatcgacgagcgcctgatcacccccgacggctccatgctgttccgagtaaccatcaacagcctcgaggactacaaggacgacgatgacaagtaa

#### NLS–HA–L7Ae–SmBiT

atgggcccaaaaaagaaaagaaaagttggctacccctacgacgtgcccgactacgccatcgaaggccgccatatgctagccgttaaaatgtacgtgagatttgaggttcctgaggacatgcagaacgaagctctgagtctgctggagaaggttagggagagcggtaaggtaaagaaaggtaccaacgagacgacaaaggctgtggagaggggactggcaaagctcgtttacatcgcagaggatgttgacccgcctgagatcgttgctcatctgcccctcctctgcgaggagaagaatgtgccgtacatttacgttaaaagcaagaacgaccttggaagggctgtgggcattgaggtgccatgcgcttcggcagcgataatcaacgagggagagctgagaaaggagcttggaagccttgtggagaagattaaaggccttcagaaggcggatccaccggtatgcatcgccaccgccgtgaccggctaccgcctgtttgaggagattctg

#### L7Ae–VenusΔC12–SmBiT

atgggctacgtgagatttgaggttcctgaggacatgcagaacgaagctctgagtctgctggagaaggttagggagagcggtaaggtaaagaaaggtaccaacgagacgacaaaggctgtggagaggggactggcaaagctcgtttacatcgcagaggatgttgacccgcctgagatcgttgctcatctgcccctcctctgcgaggagaagaatgtgccgtacatttacgttaaaagcaagaacgaccttggaagggctgtgggcattgaggtgccatgcgcttcggcagcgataatcaacgagggagagctgagaaaggagcttggaagccttgtggagaagattaaaggccttcagaaggcggatccaccggtatgcatcgccaccgccgtgagcaagggcgaggagctgttcaccggggtggtgcccatcctggtcgagctggacggcgacgtaaacggccacaagttcagcgtgtccggcgagggcgagggcgatgccacctacggcaagctgaccctgaagctgatctgcaccaccggcaagctgcccgtgccctggcccaccctcgtgaccaccctgggctacggcctgcagtgcttcgcccgctaccccgaccacatgaagcagcacgacttcttcaagtccgccatgcccgaaggctacgtccaggagcgcaccatcttcttcaaggacgacggcaactacaagacccgcgccgaggtgaagttcgagggcgacaccctggtgaaccgcatcgagctgaagggcatcgacttcaaggaggacggcaacatcctggggcacaagctggagtacaactacaacagccacaacgtctatatcaccgccgacaagcagaagaacggcatcaaggccaacttcaagatccgccacaacatcgaggacggcggcgtgcagctcgccgaccactaccag cagaacacccccatcggcgacggccccgtgctgctgcccgacaaccactacctgagctaccagtccgccctgagcaaagaccccaacgagaagcgcgatcacatggtcctgctggagttcgtgaccgccgcgatggtgaccggctaccgcctgtttgaggagattctg

#### NLS–HA–L7Ae–VenusΔC12–SmBiT–IRES–Nun–LgBiT–FLAG

atgggcccaaaaaagaaaagaaaagttggctacccctacgacgtgcccgactacgccatcgaaggccgccatatgctagccgttaaaatgatgggctacgtgagatttgaggttcctgaggacatgcagaacgaagctctgagtctgctggagaaggttagggagagcggtaaggtaaagaaaggtaccaacgagacgacaaaggctgtggagaggggactggcaaagctcgtttacatcgcagaggatgttgacccgcctgagatcgttgctcatctgcccctcctctgcgaggagaagaatgtgccgtacatttacgttaaaagcaagaacgaccttggaagggctgtgggcattgaggtgccatgcgcttcggcagcgataatcaacgagggagagctgagaaaggagcttggaagccttgtggagaagattaaaggccttcagaaggcggatccaccggtatgcatcgccaccgccgtgagcaagggcgaggagctgttcaccggggtggtgcccatcctggtcgagctggacggcgacgtaaacggccacaagttcagcgtgtccggcgagggcgagggcgatgccacctacggcaagctgaccctgaagctgatctgcaccaccggcaagctgcccgtgccctggcccaccctcgtgaccaccctgggctacggcctgcagtgcttcgcccgctaccccgaccacatgaagcagcacgacttcttcaagtccgccatgcccgaaggctacgtccaggagcgcaccatcttcttcaaggacgacggcaactacaagacccgcgccgaggtgaagttcgagggcgacaccctggtgaaccgcatcgagctgaagggcatcgacttcaaggaggacggcaacatcctggggcacaagctggagtacaactacaacagccacaacgtctatatcaccgccgacaagcagaagaacggcatcaaggccaacttcaagatccgccacaacatcgaggacggcggcgtgcagctcgccgaccactaccagcagaacacccccatcggcgacggccccgtgctgctgcccgacaaccactacctgagctaccagtccgccctgagcaaagaccccaacgagaagcgcgatcacatggtcctgctggagttcgtgaccgccggcggcggaggctctgtgaccggctaccgcctgtttgaggagattctgtaacgtacgacgcgtggatccctcccccccccctaacgttactggccgaagccgcttggaataaggccggtgtgcgtttgtctatatgttattttccaccatattgccgtcttttggcaatgtgagggcccggaaacctggccctgtcttcttgacgagcattcctaggggtctttcccctctcgccaaaggaatgcaaggtctgttgaatgtcgtgaaggaagcagttcctctggaagcttcttgaagacaaacaacgtctgtagcgaccctttgcaggcagcggaaccccccacctggcgacaggtgcctctgcggccaaaagccacgtgtataagatacacctgcaaaggcggcacaaccccagtgccacgttgtgagttggatagttgtggaaagagtcaaatggctctcctcaagcgtattcaacaaggggctgaaggatgcccagaaggtaccccattgtatgggatctgatctggggcctcggtgcacatgctttacatgtgtttagtcgaggttaaaaaaacgtctaggccccccgaaccacggggacgtggttttcctttgaaaaacacgatgataatatggccacagcctgcaggatggatcgcgggcttacgtcgcgtgaccgccgccgtatcgcgcgttgggaaaagcgcatcgcttacgccctgaagaacggagcggatccaccggtatgcatcgccaccgccttcacactcgaagatttcgttggggactgggaacagacagccgcctacaacctggaccaagtccttgaacagggaggtgtgtccagtttgctgcagaatctcgccgtgtccgtaactccgatccaaaggattgtccggagcggtgaaaatgccctgaagatcgacatccatgtcatcatcccgtatgaaggtctgagcgccgaccaaatggcccagatcgaagaggtgtttaaggtggtgtaccctgtggatgatcatcactttaaggtgatcctgccctatggcacactggtaatcgacggggttacgccgaacatgctgaactatttcggacggccgtatgaaggcatcgccgtgttcgacggcaaaaagatcactgtaacagggaccctgtggaacggcaacaaaattatcgacgagcgcctgatcacccccgacggctccatgctgttccgagtaaccatcaacagcctcgaggactacaaggacgacgatgacaag

#### MALAT1–mC-4-B

gtaaaggactggggccccgcaactggcctctcctgccctcttaagcgcagcgccattttagcaacgcagaagcccggcgccgggaagcctcagctcgcctgaaggcaggtcccctctgacgcctccgggagcccaggtttcccagagtccttgggacgcagcgacgagttgtgctgctatcttagctgtccttataggctggccattccaggtggtggtatttagataaaaccactcaaactctgcagtttggtcttggggtttggaggaaagcttttatttttcttcctgctccggttcagaaggtctgaagctcatacctaaccgggcataacacagaatctgcaaaacaaaaacccctaaaaaagcagacccagagcagtgtaaacacttctgggtgtgtccctgactggctgcccaaggtctctgtgtcttcggagacaaagccattcgcttagttggtctactttaaaaggccacttgaactcgctttccatggcgatttgccttgtgagcactttcaggagagcctggaagctgaaaaacggtagaaaaatttccgtgcgggccgtggggggctggcggcaactggggggccgcagatcagagtgggccactggcagccaacggcccccggggctcaggcggggagcagctctgtggtgtgggattgaggcgttttccaagagtgggttttcacgtttctaagatttcccaagcagacagcccgtgctgctccgatttctcgaacaaaaaagcaaaacgtgtggctgtcttgggagcaagtcgcaggactgcaagcagttgggggagaaagtccgccattttgccacttctcaaccgtccctgcaaggctggggctcagttgcgtaatggaaagtaaagccctgaactatcacactttaatcttccttcaaaaggtggtaaactatacctactgtccctcaagagaacacaagtgctttaagaggtattttaaaagttccgggggttttgtgaggtgtttgatgacccgtttaaaatatgatttccatgtttcttttgtctaaagtttgcagctcaaatctttccacacgctagtaatttaagtatttctgcatgtgtagtttgcattcaagttccataagctgttaagaaaaatctagaaaagtaaaactagaacctatttttaaccgaagaactactttttgcctccctcacaaaggcggcggaaggtgatcgaattccggtgatgcgagttgttctccgtctataaatacgcctcgcccgagctgtgcggtaggcattgaggcagccagcgcaggggcttctgctgagggggcaggcggagcttgaggaaacc gcagatgagtttttttctctttgaaagatagagattaatacaactacttaaaaaatatagtcaataggttactaagatattgcttagcgttaagtttttaacgtaattttaatagcttaagattttaagagaaaatatgaagacttagaagagtagcatgaggaaggaaaagataaaaggtttctaaaacatgacggaggttgagatgaagcttcttcatggagtaaaaaatgtatttaaaagaaaattgagagaaaggactacagagccccgaattaataccaatagaagggcaatgcttttagattaaaatgaaggtgacttaaacagcttaaagtttagtttaaaagttgtaggtgattaaaataatttgaaggcgatcttttaaaaagagattaaactgaaggtgattaaaagaccttgaaatccatgacgcagggagaattgcgtcatttaaagcctagttaacgcatttactaaacgcagacgaaaatggaaagattaattgggagtggtaggatgaaacaatttggagaagatagaagtttgaagtggaaaactggaagacagaagtacgggaaggcgaagaaaagaatagagaagatagggaaattagaagataaaaacatacttttagaagaaaaaagataaatttaaacctgaaaagtaggaagcagaagaaaaaagacaagctaggaaacaaaaagctaagggcaaaatgtacaaacttagaagaaaattggaagatagaaacaagatagaaaatgaaaatattgtcaagagtttcagatagaaaatgaaaaacaagctaagacaagtattggagaagtatagaagatagaaaaatataaagccaaaaattggataaaatagcactgaaaaaatgaggaaattattggtaaccaatttattttaaaagcccatcaatttaatttctggtggtgcagaagttagaaggtaaagcttgagaagatgagggtgtttacgtagaccagaaccaatttagaagaatacttgaagctagaaggggaagttggttaaaaatcacatcaaaaagctactaaaaggactggtgtaatttaaaaaaaactaaggcagaaggcttttggaagagttagaagaatttggaaggccttaaatatagtagcttagtttgaaaaatgtgaaggactttcgtaacggaagtaattcaagatcaagagtaattaccaacttaatgtttttgcattggactttgagttaagattattttttaaatcctgaggactagcattaattgacagctgacccaggtgctacacagaagtggattcagtgagtctaggaagacagcagcagacaggattccaggaaccagtgtttgatgaagctaggactgaggagcaagcgagcaagcagcagttcgtggtgaagataggaaaagagtccaggagccagtgcgatttggtgaaggaagctaggaagaaggaaggagcgctaacgatttggtggtgaagctaggaaaaaggattccaggaaggagcgagtgcaatttggtgatgaaggtagcaggcggcttggcttggcaaccacacggaggaggcgagcaggcgttgtgcgtagaggatcctagaccagcatgccagtgtgccaaggccacagggaaagcgagtggttggtaaaaatccgtgaggtcggcaatatgttgtttttctggaacttacttatggtaaccttttatttattttctaatataatgggggagtttcgtactgaggtgtaaagggatttatatggggacgtaggccgatttccgggtgttgtaggtttctctttttcaggcttatactcatgaatcttgtctgaagcttttgagggcagactgccaagtcctggagaaatagtagatggcaagtttgtgggtttttttttttttacacgaatttgaggaaaaccaaatgaatttgatagccaaattgagacaatttcagcaaatctgtaagcagtttgtatgtttagttggggtaatgaagtatttcagttttgtgaatagatgacctgtttttacttcctcaccctgaattcgttttgtaaatgtagagtttggatgtgtaactgaggcgggggggagttttcagtatttttttttgtgggggtgggggcaaaatatgttttcagttctttttcccttaggtctgtctagaatcctaaaggcaaatgactcaaggtgtaacagaaaacaagaaaatccaatatcaggataatcagaccaccacaggtttacagtttatagaaactagagcagttctcacgttgaggtctgtggaagagatgtccattggagaaatggctggtagttactcttttttccccccacccccttaatcagactttaaaagtgcttaaccccttaaacttgttattttttacttgaagcattttgggatggtcttaacagggaagagagagggtgggggagaaaatgtttttttctaagattttccacagatgctatagtactattgacaaactgggttagagaaggagtgtaccgctgtgctgttggcacgaacaccttcagggactggagctgcttttatccttggaagagtattcccagttgaagctgaaaagtacagcacagtgcagctttggttcatattcagtcatctcaggagaacttcagaagagcttgagtaggccaaatgttgaagttaagttttccaataatgtgacttcttaaaagttttattaaaggggaggggcaaatattggcaattagttggcagtggcctgttacggttgggattggtggggtgggtttaggtaattgtttagtttatgattgcagataaactcatgccagagaacttaaagtcttagaatggaaaaagtaaaggaatatcaacttccaagttggcaagtaactcccaatgatttagtttttttccccccagtttgaattgggaagctgggggaagttaaatatgagccactgggtgtaccagtgcattaatttgggcaaggaaagtgtcataatttgatactgtatctgttttccttcaaagtatagagcttttggggaaggaaagtattgaactgggggttggtctggcctactgggctgacattaactacaattatgggaaatgcaaaagttgtttggatatggtagtgtgtggttctcttttggaatttttttcaggtgatttaataataatttaaaactactatagaaactgcagagcaaaggaagtggcttaatgatcctgaagggatttcttctgatggtagcttttgtattatcaagtaagattctattttcagttgtgtgtatgcaagtttttttttagtgtaggagaaatacttttccattgtttaactgcaaaacaagatgttaaggtatgcttcaaaaattttgtaaattgtttattttaaacttatctgtttgtaaattgtaactgattaagaattgtgatagttcagcttgaatgtctcttagagggtgggcttttgttgatgagggaggggaaactttttttttttctatagacttttttcagataacatcttctgagtcataaccagcctggcagtatgatggcctagatgcagagaaaacagctccttggtgaattgataagtaaaggcagaaaagattatatgtcatacctccattggggaataagcataaccctgagattcttactactgatgagaacattatctgcatatgccaaaaaattttaagcaaatgaaagctaccaatttaaagttacggaatctaccattttaaagttaattgcttgtcaagctataaccacaaaaataatgaattgatgagaaatacaatgaagaggcaatgtccatctcaaaatactgcttttacaaaagcagaataaaagcgaaaagaaatgaaaatgttacactacattaatcctggaataaaagaagccgaaataaatgagagatgagttgggatcaagtggattgaggaggctgtgctgtgtgccaatgtttcgtttgcctcagacaggtatctcttcgttatcagaagagttgcttcatttcatctgggagcagaaaacagcaggcagctgttaacagataagtttaacttgcatctgcagtattgcatgttagggataagtgcttatttttaagagctgtggagttcttaaatatcaaccatggcactttctcctgaccccttccctaggggatttcaggattgagaaatttttccatcgagcctttttaaaattgtaggacttgttcctgtgggcttcagtgatgggatagtacacttcactcagaggcatttgcatctttaaataatttcttaaaagcctctaaagtgatcagtgccttgatgccaactaaggaaatttgtttagcattgaatctctgaaggctctatgaaaggaatagcatgatgtgctgttagaatcagatgttactgctaaaatttacatgttgtgatgtaaattgtgtagaaaaccattaaatcattcaaaataataaactatttttattagagaatgtatacttttagaaagctgtctccttatttaaataaaatagtgtttgtctgtagttcagtgttggggcaatcttgggggggattcttctctaatctttcagaaactttgtctgc gaacactctttaatggaccagatcaggatttgagcggaagaacgaatgtaactttaaggcaggaaagacaaattttattcttcataaagtgatgagcatataataattccaggcacatggcaatagaggccctctaaataaggaataaataacctcttagacaggtgggagattatgatcagagtaaaaggtaattacacattttatttccagaaagtcaggggtctataaattgacagtgattagagtaatactttttcacatttccaaagtttgcatgttaactttaaatgcttacaatcttagagtggtaggcaatgttttacactattgaccttatatagggaagggagggggtgcctgtggggttttaaagaattttcctttgcagaggcatttcatccttcatgaagccattcaggattttgaattgcatatgagtgcttggctcttccttctgttctagtgagtgtatgagaccttgcagtgagtttatcagcatactcaaaatttttttcctggaatttggagggatgggaggagggggtggggcttacttgttgtagctttttttttttacagacttcacagagaatgcagttgtcttgacttcaggtttgtctgttctgttggcaagtaaatgcagtactgttctgatcccgctgctattagaatgcattgtgaaacgactggagtatgattaaaagttgtgttccccaatgcttggagtagtgattgttgaaggaaaaaatccagctgagtgataaaggctgagtgttgaggaaatttctgcagttttaagcagtcgtatttgtgattgaagctgagtacattttgctggtgtatttttaggtaaaatgctttttgttcatttctggtggtgggaggggactgaagcctttagtcttttccagatgcaaccttaaaatcagtgacaagaaacattccaaacaagcaacagtcttcaagaaattaaactggcaagtggaaatgtttaaacagttcagtgatctttagtgcattgtttatgtgtgggtttctctctcccctcccttggtcttaattcttacatgcaggaacactcagcagacacacgtatgcgaagggccagagaagccagacccagtaagaaaaaatagcctatttactttaaataaaccaaacattccattttaaatgtggggattgggaaccactagttctttcagatggtattcttcagactatagaaggagcttccagttgaattcaccagtggacaaaatgaggaaaacaggtgaacaagctttttctgtatttacatacaaagtcagatcagttatgggacaatagtattgaatagatttcagctttatgctggagtaactggcatgtgagcaaactgtgttggcgtgggggtggaggggtgaggtgggcgctaagcctttttttaagatttttcaggtacccctcactaaaggcaccgaaggcttaaagtaggacaaccatggagccttcctgtggcaggagagacaacaaagcgctattatcctaaggtcaagagaagtgtcagcctcacctgatttttattagtaatgaggacttgcctcaactccctctttctggagtgaagcatccgaaggaatgcttgaagtacccctgggcttctcttaacatttaagcaagctgtttttatagcagctcttaataataaagcccaaatctcaagcggtgcttgaaggggagggaaagggggaaagcgggcagccacttttccctagcttttccagaagcctgttaaaagcaaggtctccccacaagcaacttctctgccacatcgccaccccgtgccttttgatctagcacagacccttcacccctcacctcgatgcagccagtagcttggatccttgtgggcatgatccataatcggtttcaaggtaacgatggtgtcgaggtctttggtgggttgaactatgttagaaaaggccattaatttgcctgcaaattgttaacagaagggtattaaaaccacagctaagtagctctattataatacttatccagtgactaaaaccaacttaaaccagtaagtggagaaataacatgttcaagaactgtaatgctgggtgggaacatgtaacttgtagactggagaagataggcatttgagtggctgagagggcttttgggtgggaatgcaaaaattctctgctaagactttttcaggtgaacataacagacttggccaagctagcatcttagcggaagctgatctccaatgctcttcagtagggtcatgaaggtttttcttttcctgagaccatgacggccctgaaaaagggccagagtctcaggttttgctttttggcctttttctagcttaaaaaaaaaaaaagcaaaa

#### MALAT1–mC-4-B^mut^

Gtaaaggactggggccccgcaactggcctctcctgccctcttaagcgcagcgccattttagcaacgcagaagcccggcgccgggaagcctcagctcgcctgaaggcaggtcccctctgacgcctccgggagcccaggtttcccagagtccttgggacgcagcgacgagttgtgctgctatcttagctgtccttataggctggccattccaggtggtggtatttagataaaaccactcaaactctgcagtttggtcttggggtttggaggaaagcttttatttttcttcctgctccggttcagaaggtctgaagctcatacctaaccgggcataacacagaatctgcaaaacaaaaacccctaaaaaagcagacccagagcagtgtaaacacttctgggtgtgtccctgactggctgcccaaggtctctgtgtcttcggagacaaagccattcgcttagttggtctactttaaaaggccacttgaactcgctttccatggcgatttgccttgtgagcactttcaggagagcctggaagctgaaaaacggtagaaaaatttccgtgcgggccgtggggggctggcggcaactggggggccgcagatcagagtgggccactggcagccaacggcccccggggctcaggcggggagcagctctgtggtgtgggattgaggcgttttccaagagtgggttttcacgtttctaagatttcccaagcagacagcccgtgctgctccgatttctcgaacaaaaaagcaaaacgtgtggctgtcttgggagcaagtcgcaggactgcaagcagttgggggagaaagtccgccattttgccacttctcaaccgtccctgcaaggctggggctcagttgcgtaatggaaagtaaagccctgaactatcacactttaatcttccttcaaaaggtggtaaactatacctactgtccctcaagagaacacaagtgctttaagaggtattttaaaagttccgggggttttgtgaggtgtttgatgacccgtttaaaatatgatttccatgtttcttttgtctaaagtttgcagctcaaatctttccacacgctagtaatttaagtatttctgcatgtgtagtttgcattcaagttccataagctgttaagaaaaatctagaaaagtaaaactagaacctatttttaaccgaagaactactttttgcctccctcacaaaggcggcggaaggtgatcgaattccggtgatgcgagttgttctccgtctataaatacgcctcgcccgagctgtgcggtaggcattgaggcagccagcgcaggggcttctgctgagggggcaggcggagcttgaggaaaccgcagatgagtttttttctctttgaaagatagagattaatacaactacttaaaaaatatagtcaataggttactaagatattgcttagcgttaagtttttaacgtaattttaatagcttaagattttaagagaaaatatgaagacttagaagagtagcatgaggaaggaaaagataaaaggtttctaaaacatgacggaggttgagatgaagcttcttcatggagtaaaaaatgtatttaaaagaaaattgagagaaaggactacagagccccgaattaataccaatagaagggcaatgcttttagattaaaatgaaggtgacttaaacagcttaaagtttagtttaaaagttgtaggtgattaaaataatttgaaggcgatcttttaaaaagagattaaactgaaggtgattaaaagaccttgaaatccatgacgcagggagaattgcgtcatttaaagcctagttaacgcatttactaaacgcagacgaaaatggaaagattaattgggagtggtaggatgaaacaatttggagaag atagaagtttgaagtggaaaactggaagacagaagtacgggaaggcgaagaaaagaatagagaagatagggaaattagaagataaaaacatacttttagaagaaaaaagataaatttaaacctgaaaagtaggaagcagaagaaaaaagacaagctaggaaacaaaaagctaagggcaaaatgtacaaacttagaagaaaattggaagatagaaacaagatagaaaatgaaaatattgtcaagagtttcagatagaaaatgaaaaacaagctaagacaagtattggagaagtatagaagatagaaaaatataaagccaaaaattggataaaatagcactgaaaaaatgaggaaattattggtaaccaatttattttaaaagcccatcaatttaatttctggtggtgcagaagttagaaggtaaagcttgagaagatgagggtgtttacgtagaccagaaccaatttagaagaatacttgaagctagaaggggaagttggttaaaaatcacatcaaaaagctactaaaaggactggtgtaatttaaaaaaaactaaggcagaaggcttttggaagagttagaagaatttggaaggccttaaatatagtagcttagtttgaaaaatgtgaaggactttcgtaacggaagtaattcaagatcaagagtaattaccaacttaatgtttttgcattggactttgagttaagattattttttaaatcctgaggactagcattaattgacagctgacccaggtgctacacagaagtggattcagtgagtctaggaagacagcagcagacaggattccaggaaccagtgtttgatgaagctaggactgaggagcaagcgagcaagcagcagttcgtggtgaagataggaaaagagtccaggagccagtgcgatttggtgaaggaagctaggaagaaggaaggagcgctaacgatttggtggtgaagctaggaaaaaggattccaggaaggagcgagtgcaatttggtgatgaaggtagcaggcggcttggcttggcaaccacacggaggaggcgagcaggcgttgtgcgtagaggatcctagaccagcatgccagtgtgccaaggccacagggaaagcgagtggttggtaaaaatccgtgaggtcggcaatatgttgtttttctggaacttacttatggtaaccttttatttattttctaatataatgggggagtttcgtactgaggtgtaaagggatttatatggggacgtaggccgatttccgggtgttgtaggtttctctttttcaggcttatactcatgaatcttgtctgaagcttttgagggcagactgccaagtcctggagaaatagtagatggcaagtttgtgggtttttttttttttacacgaatttgaggaaaaccaaatgaatttgatagccaaattgagacaatttcagcaaatctgtaagcagtttgtatgtttagttggggtaatgaagtatttcagttttgtgaatagatgacctgtttttacttcctcaccctgaattcgttttgtaaatgtagagtttggatgtgtaactgaggcgggggggagttttcagtatttttttttgtgggggtgggggcaaaatatgttttcagttctttttcccttaggtctgtctagaatcctaaaggcaaatgactcaaggtgtaacagaaaacaagaaaatccaatatcaggataatcagaccaccacaggtttacagtttatagaaactagagcagttctcacgttgaggtctgtggaagagatgtccattggagaaatggctggtagttactcttttttccccccacccccttaatcagactttaaaagtgcttaaccccttaaacttgttattttttacttgaagcattttgggatggtcttaacagggaagagagagggtgggggagaaaatgtttttttctaagattttccacagatgctatagtactattgacaaactgggttagagaaggagtgtaccgctgtgctgttggcacgaacaccttcagggactggagctgcttttatccttggaagagtattcccagttgaagctgaaaagtacagcacagtgcagctttggttcatattcagtcatctcaggagaacttcagaagagcttgagtaggccaaatgttgaagttaagttttccaataatgtgacttcttaaaagttttattaaaggggaggggcaaatattggcaattagttggcagtggcctgttacggttgggattggtggggtgggtttaggtaattgtttagtttatgattgcagataaactcatgccagagaacttaaagtcttagaatggaaaaagtaaaggaatatcaacttccaagttggcaagtaactcccaatgatttagtttttttccccccagtttgaattgggaagctgggggaagttaaatatgagccactgggtgtaccagtgcattaatttgggcaaggaaagtgtcataatttgatactgtatctgttttccttcaaagtatagagcttttggggaaggaaagtattgaactgggggttggtctggcctactgggctgacattaactacaattatgggaaatgcaaaagttgtttggatatggtagtgtgtggttctcttttggaatttttttcaggtgatttaataataatttaaaactactatagaaactgcagagcaaaggaagtggcttaatgatcctgaagggatttcttctgatggtagcttttgtattatcaagtaagattctattttcagttgtgtgtatgcaagtttttttttagtgtaggagaaatacttttccattgtttaactgcaaaacaagatgttaaggtatgcttcaaaaattttgtaaattgtttattttaaacttatctgtttgtaaattgtaactgattaagaattgtgatagttcagcttgaatgtctcttagagggtgggcttttgttgatgagggaggggaaactttttttttttctatagacttttttcagataacatcttctgagtcataaccagcctggcagtatgatggcctagatgcagagaaaacagctccttggtgaattgataagtaaaggcagaaaagattatatgtcatacctccattggggaataagcataaccctgagattcttactactgatgagaacattatctgcatatgccaaaaaattttaagcaaatgaaagctaccaatttaaagttacggaatctaccattttaaagttaattgcttgtcaagctataaccacaaaaataatgaattgatgagaaatacaatgaagaggcaatgtccatctcaaaatactgcttttacaaaagcagaataaaagcgaaaagaaatgaaaatgttacactacattaatcctggaataaaagaagccgaaataaatgagagatgagttgggatcaagtggattgaggaggctgtgctgtgtgccaatgtttcgtttgcctcagacaggtatctcttcgttatcagaagagttgcttcatttcatctgggagcagaaaacagcaggcagctgttaacagataagtttaacttgcatctgcagtattgcatgttagggataagtgcttatttttaagagctgtggagttcttaaatatcaaccatggcactttctcctgaccccttccctaggggatttcaggattgagaaatttttccatcgagcctttttaaaattgtaggacttgttcctgtgggcttcagtgatgggatagtacacttcactcagaggcatttgcatctttaaataatttcttaaaagcctctaaagtgatcagtgccttgatgccaactaaggaaatttgtttagcattgaatctctgaaggctctatgaaaggaatagcatgatgtgctgttagaatcagatgttactgctaaaatttacatgttgtgatgtaaattgtgtagaaaaccattaaatcattcaaaataataaactatttttattagagaatgtatacttttagaaagctgtctccttatttaaataaaatagtgtttgtctgtagttcagtgttggggcaatcttgggggggattcttctctaatctttcagaaactttgtctgcgaacactctttaatggaccagatcaggatttgagcggaagaacgaatgtaactttaaggcaggaaagacaaattttattcttcataaagtgatgagcatataataattccaggcacatggcaatagaggccctctaaataaggaataaataacctcttagacaggtgggagattatgatcagagtaaaaggtaattacacattttatttccagaaagtcaggggtctataaattgacagtgattagagtaatactttttcacatttccaaagtttgcatgttaactttaaatgcttacaatcttagagtggtaggcaatgttttacactattgaccttatatagggaagggagggggtgcctgtggggttttaaagaattttcctttgcagaggcatttcatccttcatgaagccattcaggattttgaattgcatatgagtgcttggctcttccttctgttctagtgagtgtatgagaccttgcagtgagtttatcagcatactcaaaatttttttcctggaatttggagggatgggaggagggggtgggg cttacttgttgtagctttttttttttacagacttcacagagaatgcagttgtcttgacttcaggtttgtctgttctgttggcaagtaaatgcagtactgttctgatcccgctgctattagaatgcattgtgaaacgactggagtatgattaaaagttgtgttccccaatgcttggagtagtgattgttgaaggaaaaaatccagctgagtgataaaggctgagtgttgaggaaatttctgcagttttaagcagtcgtatttgtgattgaagctgagtacattttgctggtgtatttttaggtaaaatgctttttgttcatttctggtggtgggaggggactgaagcctttagtcttttccagatgcaaccttaaaatcagtgacaagaaacattccaaacaagcaacagtcttcaagaaattaaactggcaagtggaaatgtttaaacagttcagtgatctttagtgcattgtttatgtgtgggtttctctctcccctcccttggtcttaattcttacatgcaggaacactcagcagacacacgtatgcgaagggccagagaagccagacccagtaagaaaaaatagcctatttactttaaataaaccaaacattccattttaaatgtggggattgggaaccactagttctttcagatggtattcttcagactatagaaggagcttccagttgaattcaccagtggacaaaatgaggaaaacaggtgaacaagctttttctgtatttacatacaaagtcagatcagttatgggacaatagtattgaatagatttcagctttatgctggagtaactggcatgtgagcaaactgtgttggcgtgggggtggaggggtgaggtgggcgctaagcctttttttaagatttttcaggtacccctcactaaaggcaccgaaggcttaaagtaggacaaccatggagccttcctgtggcaggagagacaacaaagcgctattatcctaaggtcaagagaagtgtcagcctcacctgatttttattagtaatgaggacttgcctcaactccctctttctggagtgaagcatccgaaggaatgcttgaagtacccctgggcttctcttaacatttaagcaagctgtttttatagcagctcttaataataaagcccaaatctcaagcggtgcttgaaggggagggaaagggggaaagcgggcagccacttttccctagcttttccagaagcctgttaaaagcaaggtctccccacaagcaacttctctgccacatcgccaccccgtgccttttgatctagcacagacccttcacccctcacctcgatgcagccagtagcttggatccttgtgggcatgatccataatcggtttcaaggtaacgatggtgtcgaggtctttggtgggttgaactatgttagaaaaggccattaatttgcctgcaaattgttaacagaagggtattaaaaccacagctaagtagctctattataatacttatccagtgactaaaaccaacttaaaccagtaagtggagaaataacatgttcaagaactgtaatgctgggtgggaacatgtaacttgtagactggagaagataggcatttgagtggctgagagggcttttgggtgggaatgcaaaaattctctgctaagactttttcaggtgaacataacagacttggccaagctagcatcttagcggaagctgatctccaatgctcttcagtagggtcatgaaggtttttcttttcctgagaccatgacggccctaaaaaagggccagagtctcaggttttgctttttggcctttttctagcttaaaaaaaaaaaaagcaaaa

##### minimal CMV (miniCMV) promoter

ggtaggcgtgtacggtgggaggtctatataagcagagct

(SphI postoctamer homology–octamer–proximal sequence element–TATA box)

##### minimal U6 (miniU6) promoter

gagggcctatttcccatgattccttcatatttgcatatacgatagcttaccgtaacttgaaagtatttcgatttcttggctttatatatcttgtggaaaggacgaaacacc

##### minimal U6 promoter–mC-4-B–drz-Mtgn-3–polyT

gagggcctatttcccatgattccttcatatttgcatatacgatagcttaccgtaacttgaaagtatttcgatttcttggctttatatatcttgtggaaaggacgaaacaccgggttccatgacggccctgaaaaagggccagagaacccgagggatagtagacagagcctccccgtggctcccttggataaccaactgatactgtacttttttt

##### minimal U6 promoter–mC-4-B^dual^ ^mut^–drz-Mtgn-3–polyT

gagggcctatttcccatgattccttcatatttgcatatacgatagcttaccgtaacttgaaagtatttcgatttcttggctttatatatcttgtggaaaggacgaaacaccgggttccattccggcccttacatagggccaacgaacccgagggatagtagacagagcctccccgtggctcccttggataaccaactgatactgtacttttttt

##### minimal U6 promoter (inP3)

gagggcctatttcccatgattccttcatatttgcatatacgatagcttaccgtaacttgaaagtatttcgatttcttgGTAgctttatatatcttgtggaaaggacgaaacacc

##### minimal U6 promoter (inP5)

gagggcctatttcccatgattccttcatatttgcatatacgatagcttaccgtaacttgaaagtatttcgatttcttgGTACCgctttatatatc ttgtggaaaggacgaaacacc

